# Topology of molecular networks offers signaling insusceptibility to temperature and ionic strength changes

**DOI:** 10.1101/2025.10.09.680621

**Authors:** Egor S. Korenkov, Maxim P. Nikitin

## Abstract

The equilibrium constants of chemical reactions fundamentally depend on temperature, posing challenges for living systems. However, many conformer organisms do not maintain stable internal temperature. This raises the question: can molecular signaling pathways inherently resist temperature susceptibility? Molecular commutation is a recently discovered, fundamentally distinct mechanism for biological information processing and storage that underlies highly complex signal processing and computation using only molecular interactions governed by the law of mass action. Here we show that molecular commutation enables biological information processing networks to become temperature independent through compensatory reactions (i.e., network topology). Using complex logic gates, receptor-activator networks, and signaling systems with non-linear, non-monotonic functions (e.g., x^2^, x^3^), we show that topological compensation preserves signaling function across temperatures up to 0.1–50 °C, despite dissociation constant changes of up to nine orders of magnitude. This mechanism also stabilizes systems against dramatic ionic strength shifts (e.g., Na^+^ from 0.1–1 M). Thus, topological compensation is a unique homeostasis mechanism that may be used by delicate biological systems of arbitrarily high complexity.

**Graphical abstract:** 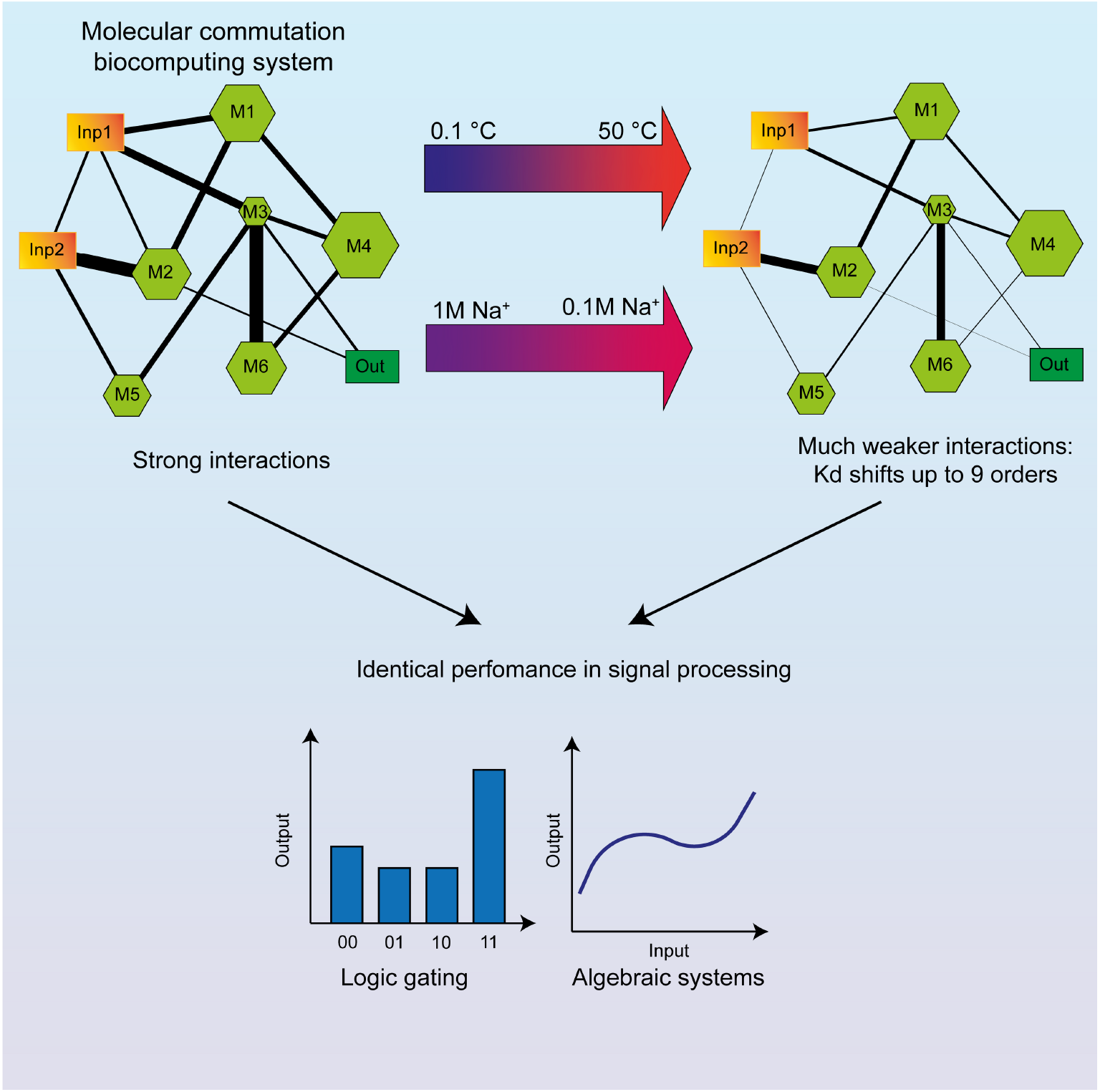

## Introduction

The Boltzmann distribution fundamentally dictates that equilibrium parameters of any chemical reaction, such as equilibrium constants, follow an exponential dependence on temperature of the form e^E/kT^. This principle plays a critical role in biological systems, as even small variations in temperature can lead to significant changes in the parameters of biochemical reactions, potentially resulting in severe physiological consequences at the organismal level. It is therefore unsurprising that many species have evolved complex homeostatic mechanisms to maintain a stable internal temperature.

However, a broad range of organisms do not exhibit temperature homeostasis. For instance, numerous pathogenic bacteria^1^ experience temperature shifts of approximately 30 °C during their transition from environmental reservoirs, such as soil, to the human host. Similarly, many eurythermal fish species—such as *Cyprinus carpio*^2^ and *Fundulus heteroclitus*^3^ —are capable of thriving in environments ranging from near-freezing (~3 °C) to warm summer waters (~35 °C). Moreover, even humans can tolerate prolonged exposure (hours to days) of extremities, to near-freezing temperatures without significant harm, provided moisture is absent^4^.

A question arises as to how, in all these cases, the complex regulatory and signaling networks of biochemical reactions, which form basics of any living system, continue to properly function, given that the parameters of all the constituent reactions vary dramatically.

In general, signal transduction and processing using molecular systems have long been the subject of active investigation^5,6^. In fact, to date, a large number of biocomputing approaches, i.e., artificial molecular systems that can process data and various signals, have been developed based on enzymatic networks^7^, nucleic acids^8^, nanoscale structures^9,10^, among many others. However, most of these systems could hardly have evolved in nature as they require rather specific and unnatural design of participating molecules or nanostructures.

Conversely, numerous studies have identified and characterized concrete examples of computational pathways in living organisms, including interacting protein networks^11^, as well as instances of biocomputation at the level of large receptor systems^12^ or even entire cells^13^. In such systems, one of the key factors is the presence of signal-analyzing and -transforming subsystems. For example, networks of dimerizing transcription factors cannot function in the absence of respective enzymatic systems^11^, while olfactory receptors^12^ require the central nervous system for signal processing.

A recent 2023 study^14,15^ demonstrated the biomolecular information processing principle of “molecular commutation” by experimentally proving that highly complex information processing tasks can be performed by molecular ensembles solely through their reversible low-affinity all-to-all molecular association/dissociation reactions. Moreover, the study experimentally showed that the outcome of such information processing can have a significant direct impact on the key biological process, such as translation.

Molecular commutation” (which is sometimes referred to as “competitive dimerization network computations”) has a notable distinction from other systems involving promiscuous (many-to-many) interactions^16^. In the molecular commutation regime, the entirety of these complex computations happens according to law of mass action via low-affinity interactions of molecules (i.e., when dissociation constants are around concentration levels of participating molecules), and doesn’t require any other supplementary complex system for the computation to occur. In molecular commutation paradigm, any molecule—regardless of specific properties—can act as a signal integrator, i.e., without the need for any higher-level signal-integrating systems such as the abovementioned enzymes or brain cell networks. Currently, this computational principle is rapidly developing: theoretical studies have explored the properties of such computations^17^, while both theoretical and experimental work has demonstrated increasingly complex computations, including, for example, the construction of classifying neural networks^18–20^.

In this work, we report a fundamentally novel property of molecular commutation networks for biological systems: these networks can overcome the inherent Boltzmann temperature dependence, preserving their computational functionality across a wide temperature range with arbitrarily high accuracy. We demonstrate a feature that we term “topological compensation”: to counteract a change in reactions of interest, one can expand the topology of the molecular commutation network with compensatory reactants and properly adjust their affinity parameters toward other members of the network. As molecular commutation signal processing relies solely on low-affinity interactions between the molecules, such compensatory reactants are always abundantly available in virtually any complex biomolecular system, such as cells. As we show below, such “topological compensation” enables the temperature-independent realization of logical gates (such as YES, NOT, AND), complex analog signals of arbitrary algebraic functions (e.g., x^2^, x^3^), and even large-scale receptor-activator networks.

We show that such networks can be rationally designed by introducing a small number of compensatory reactions into existing computational circuits, or they can emerge evolutionarily through stochastic optimization to function correctly across a predefined temperature range. Importantly, topological compensation alone is sufficient to achieve temperature invariance even in large-scale, multi-component systems comprising hundreds of molecular species, and in networks with arbitrarily complex computational architectures.

## Results

### Concept

As we noted previously^14,15^, information processing systems based on molecular commutation can be implemented with any type of molecule-proteins, DNA, carbohydrates, small molecules, etc. However, DNA is currently the most convenient model for creating and testing molecular circuitry as the affinity constants between oligonucleotides can be easily estimated from their sequences (which is incomparably harder for proteins at the moment^21^).

First, we will focus on creating biocomputing systems that function similarly at temperatures of 25 and 50 °C. All oligonucleotide duplexes exhibit significantly lower stability at 50 °C compared to 25 °C. **Figure 1A** shows the distribution of dissociation constants (Kd) at these two temperatures for a dataset of 10,000 randomly generated oligonucleotide pairs (length 20 nucleotides) containing 1 to 10 mismatches, as predicted by NUPACK^22,23^. On average, Kd increases by several orders of magnitude, with the distribution spanning roughly two orders of magnitude. For instance, if Kd values at 25 °C are constrained to the range of 10 to 100 picomolar (**Figure 1B**), the corresponding values at 50 °C typically fall within the range of 0.1 to 10 micromolar. If we consider duplexes that exhibit similar dissociation constants at 25 °C, and among these, we refer to those that maintain stronger binding at 50 °C (i.e., have lower Kd values) as “hard-to-melt,” while those with weaker binding at 50 °C are termed “easy-to-melt.”

**Figure 1.**
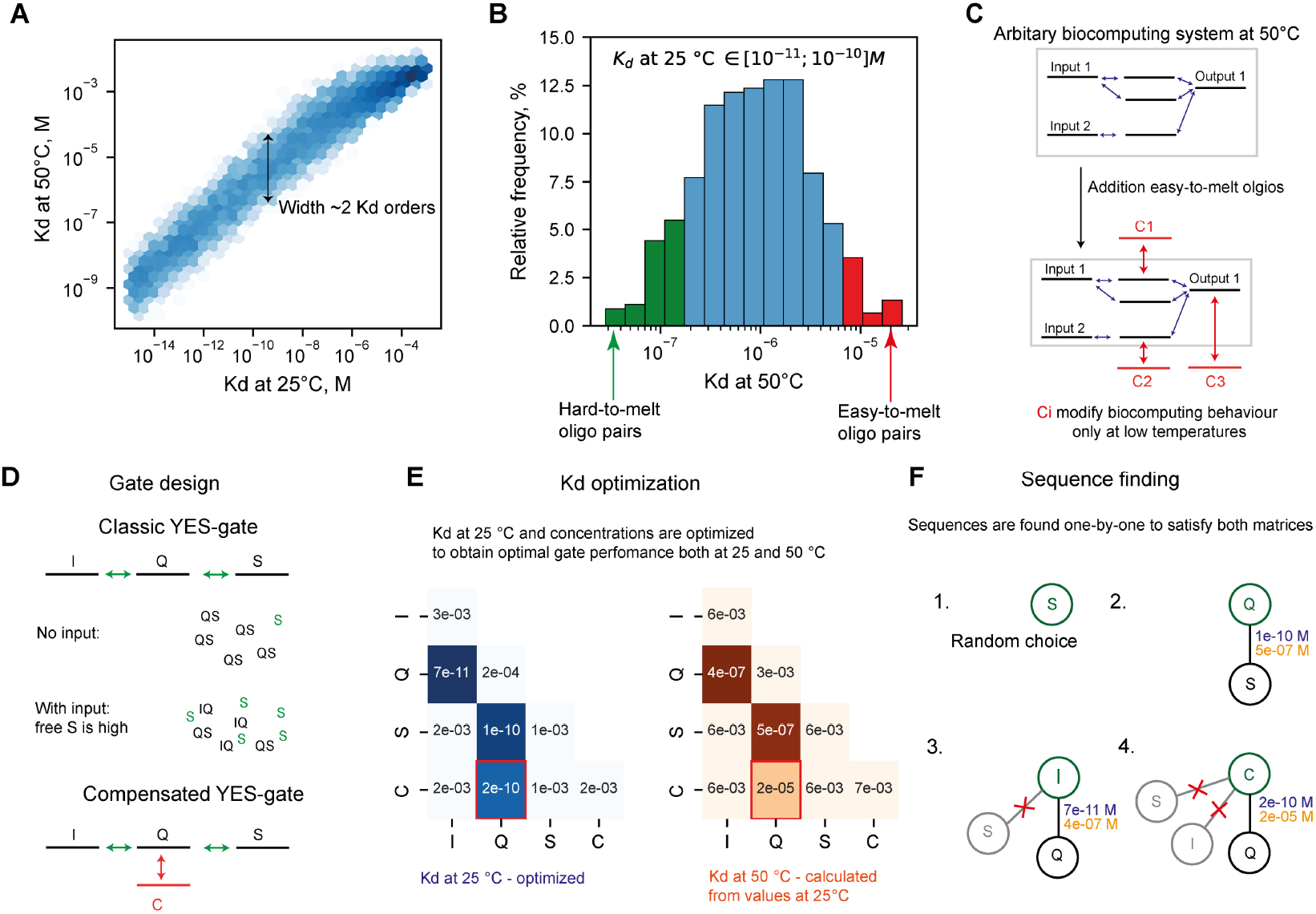
Demonstration of the topological compensation concept. (A) Distribution of dissociation constants (Kd) at 25 °C and 50 °C for 20-nucleotide long oligonucleotides with up to 10 mismatches, as predicted by NUPACK. (B) Distribution of Kd values at 50 °C for oligonucleotide pairs that exhibit Kd values between 10 and 100 pM at 25 °C. (C) General design of a temperature-independent biocomputing system. (D) Design of a conventional YES gate and its temperature-independent counterpart. (E) Optimization of binding parameters for YES gate components. (F) Overview of the sequence selection algorithm.

Let’s consider a generic biocomputing system (**Figure 1C**), that operates optimally at 50 °C but exhibits altered behavior at lower temperatures. We can introduce new participants to this system, that are extremely “easy-to-melt”, but still have considerably strong binding at 25 °C. The oligonucleotides primarily influence system behavior at lower temperatures, while exerting minimal effects at elevated temperatures. Consequently, if the system is optimized to function effectively at low temperature, its performance can become effectively temperature-independent, despite the fact that individual reaction components are themselves sensitive to temperature changes.

As a specific example of this concept, we turn to the molecular YES gate, following the original work on molecular commutation^14^ (**Figure 1D**). This system consists of three oligonucleotides, I (input), Q (quencher) and S (signal). The logical input is defined as 0 when I is absent and 1 when I is present at a fixed concentration. The oligonucleotide S serves as the output. The logical output is considered 0 when the concentration of single-stranded S is below a defined threshold, and 1 when it exceeds that threshold.

Now, when I is absent, Q binds to S, leaving almost no free S. When I is present, it competes for Q, occupying it, leaving much more free S. In these terms, the system behaves like a Boolean function YES: logic output of this system is always the same as the input.

This YES-gate operates at 50 °C and exhibits two distinct concentrations of free S, depending on whether the input is absent or present. When the temperature is lowered, both values tend to decrease due to enhanced binding affinities. To counteract this effect, we introduce a compensatory, “easy-to-melt” oligonucleotide C, which binds to Q. This oligo increases the concentration of free S under both input conditions — but only at 25 °C. Achieving identical free S concentrations at both temperatures, however, requires careful optimization.

We divide this optimization process into two stages. First, we determine the dissociation constants and initial concentrations necessary for the system to function as desired (**Figure 1E**). At this stage, we ignore the specific oligonucleotide sequences and instead work with abstract parameters. Every dimerization network can be fully described by a matrix of dissociation constants (Kd) and the initial component concentrations. Given these parameters, the system of equilibrium equations can be solved numerically to compute the resulting free S concentration and simulate system behavior. This leads to a purely numerical continuous optimization problem—in this case, involving the optimization of four dissociation constants and four initial concentrations to match target free S levels at both temperatures. The optimization is performed using the Bayesian adaptive direct search^24^, via the PyBADS software^25^. PyBADS alternates between a series of fast, local Bayesian optimization steps and a systematic, slower exploration of a mesh grid.

However, it is not possible to vary the dissociation constants (Kd) at 25 °C and 50 °C independently, as the thermodynamic properties of oligonucleotide binding impose a specific relationship between them. To account for this constraint, we classify each interaction as either “hard-to-melt” (e.g., S–Q and I–Q interactions) or “easy-to-melt” (e.g., the Q–C interaction). These classifications correspond to characteristic positions within a predefined distribution of Kd values across temperatures (**Figure 1A**). Thus, once the Kd at 25 °C is specified for a given interaction, the Kd at 50 °C can be uniquely determined based on the interaction type. This approach reduces the number of independent parameters in the model and ensures that the simulated system behavior reflects the underlying physical constraints of oligonucleotide thermodynamics. A formal description of the Kd recalculation process for different temperatures and «easy-to-melt» levels is provided in the Materials and Methods section; see also **Figure S1** and **Table S1**.

After identifying the optimal parameters, we proceed to the sequence design stage (**Figure 1F**). We begin by randomly initializing the sequence of the first oligonucleotide, for example, S. Next, we design the sequence of Q, which is intended to interact with S. Q is initially set as the fully complementary sequence to S, and then systematically mutated until the dissociation constants K_d_(SQ) at both temperatures closely match the previously optimized values. After that, we repeat the process for I, so that K_d_ (IQ) is close enough to its the optimized value, and K_d_ (IS) is larger than some off-target threshold. The same procedure is also applied to the remaining oligonucleotides in the system (oligonucleotide C in this particular case), to ensure all interactions satisfy their respective design criteria. Once all oligonucleotide sequences have been determined, a final simulation of the system is performed to verify that the resulting behavior matches the intended design. The performance of the optimized system is also compared with that of the control system, which only lacks the compensatory oligo C. It is observed that in the optimized system, the output signals corresponding to both possible input values are nearly identical at both temperature conditions (**Figure 2A**).

**Figure 2.**
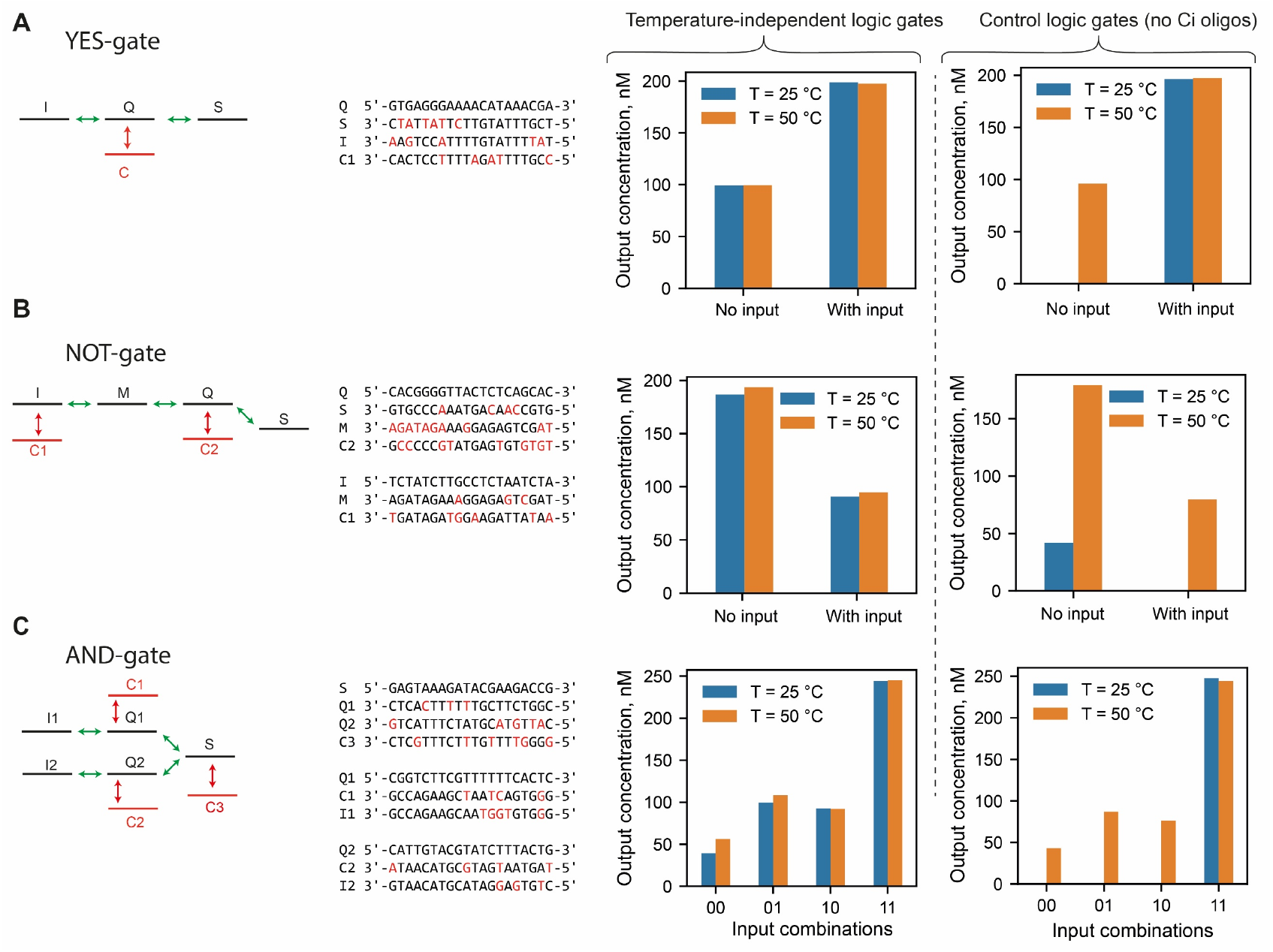
Design and predicted performance of temperature-independent logic gates. (A) YES gate; (B) NOT gate; (C) AND gate. For each gate, the left panel shows the performance of the optimized compensated system, while the right panel shows the performance of the same system without compensatory oligonucleotides.

Using the same design principle, we construct more complex logic gates. The NOT gate is the logical inverse of the YES gate: when the input is 0, the output should be 1, and vice versa. This functionality is implemented using a four-oligonucleotide scheme (**Figure 2B**). When the input I is absent, Q preferentially binds to M, leaving S mostly unbound and therefore in the “on” state. When I is present, Q binds to I, reducing its availability to bind M; the remaining unbound Q then binds to S, resulting in S being predominantly in the bound state, which corresponds to output 0.

In this case, two easy-to-melt oligonucleotides are required for temperature compensation: one that binds to I and another that binds to Q. Using this interaction topology, we apply the same sequence design procedure to determine the sequences of all participating oligonucleotides, ensuring that the desired thermodynamic behavior is maintained across temperatures.

The AND gate is a two-input logic function in which the output is 1 only when both inputs are present. Its topology resembles two parallel YES gates that share a common output oligonucleotide, S (**Figure 2C**). Each of the input-responsive strands, Q1 and Q2, can independently bind to S with sufficient affinity such that the presence of either Q1 or Q2 alone results in S being mostly bound.

To enforce temperature robustness we introduce compensatory, easy-to-melt oligonucleotides for both Q1 and Q2, as in the YES-gate design. Additionally, we include a compensatory oligo targeting S to fine-tune the system’s response and achieve correct behavior at both temperatures. Sequence optimization is then performed for all components following the same procedure.

For all three gates, we also consider control systems—identical to the ones just described but lacking the compensatory oligos Ci. In all three cases, the output concentration in the control systems changes significantly with temperature, in contrast to the optimized systems.

To simplify the design of the described systems, we considered only two input conditions: complete absence or presence at a single fixed concentration. Nevertheless, the final gates maintain the same input–response pattern at both temperatures across all intermediate input concentrations (**Figure S2**).

Importantly, if instead of DNA they networks consist of proteins or other molecules with the same mutual affinities as in the examples above and below, we would expect similar behavior of the respective systems.

### Experimental realization of temperature-independent gates

Now, after we have described the general theory for designing temperature-independent logic gates, we want to demonstrate the real-life feasibility of implementing this idea in a wet lab experiment using the YES gate as an example.

The logic gates described above operate under equilibrium thermodynamics. However, for practical implementation in experimental settings, it is essential that all components reach equilibrium within a reasonable time frame. Long, fully complementary oligonucleotides exemplify extremely stable, non-covalent interactions that behave as effectively irreversible and remain stable almost indefinitely under room temperature conditions^26^.

To identify an upper limit for oligonucleotide affinity that still allows equilibration within several hours, we performed a screening wet lab experiment. We used a classical YES-gate configuration in which the output strand S was labeled with the fluorophore Cyanine 3, and the quencher strand Q was labeled with BHQ2. In this setup, the concentration of free S directly correlates with fluorescence intensity. We tested several Q sequences with varying affinities to S, corresponding to dissociation constants (Kd) ranging from 3.2 nM to 8.6 fM (as predicted by NUPACK). For each Q, we used an input strand I that was identical to S except for having one fewer mismatch with Q. As a result, the affinity of I for Q was expected to be stronger than that of S for Q by design.

We initially mixed Q with S and incubated the mixture for 10 minutes to allow complex formation. Subsequently, we added the input strand I and monitored fluorescence over a 4-hour period. The resulting fluorescence levels, normalized as a percentage of the maximum possible signal (measured after 16 hours of incubation following a 1-hour annealing step at 60 °C), are shown in **Figure 3A**.

**Figure 3.**
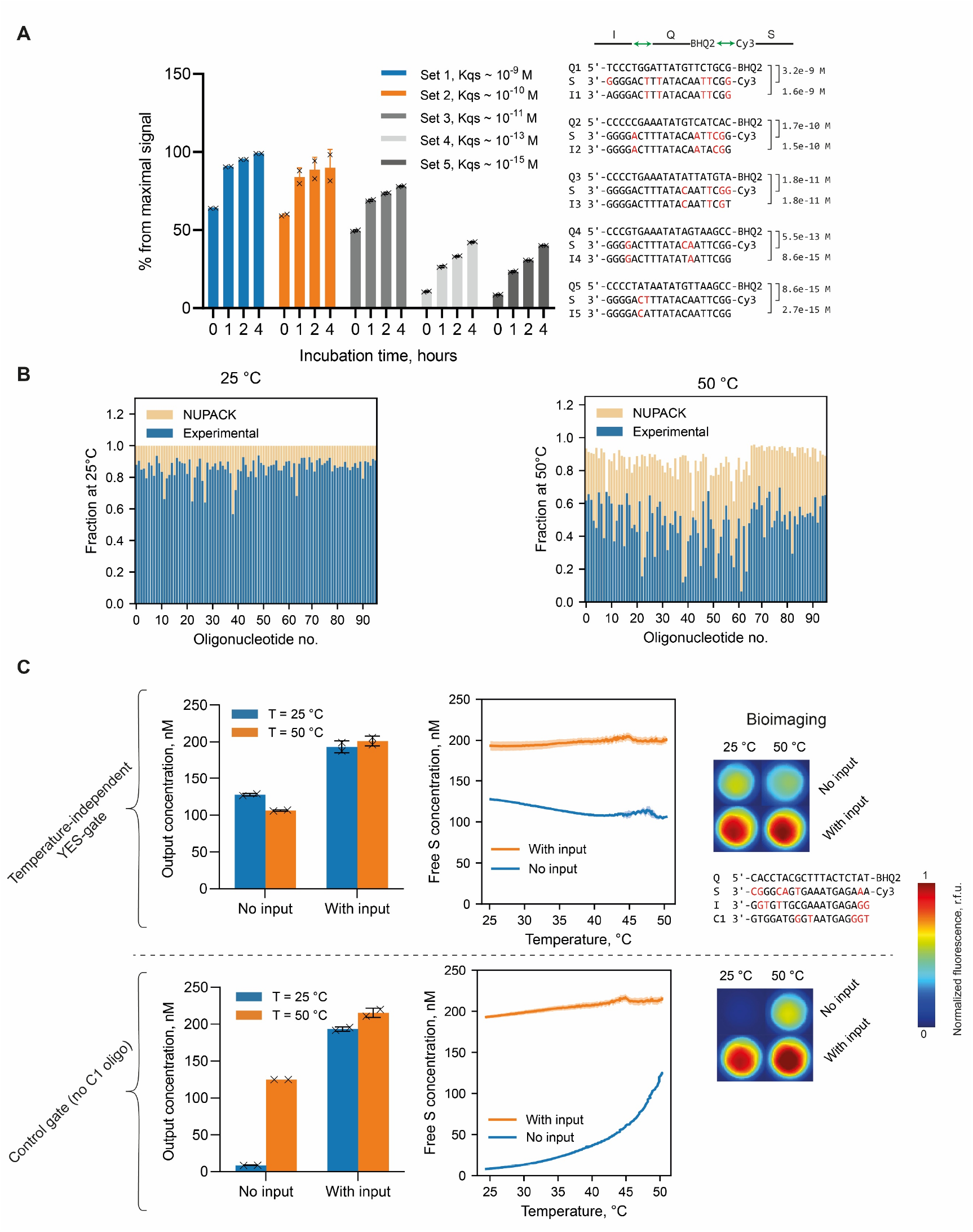
Experimental realization of a temperature-independent YES gate. (A) Displacement kinetics of strand Q by input strand I from the QS complex at different affinity levels. The concentration of each oligonucleotide is 100 nM. (B) Accuracy of NUPACK affinity predictions at 25 °C and 50 °C. Affinity is represented as the concentration of the resulting duplex when both strands are mixed at 1 µM. (C) Experimental performance of the temperature-independent YES gate. Shaded areas indicate 95% confidence intervals (n = 2 replicates). Melting curves are shown alongside fluorescent images of the reaction wells for both the compensated and non-compensated YES gate designs. Due to the temperature dependence of Cyanine3 quantum yield, absolute fluorescence values differ between 25 °C and 50 °C. For clarity, images were linearly adjusted so that control wells containing Cyanine3 exhibit the same mean fluorescence.

The data reveal a clear trend: higher-affinity duplexes equilibrate more slowly. Duplexes with dissociation constants around 1 nM approach equilibrium within approximately 1 hour, while those with Kd values near or below 100 fM fail to reach even 50% of their maximum signal after 4 hours. Based on these findings, we impose a practical lower limit of approximately Kd > 0.01 nM in all sequence optimization procedures (including those described previously) to balance high binding affinity with acceptable equilibration speed. In this experiment all oligonucleotides had a concentration of 100 nM.

To successfully implement the logic gates in a wet lab experimental setting, the oligonucleotides used must exhibit dissociation constants (Kd) that closely match those obtained through prior thermodynamic optimization. However, this condition is not always met in practice, as previous studies have shown that affinity predictions by tools such as NUPACK can be inaccurate, particularly for oligonucleotide duplexes with low complementarity. Moreover, since our experimental setup uses fluorescent labels, they may also contribute to the interaction energy^27^, which is not accounted for by NUPACK.

To verify this limitation and manually identify suitable oligonucleotide sequences for a temperature-independent YES gate, we conducted a systematic screening experiment using the same YES-gate configuration. In this screening, we tested 96 different input strands (I) against a fixed pair of strands, S and Q.

We measured the fluorescence output of each gate to infer the affinity of the I–Q interaction and plotted the results as the equilibrium concentration of the IQ duplex, assuming both strands were initially present at 1 μM (**Figure 3B**). This value ranges from 0 to 1, with values closer to 1 indicating stronger binding. Measurements were performed at both 25 °C and 50 °C. In all cases, the experimentally measured apparent affinities were significantly lower than those predicted by NUPACK. For example, at 25 °C, the predicted affinities were close to 1 for all tested sequences, whereas the measured values ranged from 0.5 to 0.95.

Based on the screening results, we selected suitable candidates for constructing a temperature-independent YES gate. Oligonucleotides that exhibited the strongest binding to Q at both 25 °C and 50 °C were chosen to serve as input strands (I). Candidates for the compensatory strand C were selected from sequences that showed relatively strong binding at 25 °C but significantly weaker binding at 50 °C, consistent with the desired easy-to-melt behavior.

With experimentally validated affinities in hand, we re-ran the Bayesian optimization procedure to determine the optimal concentrations of all components under the new constraints. These optimized concentrations were then used in the final experimental implementation of the gate.

Using the selected oligonucleotide candidates, we measured the performance of the temperature-compensated YES gate across the target temperature range of 25–50 °C and compared it to a non-compensated version of the gate. Melting curves for both systems are presented in **Figure 3C**, along with fluorescence images of the reaction wells at 25 °C and 50 °C.

In the non-compensated gate, the melting curve corresponding to the input = 0 condition shows a substantial increase in fluorescence, reflecting a 15-fold change in the concentration of free S over the temperature range. In contrast, the temperature-compensated gate maintains a stable output, with the variation in free S concentration remaining below 20% across temperatures, regardless of input state. Therefore, the performed wet lab experiments suggest that topological compensation approach indeed can achieve a high degree of temperature independence in biocomputing systems.

### Exploring gates performance at various conditions

Next, we turned again to computational methods to demonstrate the capabilities of topological compensation for more complex conditions and for more sophisticated biocomputational systems.

Here, we constructed a YES gate designed to operate over a significantly broader temperature range—from 0.1 °C to 50 °C. While the previously described design strategy was still effective in generating sequences that performed well at the extremes of this range, we observed substantial shifts in the concentration of free S at intermediate temperatures (**Figure 4A**).

**Figure 4.**
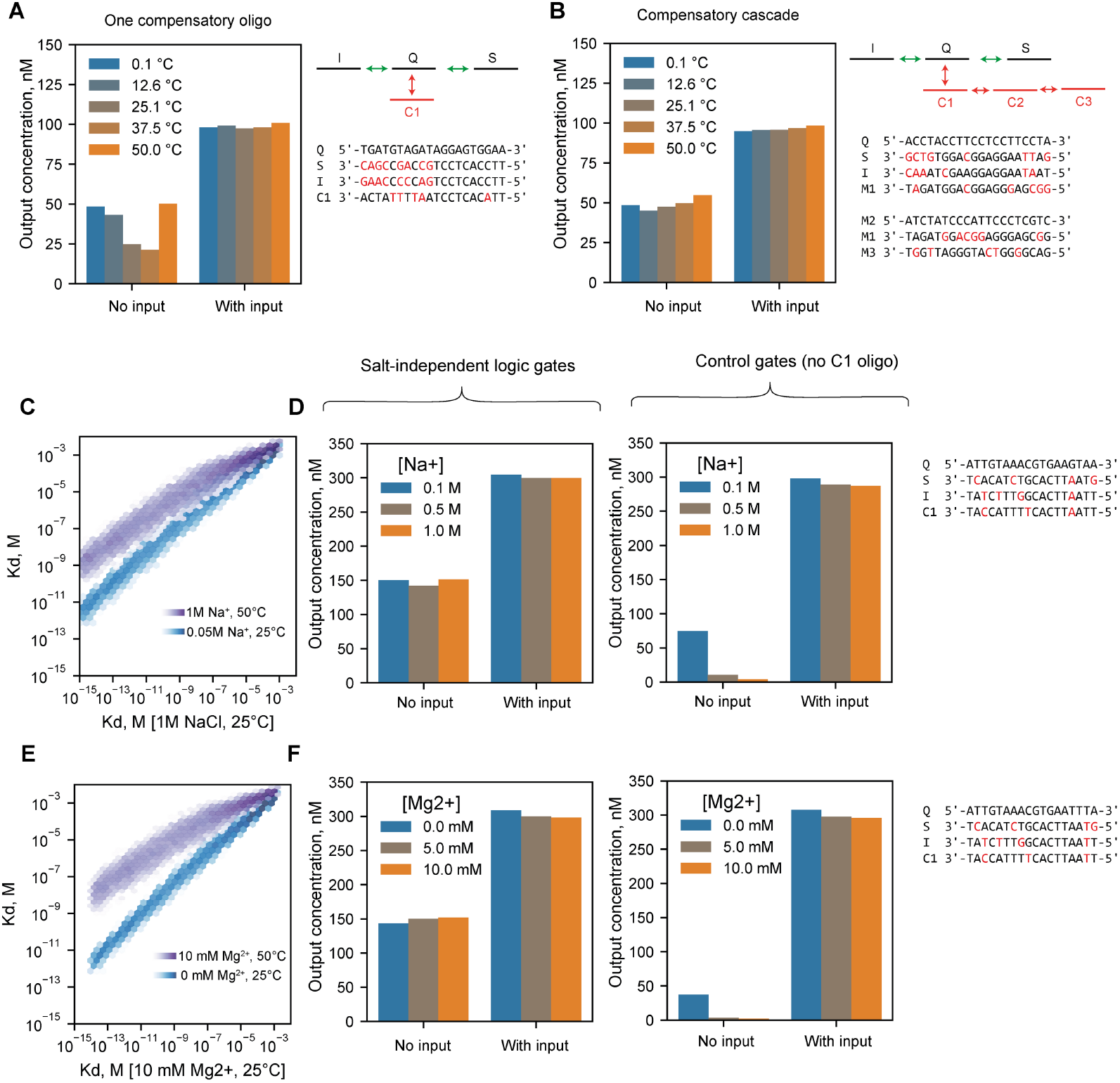
Predicted performance of gates across a 50 °C temperature range and under varying salt concentrations. (A) Predicted performance of a gate with a single compensatory oligonucleotide at intermediate temperatures between 0.1 °C and 50 °C. (B) Predicted performance of a gate with a compensatory cascade across the 0.1 °C–50 °C temperature range. (C) Distribution of dissociation constants (Kd) under high and low concentrations of Na^+^ and Mg^2+^, compared to the distribution caused by temperature variation for the same oligonucleotides. (D) Predicted performance of salt-independent YES gates across Na^+^ concentrations ranging from 0.1 M to 1 M. The left panel shows the performance of the optimized gate with compensatory oligonucleotide, while the right panel shows the same gate without compensation. (E, F) Same as (C, D), but for variation in Mg^2+^ concentration.

To address these deviations, we introduced a cascade of compensatory oligonucleotides in place of a single compensator (**Figure 4B**). In this configuration, C1 binds to Q, C2 binds to C1, and C3 binds to C2. The interaction between C1 and Q is designed to be the most “hard-to-melt” among the compensators—though still “easier-to-melt” than the core interactions between I–Q and Q–S —while each successive interaction in the cascade (Q-C1, C2–C1 and C3–C2) is progressively easier to melt.

As the temperature increases, the least stable interaction (C3–C2) dissociates first, followed by C2–C1, and finally C1–Q. This hierarchical compensation allows the system to gradually adjust to temperature changes, providing dynamic correction across the entire range. Using this strategy, we achieved stable and uniform gate performance at all intermediate temperatures. It is worth noting the exceptionally large change in the Kd of the participating reactions—just over 9 orders of magnitude across the temperature range from 0.1 to 50 °C.

In terms of hybridization efficiency, an increase in temperature has a similar effect to a decrease in ionic strength—both conditions lead to significantly weaker binding between oligonucleotide strands. To illustrate this, we plotted the distributions of dissociation constants (Kd) under low and high salt concentrations for two biologically relevant ions, Na^+^ and Mg^2+^, and compared them to Kd distributions at 25 °C and 50 °C (**Figure 4C, E**). For sodium, the concentration was varied from 50 mM to 1 M in the absence of Mg^2+^, while for magnesium, the concentration ranged from 0 to 10 mM in the presence of 50 mM Na^+^. These endpoint values were selected to align with the built-in limitations of the NUPACK thermodynamic model.

For both salts, the average shift in Kd resulting from a decrease in ionic strength is smaller than that caused by an increase in temperature. However, the characteristic width of the Kd distribution is also narrower under ionic strength variation. This implies that although the magnitude of compensation required is lower in the case of salt concentration changes, the diversity of available “easy-to-melt” and “hard-to-melt” oligonucleotides is also more limited.

To design a YES gate capable of functioning across a range of salt concentrations, we used the previously optimized sequences for the temperature-independent YES gate (**Figure 2A**) as a starting point. We then applied an evolutionary algorithm to refine both the sequences and their concentrations for robustness under varying ionic conditions. Using this approach, we successfully identified functional designs for both sodium and magnesium concentration variations, resulting in salt-independent logic gates (**Figure 4D, F**).

### Receptor activation in complex networks

Up to this point, we have focused on biochemical systems designed to respond to changes in the concentration of one or more molecular participants. We now turn our attention to a model receptor–activator system (**Figure 5A**). In this setup, we consider that binding of the activator to its corresponding receptor activates the receptor, while dissociation of the complex returns the receptor to its inactive state. There are many possible implementations of such systems in nature; for example, activation may occur due to conformational changes in a receptor upon its binding to a ligand^28^. Here, we treat the receptor activation level as the “output” of the system, by analogy with logic gates.

**Figure 5.**
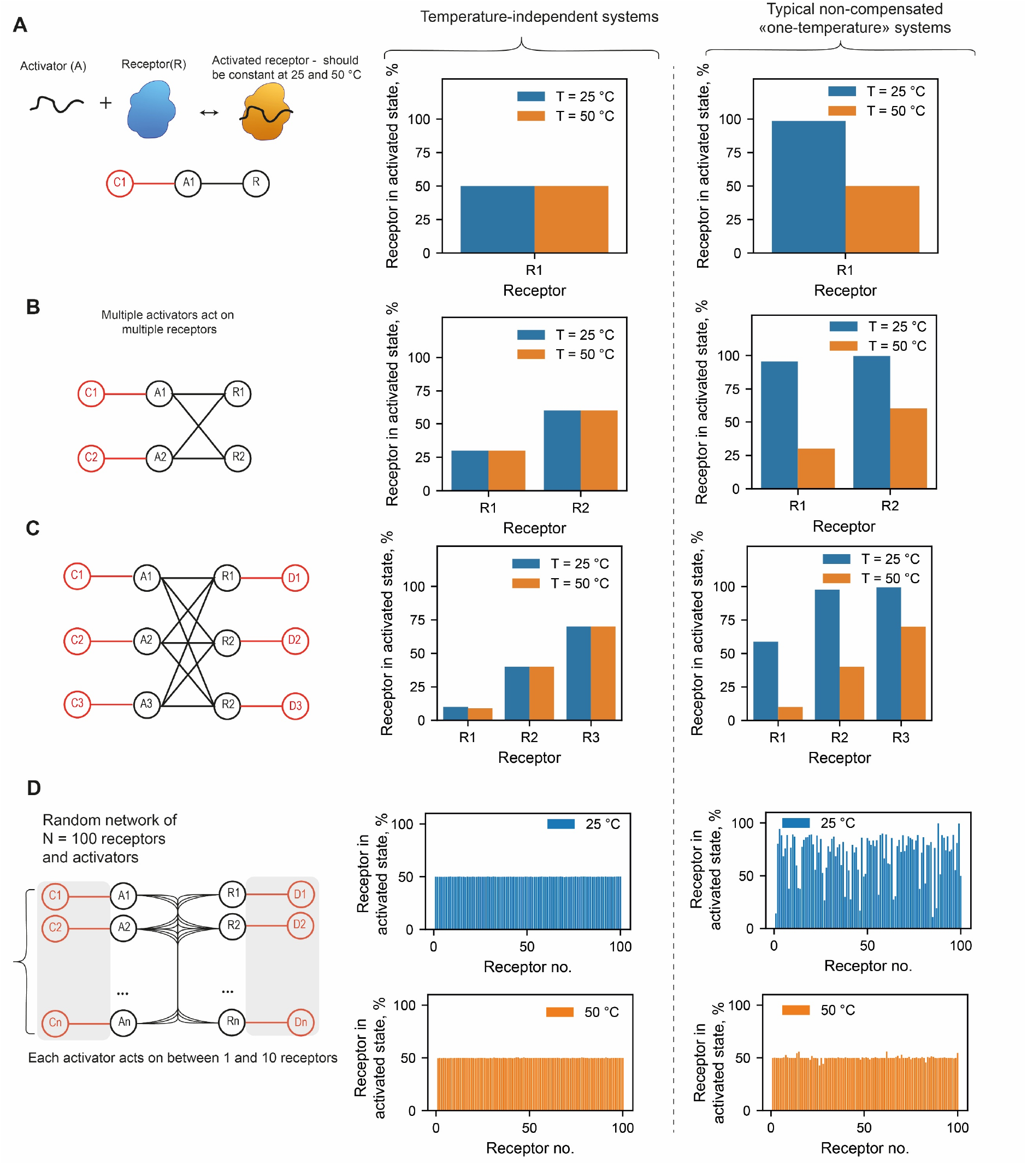
Temperature-independence of receptor–activator networks. (A–C) Network topology and predicted percentage of activated receptors for systems with 1, 2, and 3 receptor–activator pairs. Each receptor is assigned a predefined target activation level. (D) Topology and predicted performance of a large-scale network with 100 receptors and 100 activators. Each receptor is regulated by between 1 and 10 activators. For clarity, all receptors are assigned a 50% activation target. For (A-D) left panels show optimized systems with compensatory components; right panels show systems without compensation, optimized separately to have predefined activation levels at 50 °C only.

Suppose that the level of receptor activation must remain constant across varying external conditions—for example, across a range of temperatures. As in previous sections, we focus on two representative temperature points: 25 °C and 50 °C.

In the simple case of a single receptor (R1) and a single activator (A1), temperature-induced changes can be effectively compensated using just one additional component, C1. As temperature increases, both the A1–R1 and C1–A1 interactions weaken. On one hand, the reduced affinity between A1 and R1 leads to an increase in the concentration of free (and thus non-active) R1, resulting in a lower level of receptor activation. On the other hand, the weakened interaction with C1 increases the concentration of free A1, which in turn promotes more binding to R1 and counteracts the temperature effect. If the strengths of these interactions are properly optimized, the opposing effects can effectively cancel each other out, maintaining a stable level of receptor activation across temperatures. In our specific case, the target activation level of R1 is set arbitrarily to 50% (**Figure 5A**).

We can now extend this system to a more complex case involving two receptors (R1 and R2) and two activators (A1 and A2), where each receptor can be activated by either activator. Let us fix arbitrary target activation levels—for example, 30% activation for R1 and 60% for R2. These activation states can be maintained across temperatures (25 °C and 50 °C) using a similar compensation strategy. Specifically, we introduce two compensatory components, C1 and C2, which bind to A1 and A2, respectively. By tuning the interaction strengths of these compensators, the system can achieve temperature-independent activation levels for both receptors (**Figure 5B**).

In the more complex case involving three receptors and three activators, compensation requires an additional design consideration. In this scenario, we introduce compensatory participants that bind to the receptors without inducing activation, thereby competing with the activators for receptor binding sites. These non-activating competitors effectively modulate the concentration of free receptors available for activation, helping to balance the system’s response across temperatures. Using this strategy, we successfully maintained arbitrary, predefined activation levels for all three receptors (10%, 40% and 70% for R1, R2 and R3) at both 25 °C and 50 °C (**Figure 5C**).

The capabilities of the described approach for N = 3 prove to be remarkably extensive. Let us consider a complex network comprising 100 activators and 100 receptors, with each activator acting on between one and ten distinct receptors. As in previous cases, each activator and receptor is influenced by a single compensatory component. For simplicity, we assume that the activation level of each receptor must be precisely 50% at both temperatures.

To determine the optimal interaction parameters, approximately 700 affinities and 400 concentrations must be adjusted. The dimensionality of this problem significantly exceeds the capacity of the optimization algorithms employed. Therefore, we adopted the following strategy. We identified the three receptors with the largest deviations from the target activation levels and performed optimization of the interaction affinities with all associated activators, their individual compensators, as well as the compensators of the corresponding activators. These steps were alternated with the optimization of the concentrations of the same components. The receptors with the greatest deviations were then re-identified, and the process was iterated until the desired accuracy was achieved. The distribution of the number of activators acting on a single receptor is shown in **Figure S3**, with an average of 5.

Using this algorithm, we were able to attain a high degree of consistency in receptor activation levels across all 100 receptors within a few hundred optimization cycles, and the mean deviation from 50% level was less than 0.2% (**Figure 5D**). This demonstrates that the compensation approach remains effective even in systems with increased interaction complexity and multiple competing components.

For all four receptor–activator network cases, we also examine typical control systems—lacking compensatory oligos and optimized for operation at a single temperature (50 °C). In all cases, the control systems show substantial differences in activation levels at 25 and 50 °C.

### Algebraic systems

In this section, we explore more advanced biocomputing systems capable of exhibiting complex responses to varying levels of input signals. Consider an input strand I, an output strand S, and an arbitrary black-box set of intermediate components M1, M2, …, M_n_. We define a normalized and centered input variable X as:

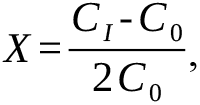

Where C_I_ is the total concentration of input I, and C_0_ is an arbitrarily chosen reference concentration (set to 5 µM). Similarly, the output variable Y is defined as the centered equilibrium concentration of free S

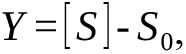

where S_0_ is reference concentration, set to 20 nM. Previous work^14^ has shown that by carefully selecting the sequences and concentrations of I, S, and all intermediate participants M_i_ it is possible to engineer complex nonlinear relationships between Y and X. Specific parameter sets have been identified that allow the system to approximate mathematical functions such as Y = X^3^ or Y = sin(X).

We now demonstrate that such complex biocomputing systems can also be made temperature-independent. To achieve this, we first optimize the system to function at 50 °C. Specifically, we minimize the mean squared error between the system’s output and a predefined target function by adjusting the dissociation constants and initial concentrations of N = 10 oligonucleotides using Bayesian optimization.

We begin by optimizing the dissociation constants and concentrations of the compensatory oligonucleotides at the lower temperature. Next, we perform a global optimization of all interactions and concentrations across both temperatures. In all optimization stages, dissociation constants (Kd) and concentrations are adjusted in an alternating manner: initially, concentrations are held constant while Kd values are optimized; then Kd values are fixed and concentrations are optimized. This iterative process continues until convergence.

After each step, we apply a pruning criterion: if a concentration falls below a defined threshold (0.1 nM) or a Kd value becomes too weak (>10^−4^ M), the corresponding variable is excluded from further optimization. In cases where an oligonucleotide’s concentration drops below the threshold, not only is the concentration removed from the optimization process, but all of its associated interactions are frozen as well—effectively eliminating it from the system.

Using this strategy, we successfully identified system parameters that enable robust computation of linear, quadratic and cubic functions at both 25 °C and 50 °C (**Figure 6B–D**). For comparison, we also show the typical behavior of systems optimized only at high temperature. While they perform ideally at 50 °C, they effectively cease to function computationally at low temperature.

**Figure 6.**
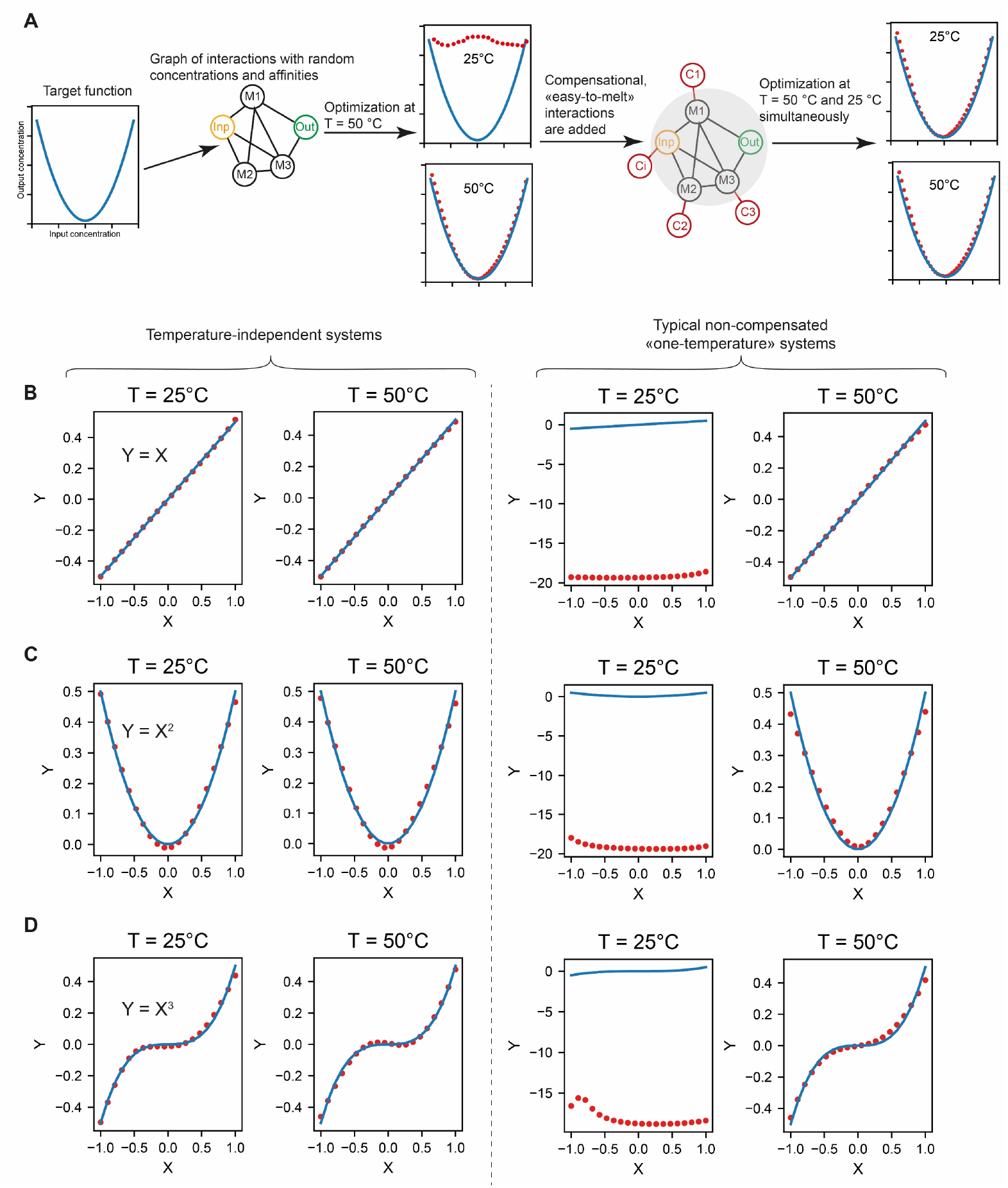
Computation of algebraic functions by biomolecular networks, independent of temperature. (A) Schematic of the optimization algorithm. (B–D) Predicted performance of corresponding systems implementing linear, quadratic, and cubic functions at 25 °C and 50 °C. Left panels show optimized systems with compensatory components; right panels show systems without compensation, optimized separately to follow the target function at 50 °C only.

Thus, we have shown that topological compensation can render even complex nonlinear biocomputational systems temperature-independent. Unlike logic gates, which were designed using a rational approach, these systems were obtained through stochastic methods. While rational approaches are easier for human understanding and experimental implementation, probabilistic, evolution-like algorithms more closely resemble the mechanisms by which such systems might have emerged in living organisms.

## Discussion

The phenomenon of topological compensation described above allows a reconsideration of both key evolutionary mechanisms and contemporary regulatory systems.

Currently, the RNA world hypothesis is recognized as one of the fundamental stages in the emergence of life^29^. Within this framework, RNA molecules form complex interacting reaction networks capable of maintaining their functionality, increasing complexity, and self-replication. Evidently, life in such a form was insufficiently developed to actively sustain constant physicochemical conditions. However, a broader range of permissible environmental conditions provides substantial evolutionary advantages, particularly in a continuously changing environment. In this context, topological compensation could have been a highly efficient and evolutionarily simple mechanism for maintaining homeostasis. Consequently, this mechanism may have played a crucial role in the development of all subsequent forms of life.

In modern organisms, complex biomolecular interaction networks are becoming increasingly recognized as critical for cellular regulation^17,30,31^, as evidenced by the growing number of recent studies and publications exploring this topic. Both in prokaryotes and eukaryotes, a significant role is played by various small RNAs, whose interactions with messenger RNA substantially define the cellular protein profile^32^. Furthermore, the extensive diversity of small RNAs within a single cell heightens the probability of their mutual interactions, leading to the formation of complex networks with strongly nonlinear behavior. For successful survival under stressful conditions, the cell must sustain the functioning of this network, as with any other vital system. For example, to protect protein systems, an entire class of protective heat-shock proteins has evolved^33^, mitigating the adverse effects of elevated temperatures on the folding of other proteins. By analogy, certain components of nucleic acid interaction networks may fulfill purely compensatory functions to safeguard these systems.

Moreover, it is currently hypothesized that such networks could be formed not only by nucleic acids but also by numerous critical proteins, including transcription factors capable of dimerization^11,34^. In our study, we demonstrate the realization of a compensatory mechanism using nucleic acids as an example. However, none of the essential assumptions are exclusive to nucleic acids. Indeed, we merely assume that the dimerization reaction behavior at low temperatures does not fully determine behavior at higher temperatures. This fact is equivalent to asserting the independence of interaction enthalpy and entropy, a condition fulfilled by the vast majority of biological interactions. Therefore, the same principles of topological compensation are equally applicable to protein-protein interactions.

## Conclusion

The described mechanism of topological compensation proves existence of non-insulated molecular systems that are capable of signal processing inherently resistant to dramatic fluctuations in physical conditions. The fundamental nature of these self-sustaining systems suggests that they may play an important regulatory role in a wide variety of living organisms, while the simplicity of their design implies the possibility of their spontaneous evolutionary emergence. Further investigation and development of such systems may give us not only better understanding of molecular signaling in living organisms, but also inspire the design of robust biotechnological processes that are more productive, compact, and less demanding of complex instrumentation to keep constant production conditions.

## Materials and methods

### Reagents

Oligonucleotides were synthesized by Lumiprobe (Moscow, Russia). Oligonucleotides modified with fluorophores or fluorescence quenchers were purified by polyacrylamide gel electrophoresis, while unmodified oligonucleotides were purified using C18 cartridges. Oligonucleotide solutions were prepared in a buffer composed of 0.1 M Tris, 1 M NaCl, 0.1 mM EDTA, 0.1 g/L Tween 20, pH 8. The reagents were obtained from the following suppliers: Dia-M (Moscow, Russia): Tris(hydroxymethyl)aminomethane; Helicon (Moscow, Russia): Sodium chloride; Applichem Panreac (Darmstadt, Germany): Ethylenediaminetetraacetic acid disodium salt dihydrate, Tween 20.

### Logic gates

Parameters of all logic gates and other systems are given in Supplementary Note 1 (except for the gates in Figure 5D, whose parameters are provided in a separate file, “Supplementary Data Figure 5D.xlsx”).

### Approximation of the temperature dependence of oligonucleotide dissociation constants

For the analysis of this relationship, we generated 10 000 pairs of 20-mer oligonucleotides. Each duplex contained between 1 and 10 mismatches, and the secondary-structure folding free energy of all oligonucleotides was constrained to zero.

For each pair, we determined the dissociation constant at 25 °C and 50 °C, converted these values to Gibbs free energies, and—assuming enthalpy and entropy to be temperature-independent— computed the corresponding enthalpies and entropies. Each oligonucleotide pair is therefore characterized by two parameters: the Gibbs free energy at 25 °C (G25) and the enthalpy (H).

We then constructed the frequency distribution of observed G25 values and partitioned it into 200 equal bins. For each bin (within which G25 varies negligibly), we derived the frequency distribution of H. From each such H distribution, we extracted the 5th, 10th, 20th, …, 80th, 90th, and 95th percentiles. By aggregating these percentile data across all G25 bins, we obtained the dependence of each percentile on G25, which we subsequently approximated using cubic B-splines (coefficients of these splines are listed in **Table S1**). Examples of obtained data and resulting approximations are given in **Figure S3** for 5th, 50th and 95th percentiles.

This approach enables prediction of possible enthalpy values—and hence Gibbs free energies at arbitrary temperatures—for any given G25. For example, the spline corresponding to the 5th percentile (p5) predicts enthalpies for the most “easy-to-melt” duplexes, whereas the 95th-percentile spline (p95) applies to the most “hard-to-melt” duplexes. These cubic-spline functions are denoted as p5, p10, etc.

### Affinity Measurement (Figure 3B)

A 40 µL mixture of oligonucleotides S and Q was added to 40 µL of oligonucleotide I, resulting in final concentrations of 100 nM for both S and Q, and 1 µM for I. The mixture was incubated at 50 °C for 1 hour, after which fluorescence was measured using a Clariostar spectrophotometer (BMG Labtech, Ortenberg, Germany) preheated to 50 °C. Excitation and emission wavelengths were set to 530 nm (bandwidth 20 nm) and 580 nm (bandwidth 30 nm), respectively.

Samples were then allowed to cool to room temperature for 2 hours, after which fluorescence measurements were repeated.

The dissociation constant K_d_ (SQ) was determined from the fluorescence level of the S and Q mixture in the absence of I. Using this value, the fluorescence levels of the samples containing varying concentrations of I were converted to corresponding dissociation constants K_d_(IQ). These constants were then converted to arbitrary C_AB_ units, which are plotted in Figure 2B according to the following relation:

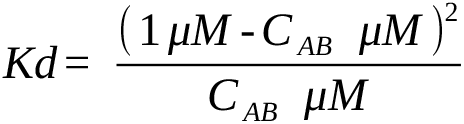

### NUPACK predictions

The NUPACK package for the Python programming language (version 4.0.1.12) was employed in this study. The method for computing equilibrium constants was adapted from the original publication describing the fundamental algorithms of the service^35^:

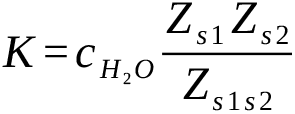

where the indices s1 and s2 refer to the interacting oligonucleotides, Z denotes the partition function of the respective complex, and 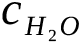 is the molar concentration of water. The partition function was computed as the pfunc attribute of the nupack.complex_analysis object. In all calculations, the maximum number of complexes parameter was set to 2.

### Equilibrium Concentration Calculations

Given the known dissociation constants and initial concentrations of oligonucleotides, the equilibrium concentrations of complexes were computed using the iterative method described by Dorp et al. ^36^

### Melting Curve Measurements (Figure 3C)

Melting curves were recorded using the LumoTrace Fluo bioimaging system equipped with a temperature control module. The heating rate was 0.15 °C/min. Fluorescence was measured once per minute with a 1-second exposure time, using a 550 ± 25 nm bandpass excitation filter and a 600 nm longpass emission filter.

## Declaration of interests

M.P.N. is stakeholder of Abisense LLC that manufactures the LumoTrace FLUO bioimaging system. M.P.N. is the inventor and the applicant of a related patent application RU2756476C2.

### Declaration of generative AI and AI-assisted technologies in the writing process

During the preparation of this work the authors used generative AI in order to translate parts of the original draft from Russian into English. After using this tool, the authors reviewed and edited the content as needed and take full responsibility for the content of the published article.

## Supporting information

Main supplementary information

Supplementary data for Figure 5D

## Acknowledgements

This research was supported by the Ministry of Science and Higher Education of the Russian Federation agreement 075-03-2025-662, project FSMG-2023-0017.

## Authors contributions

M.P.N. conceived the idea; M.P.N. and E.S.K. designed the study; E.S.K. wrote the code, carried out optimizations and experiments, visualized data, wrote original draft; M.P.N. edited manuscript and supervised the project.

## References

(1) Roncarati, D.; Vannini, A.; Scarlato, V. Temperature Sensing and Virulence Regulation in Pathogenic Bacteria. Trends Microbiol 2025, 33 (1), 66–79. 10.1016/j.tim.2024.07.009.

(2) Álvarez, M.; Molina, A.; Quezada, C.; Pinto, R.; Krauskopf, M.; Vera, M. I. Eurythermal Fish Acclimatization and Nucleolar Function: A Review. J Therm Biol 2004, 29 (7–8), 663–667. 10.1016/j.jtherbio.2004.08.036.

(3) Crawford, D. L.; Schulte, P. M.; Whitehead, A.; Oleksiak, M. F. Evolutionary Physiology and Genomics in the Highly Adaptable Killifish (Fundulus Heteroclitus). In Comprehensive Physiology; Wiley, 2020; pp 637–671. 10.1002/cphy.c190004.

(4) Zafren, K. Nonfreezing Cold Injury (Trench Foot). Int J Environ Res Public Health 2021, 18 (19), 10482. 10.3390/ijerph181910482.

(5) Maslov, S.; Ispolatov, I. Propagation of Large Concentration Changes in Reversible Protein-Binding Networks. Proceedings of the National Academy of Sciences 2007, 104 (34), 13655–13660. 10.1073/pnas.0702905104.

(6) Heo, M.; Maslov, S.; Shakhnovich, E. Topology of Protein Interaction Network Shapes Protein Abundances and Strengths of Their Functional and Nonspecific Interactions. Proceedings of the National Academy of Sciences 2011, 108 (10), 4258–4263. 10.1073/pnas.1009392108.

(7) Katz, E.; Privman, V. Enzyme-Based Logic Systems for Information Processing. Chem Soc Rev 2010, 39 (5), 1835. 10.1039/b806038j.

(8) Srinivas, N.; Parkin, J.; Seelig, G.; Winfree, E.; Soloveichik, D. Enzyme-Free Nucleic Acid Dynamical Systems. Science (1979) 2017, 358 (6369). 10.1126/science.aal2052.

(9) Nikitin, M. P.; Shipunova, V. O.; Deyev, S. M.; Nikitin, P. I. Biocomputing Based on Particle Disassembly. Nat Nanotechnol 2014, 9 (9), 716–722. 10.1038/nnano.2014.156.

(10) Tregubov, A. A.; Nikitin, P. I.; Nikitin, M. P. Advanced Smart Nanomaterials with Integrated Logic-Gating and Biocomputing: Dawn of Theranostic Nanorobots. Chem Rev 2018, 118 (20), 10294–10348. 10.1021/acs.chemrev.8b00198.

(11) Antebi, Y. E.; Linton, J. M.; Klumpe, H.; Bintu, B.; Gong, M.; Su, C.; McCardell, R.; Elowitz, M. B. Combinatorial Signal Perception in the BMP Pathway. Cell 2017, 170 (6), 1184–1196.e24. 10.1016/j.cell.2017.08.015.

(12) Wu, C.; Xu, M.; Dong, J.; Cui, W.; Yuan, S. The Structure and Function of Olfactory Receptors. Trends Pharmacol Sci 2024, 45 (3), 268–280. 10.1016/j.tips.2024.01.004.

(13) TerAvest, M. A.; Li, Z.; Angenent, L. T. Bacteria-Based Biocomputing with Cellular Computing Circuits to Sense, Decide, Signal, and Act. Energy Environ Sci 2011, 4 (12), 4907. 10.1039/c1ee02455h.

(14) Nikitin, M. P. Non-Complementary Strand Commutation as a Fundamental Alternative for Information Processing by DNA and Gene Regulation. Nat Chem 2023, 15 (1), 70–82. 10.1038/s41557-022-01111-y.

(15) Nikitin, M. P. Molecular Computing Device Based on Essentially Non-Complementary Single Stranded Nucleic Acids with Low Mutual Affinity. Patent application WO2021137740A2, 2021 (filed 2019).

(16) Nobeli, I.; Favia, A. D.; Thornton, J. M. Protein Promiscuity and Its Implications for Biotechnology. Nat Biotechnol 2009, 27 (2), 157–167. 10.1038/nbt1519.

(17) Parres-Gold, J.; Levine, M.; Emert, B.; Stuart, A.; Elowitz, M. B. Contextual Computation by Competitive Protein Dimerization Networks. Cell 2025, 188 (7), 1984–2002.e17. 10.1016/j.cell.2025.01.036.

(18) Tkachenko, A. V; Mognetti, B. M.; Maslov, S. Evolutionary Chemical Learning in Dimerization Networks.

(19) Sun, C.; Liu, X.; Zhong, J.; Zhou, Q.; Cheng, J. A Reusable Non-Complementary-DNA-Based Neural Network. 2024.

(20) Cai, H.; Zhang, X.; Qiao, R.; Wang, X.; Wei, L. Efficient Computation by Molecular Competition Networks. Phys Rev Res 2024, 6 (3), 1–7. 10.1103/PhysRevResearch.6.033208.

(21) Siebenmorgen, T.; Zacharias, M. Computational Prediction of Protein–Protein Binding Affinities. WIREs Computational Molecular Science 2020, 10 (3). 10.1002/wcms.1448.

(22) Fornace, M. E.; Porubsky, N. J.; Pierce, N. A. A Unified Dynamic Programming Framework for the Analysis of Interacting Nucleic Acid Strands: Enhanced Models, Scalability, and Speed. ACS Synth Biol 2020, 9 (10), 2665–2678. 10.1021/acssynbio.9b00523.

(23) Fornace, M. E.; Huang, J.; Newman, C. T.; Porubsky, N. J.; Pierce, M. B.; Pierce, N. A. NUPACK: Analysis and Design of Nucleic Acid Structures, Devices, and Systems. November 11, 2022. 10.26434/chemrxiv-2022-xv98l.

(24) Acerbi, L.; Ma, W. J. Practical Bayesian Optimization for Model Fitting with Bayesian Adaptive Direct Search. Adv Neural Inf Process Syst 2017, 2017-Decem, 1837–1847.

(25) Singh, G. S.; Acerbi, L. PyBADS: Fast and Robust Black-Box Optimization in Python. J Open Source Softw 2024, 9 (94), 5694. 10.21105/joss.05694.

(26) Holmstrom, E. D.; Dupuis, N. F.; Nesbitt, D. J. Pulsed IR Heating Studies of Single-Molecule DNA Duplex Dissociation Kinetics and Thermodynamics. Biophys J 2014, 106 (1), 220–231. 10.1016/j.bpj.2013.11.008.

(27) Moreira, B. G.; You, Y.; Behlke, M. A.; Owczarzy, R. Effects of Fluorescent Dyes, Quenchers, and Dangling Ends on DNA Duplex Stability. Biochem Biophys Res Commun 2005, 327 (2), 473–484. 10.1016/j.bbrc.2004.12.035.

(28) Wess, J.; Han, S.-J.; Kim, S.-K.; Jacobson, K. A.; Li, J. H. Conformational Changes Involved in G-Protein-Coupled-Receptor Activation. Trends Pharmacol Sci 2008, 29 (12), 616–625. 10.1016/j.tips.2008.08.006.

(29) Bernhardt, H. S. The RNA World Hypothesis: The Worst Theory of the Early Evolution of Life (except for All the Others). Biol Direct 2012, 7 (1), 23. 10.1186/1745-6150-7-23.

(30) Badia-i-Mompel, P.; Wessels, L.; Müller-Dott, S.; Trimbour, R.; Ramirez Flores, R. O.; Argelaguet, R.; Saez-Rodriguez, J. Gene Regulatory Network Inference in the Era of Single-Cell Multi-Omics. Nat Rev Genet 2023, 24 (11), 739–754. 10.1038/s41576-023-00618-5.

(31) Wilkinson, A. C.; Nakauchi, H.; Göttgens, B. Mammalian Transcription Factor Networks: Recent Advances in Interrogating Biological Complexity. Cell Syst 2017, 5 (4), 319–331. 10.1016/j.cels.2017.07.004.

(32) Chen, X.; Rechavi, O. Plant and Animal Small RNA Communications between Cells and Organisms. Nat Rev Mol Cell Biol 2022, 23 (3), 185–203. 10.1038/s41580-021-00425-y.

(33) Lang, B. J.; Guerrero, M. E.; Prince, T. L.; Okusha, Y.; Bonorino, C.; Calderwood, S. K. The Functions and Regulation of Heat Shock Proteins; Key Orchestrators of Proteostasis and the Heat Shock Response. Arch Toxicol 2021, 95 (6), 1943–1970. 10.1007/s00204-021-03070-8.

(34) Buchler, N. E.; Cross, F. R. Protein Sequestration Generates a Flexible Ultrasensitive Response in a Genetic Network. Mol Syst Biol 2009, 5 (1). 10.1038/msb.2009.30.

(35) Dirks, R. M.; Bois, J. S.; Schaeffer, J. M.; Winfree, E.; Pierce, N. A. Thermodynamic Analysis of Interacting Nucleic Acid Strands. SIAM Review 2007, 49 (1), 65–88. 10.1137/060651100.

(36) van Dorp, M. G. A.; Berger, F.; Carlon, E. Computing Equilibrium Concentrations for Large Heterodimerization Networks. Phys Rev E 2011, 84 (3), 036114. 10.1103/PhysRevE.84.036114.

