## Supplementary material for "Topology of molecular networks offers signaling insusceptibility to temperature and ionic strength changes": Main supplementary information

^*^Corresponding author

### Supplementary figures

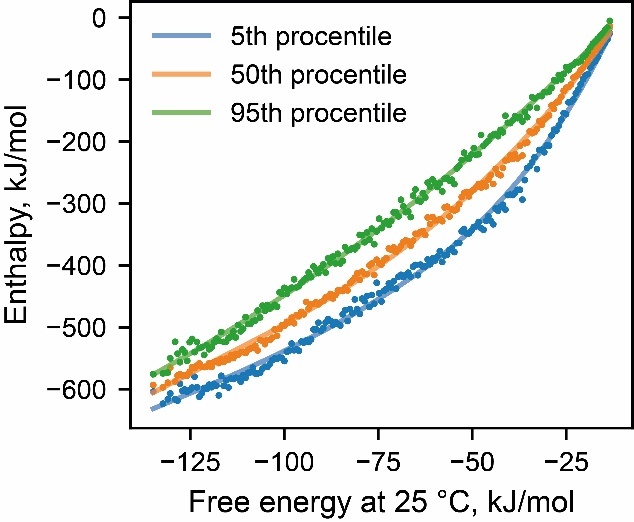

**Figure S1**. Scatter plot of enthalpy on Gibbs free energy at 25 °C and approximation with B-splines (solid lines) for 3 selected percentiles.

**Table S1**. Coefficients of cubic B-splines used to approximate dependence of enthalpy on Gibbs free energy at 25 °C. t refers to root vectors, and c are coefficients.

| **p5** | |  | **p10** | **p20** | **p30** | **p40** | **p50** |
| --- | --- | --- | --- | --- | --- | --- | --- |
| t | c | t | c | c | c | c | c |
| -134.8410 | -631.4712 | -134.8410 | -632.9235 | -622.5967 | -614.9400 | -609.8125 | -605.3929 |
| -134.8410 | -579.5258 | -134.8410 | -493.0558 | -489.7798 | -488.6400 | -483.5479 | -478.7154 |
| -134.8410 | -473.9449 | -134.8410 | -466.9375 | -436.3985 | -411.1192 | -390.7817 | -372.3745 |
| -134.8410 | -268.5936 | -134.8410 | -32.1964 | -28.6003 | -26.4071 | -25.3623 | -24.0606 |
| -72.1730 | -27.9923 | -13.3820 | 0.0000 | 0.0000 | 0.0000 | 0.0000 | 0.0000 |
| -13.3820 | 0.0000 | -13.3820 | 0.0000 | 0.0000 | 0.0000 | 0.0000 | 0.0000 |
| -13.3820 | 0.0000 | -13.3820 | 0.0000 | 0.0000 | 0.0000 | 0.0000 | 0.0000 |
| -13.3820 | 0.0000 | -13.3820 | 0.0000 | 0.0000 | 0.0000 | 0.0000 | 0.0000 |
| -13.3820 | 0.0000 |  |  |  |  |  |  |

**Table S1**, continued

|  | **p60** | **p70** | **p80** | **p90** | **p95** |
| --- | --- | --- | --- | --- | --- |
| t | c | c | c | c | c |
| -134.8410 | -600.8798 | -595.2991 | -590.1192 | -581.8567 | -576.4266 |
| -134.8410 | -475.4771 | -471.2282 | -461.2665 | -449.2162 | -440.2205 |
| -134.8410 | -352.0512 | -329.6438 | -306.6224 | -282.1742 | -262.7263 |
| -134.8410 | -23.4513 | -22.9364 | -22.4261 | -19.6115 | -17.7899 |
| -13.3820 | 0.0000 | 0.0000 | 0.0000 | 0.0000 | 0.0000 |
| -13.3820 | 0.0000 | 0.0000 | 0.0000 | 0.0000 | 0.0000 |
| -13.3820 | 0.0000 | 0.0000 | 0.0000 | 0.0000 | 0.0000 |
| -13.3820 | 0.0000 | 0.0000 | 0.0000 | 0.0000 | 0.0000 |

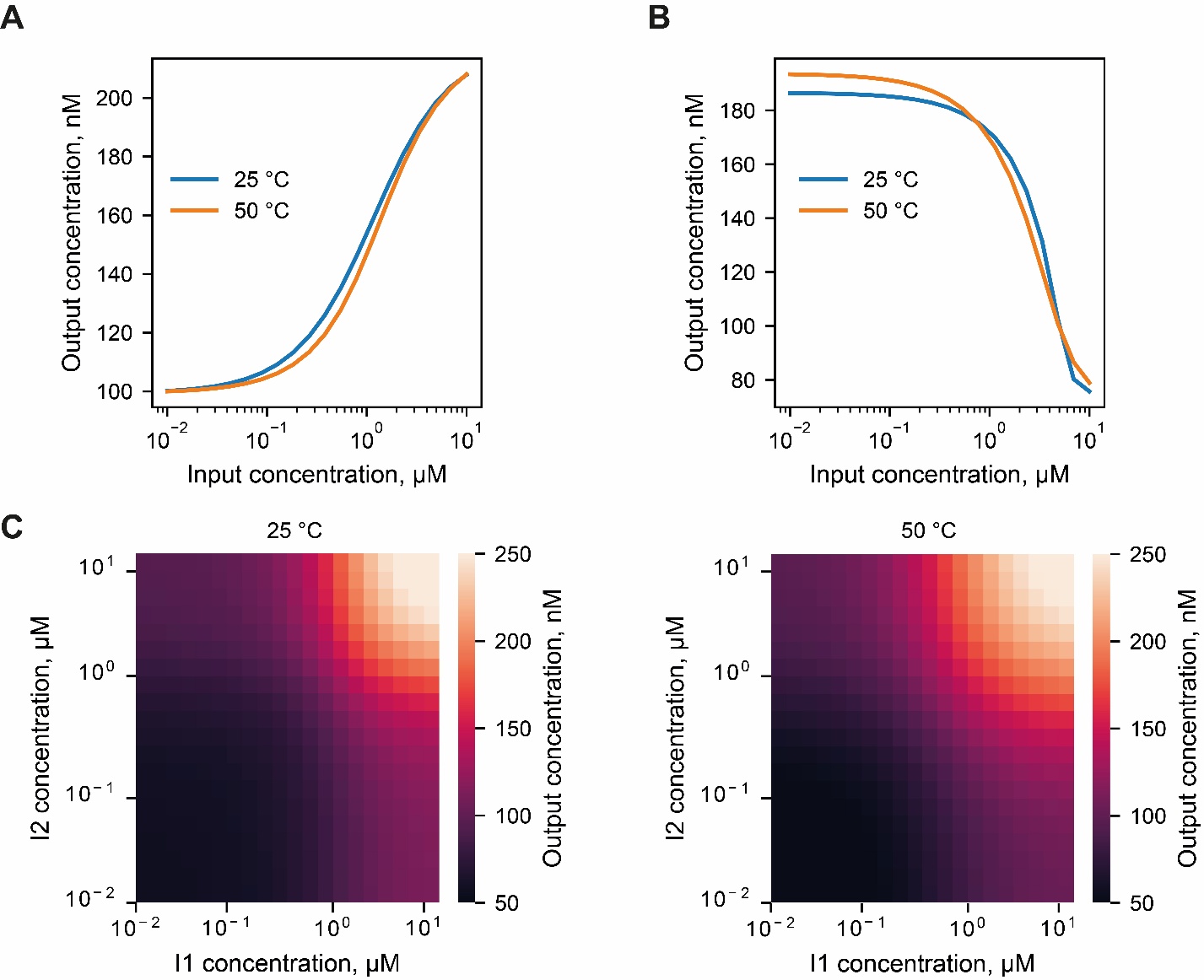

**Figure S2**. Logic gates performance under different input oligonucleotides concentrations. (A) YES-gate; (B) NOT-gate; (C) AND-gate.

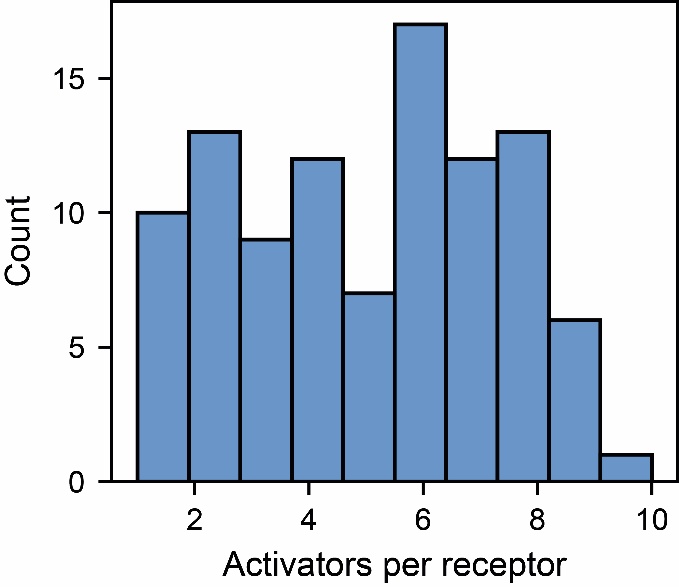

**Figure S3**. Distribution of activators per one receptor, related to **Figure 5D**.

#### Supplementary note 1: Parameters of the logic gates and other systems.

#### YES-gate

| ID | Sequence, 5'-3' | Concentration, μM | Hairpin formation energy at 25 °C, kcal/mol |
| --- | --- | --- | --- |
| I | TATTTTATGTTTTACCTGAA | 4.99 | 0.00 |
| Q | GTGAGGGAAAACATAAACGA | 0.70 | 0.00 |
| S | TCGTTTATGTTCTTATTATC | 0.22 | 0.00 |
| C | CCGTTTTAGATTTTCCTCAC | 1.50 | 0.00 |

Mutual affinity of the oligos. Concentration of the duplex C_AB_ (μM) when oligos are mixed at 1

μM each (at 25 °C), as predicted by NUPACK:

|  | I | Q | S | C |
| --- | --- | --- | --- | --- |
| I | 0.00 | 0.99 | 0.00 | 0.00 |
| Q | 0.99 | 0.00 | 0.99 | 0.99 |
| S | 0.00 | 0.99 | 0.00 | 0.00 |
| C | 0.00 | 0.99 | 0.00 | 0.00 |

Mutual affinity of the oligos. Concentration of the duplex C_AB_ (μM) when oligos are mixed at 1

μM each (at 50 °C), as predicted by NUPACK:

|  | I | Q | S | C |
| --- | --- | --- | --- | --- |
| I | 0.00 | 0.56 | 0.00 | 0.00 |
| Q | 0.56 | 0.00 | 0.52 | 0.04 |
| S | 0.00 | 0.52 | 0.00 | 0.00 |
| C | 0.00 | 0.04 | 0.00 | 0.00 |

Mutual affinity of the oligos. Approximate dissociation constant K_d_ at 25 °C in M, as predicted by NUPACK:

|  | I | Q | S | M |
| --- | --- | --- | --- | --- |
| I | 3.43E-03 | 7.35E-11 | 2.02E-03 | 2.87E-03 |
| Q | 7.35E-11 | 2.16E-04 | 9.68E-11 | 1.88E-10 |
| S | 2.02E-03 | 9.68E-11 | 1.31E-03 | 1.35E-03 |
| M | 2.87E-03 | 1.88E-10 | 1.35E-03 | 1.53E-03 |

Mutual affinity of the oligos. Approximate dissociation constant K_d_ at 50 °C in M, as predicted by NUPACK:

|  | I | Q | S | C |
| --- | --- | --- | --- | --- |
| I | 6.15E-03 | 3.52E-07 | 5.82E-03 | 5.99E-03 |
| Q | 3.52E-07 | 2.86E-03 | 4.46E-07 | 2.07E-05 |
| S | 5.82E-03 | 4.46E-07 | 5.96E-03 | 5.99E-03 |
| C | 5.99E-03 | 2.07E-05 | 5.99E-03 | 7.43E-03 |

#### NOT-gate

| ID | Sequence, 5'-3' | Concentration, μM | Hairpin formation energy at 25 °C, kcal/mol |
| --- | --- | --- | --- |
| I | TCTATCTTGCCTCTAATCTA | 5.55 | 0.00 |
| M | TAGCTGAGAGGAAAGATAGA | 5.48 | 0.00 |
| Q | CACGGGGTTACTCTCAGCAC | 4.05 | 0.00 |
| S | GTGCCAACAGTAAACCCGTG | 0.96 | -0.11 |
| C2 | TGTGTGTGAGTATGCCCCCG | 4.17 | 0.00 |
| C1 | AATATTAGAAGGTAGATAGT | 5.88 | 0.00 |

Mutual affinity of the oligos. Concentration of the duplex C_AB_ (μM) when oligos are mixed at 1

μM each (at 25 °C), as predicted by NUPACK:

|  | I | M | Q | S | C2 | C1 |
| --- | --- | --- | --- | --- | --- | --- |
| I | 0.00 | 0.99 | 0.00 | 0.00 | 0.00 | 0.94 |
| M | 0.99 | 0.00 | 0.98 | 0.00 | 0.00 | 0.00 |
| Q | 0.00 | 0.98 | 0.00 | 0.99 | 0.99 | 0.00 |
| S | 0.00 | 0.00 | 0.99 | 0.00 | 0.00 | 0.00 |
| C2 | 0.00 | 0.00 | 0.99 | 0.00 | 0.00 | 0.00 |
| C1 | 0.94 | 0.00 | 0.00 | 0.00 | 0.00 | 0.00 |

Mutual affinity of the oligos. Concentration of the duplex C_AB_ (μM) when oligos are mixed at 1

μM each (at 50 °C), as predicted by NUPACK:

|  | I | M | Q | S | C2 | C1 |
| --- | --- | --- | --- | --- | --- | --- |
| I | 0.00 | 0.65 | 0.00 | 0.00 | 0.00 | 0.01 |
| M | 0.65 | 0.00 | 0.37 | 0.00 | 0.00 | 0.00 |
| Q | 0.00 | 0.37 | 0.00 | 0.64 | 0.06 | 0.00 |
| S | 0.00 | 0.00 | 0.64 | 0.00 | 0.00 | 0.00 |
| C2 | 0.00 | 0.00 | 0.06 | 0.00 | 0.00 | 0.00 |
| C1 | 0.01 | 0.00 | 0.00 | 0.00 | 0.00 | 0.00 |

Mutual affinity of the oligos. Approximate dissociation constant K_d_ at 25 °C in M, as predicted by NUPACK:

|  | I | M | Q | S | C2 | C1 |
| --- | --- | --- | --- | --- | --- | --- |
| I | 2.81E-03 | 2.72E-11 | 8.05E-04 | 2.11E-03 | 5.86E-04 | 3.97E-09 |
| M | 2.72E-11 | 2.16E-04 | 4.59E-10 | 1.20E-03 | 2.69E-03 | 3.63E-03 |
| Q | 8.05E-04 | 4.59E-10 | 6.54E-04 | 2.84E-11 | 1.06E-10 | 1.22E-03 |
| S | 2.11E-03 | 1.20E-03 | 2.84E-11 | 1.49E-03 | 8.29E-04 | 1.16E-03 |
| C2 | 5.86E-04 | 2.69E-03 | 1.06E-10 | 8.29E-04 | 2.39E-03 | 2.29E-03 |
| C1 | 3.97E-09 | 3.63E-03 | 1.22E-03 | 1.16E-03 | 2.29E-03 | 7.68E-04 |

Mutual affinity of the oligos. Approximate dissociation constant K_d_ at 50 °C in M, as predicted by NUPACK:

|  | I | M | Q | S | C2 | C1 |
| --- | --- | --- | --- | --- | --- | --- |
| I | 6.55E-03 | 1.94E-07 | 3.41E-03 | 6.01E-03 | 4.27E-03 | 1.21E-04 |
| M | 1.94E-07 | 1.18E-03 | 1.08E-06 | 4.11E-03 | 4.80E-03 | 5.06E-03 |
| Q | 3.41E-03 | 1.08E-06 | 3.80E-03 | 1.97E-07 | 1.36E-05 | 3.79E-03 |
| S | 6.01E-03 | 4.11E-03 | 1.97E-07 | 5.40E-03 | 3.44E-03 | 3.78E-03 |
| C2 | 4.27E-03 | 4.80E-03 | 1.36E-05 | 3.44E-03 | 8.03E-03 | 6.67E-03 |
| C1 | 1.21E-04 | 5.06E-03 | 3.79E-03 | 3.78E-03 | 6.67E-03 | 3.45E-03 |

#### AND-gate

| ID | Sequence, 5'-3' | Concentration, μM | Hairpin formation energy at 25 °C, kcal/mol |
| --- | --- | --- | --- |
| I1 | GGGTGTGGTAACGAAGACCG | 5.00 | -1.40 |
| I2 | CTGTGAGGATACGTACAATG | 5.00 | -1.68 |
| Q1 | CGGTCTTCGTTTTTTCACTC | 0.74 | -0.95 |
| Q2 | CATTGTACGTATCTTTACTG | 0.74 | -1.22 |
| S | GAGTAAAGATACGAAGACCG | 0.31 | -1.40 |
| C1 | GGGTGACTAATCGAAGACCG | 0.74 | -1.58 |
| C2 | TAGTAATGATGCGTACAATA | 0.73 | -0.96 |
| C3 | GGGGTTTTGTTTCTTTGCTC | 0.31 | 0.00 |

Mutual affinity of the oligos. Concentration of the duplex C_AB_ (μM) when oligos are mixed at 1

μM each (at 25 °C), as predicted by NUPACK:

|  | I1 | I2 | Q1 | Q2 | S | C1 | C2 | C3 |
| --- | --- | --- | --- | --- | --- | --- | --- | --- |
| I1 | 0.00 | 0.00 | 0.99 | 0.00 | 0.00 | 0.00 | 0.00 | 0.00 |
| I2 | 0.00 | 0.00 | 0.00 | 0.99 | 0.00 | 0.00 | 0.00 | 0.00 |
| Q1 | 0.99 | 0.00 | 0.00 | 0.00 | 0.99 | 0.99 | 0.00 | 0.00 |
| Q2 | 0.00 | 0.99 | 0.00 | 0.01 | 0.99 | 0.00 | 0.99 | 0.00 |
| S | 0.00 | 0.00 | 0.99 | 0.99 | 0.00 | 0.00 | 0.00 | 0.13 |
| C1 | 0.00 | 0.00 | 0.99 | 0.00 | 0.00 | 0.00 | 0.00 | 0.00 |
| C2 | 0.00 | 0.00 | 0.00 | 0.99 | 0.00 | 0.00 | 0.00 | 0.00 |
| C3 | 0.00 | 0.00 | 0.00 | 0.00 | 0.13 | 0.00 | 0.00 | 0.00 |

Mutual affinity of the oligos. Concentration of the duplex C_AB_ (μM) when oligos are mixed at 1

μM each (at 50 °C), as predicted by NUPACK:

|  | I1 | I2 | Q1 | Q2 | S | C1 | C2 | C3 |
| --- | --- | --- | --- | --- | --- | --- | --- | --- |
| I1 | 0.00 | 0.00 | 0.67 | 0.00 | 0.00 | 0.00 | 0.00 | 0.00 |
| I2 | 0.00 | 0.00 | 0.00 | 0.67 | 0.00 | 0.00 | 0.00 | 0.00 |
| Q1 | 0.67 | 0.00 | 0.00 | 0.00 | 0.63 | 0.35 | 0.00 | 0.00 |
| Q2 | 0.00 | 0.67 | 0.00 | 0.00 | 0.66 | 0.00 | 0.30 | 0.00 |
| S | 0.00 | 0.00 | 0.63 | 0.66 | 0.00 | 0.00 | 0.00 | 0.00 |
| C1 | 0.00 | 0.00 | 0.35 | 0.00 | 0.00 | 0.00 | 0.00 | 0.00 |
| C2 | 0.00 | 0.00 | 0.00 | 0.30 | 0.00 | 0.00 | 0.00 | 0.00 |
| C3 | 0.00 | 0.00 | 0.00 | 0.00 | 0.00 | 0.00 | 0.00 | 0.00 |

Mutual affinity of the oligos. Approximate dissociation constant K_d_ at 25 °C in M, as predicted by NUPACK:

|  | I1 | I2 | Q1 | Q2 | S | C1 | C2 | C3 |
| --- | --- | --- | --- | --- | --- | --- | --- | --- |
| I1 | 1.79E-03 | 6.37E-03 | 3.59E-11 | 3.32E-03 | 8.97E-04 | 1.81E-03 | 5.28E-03 | 1.69E-03 |
| I2 | 6.37E-03 | 3.32E-04 | 3.15E-04 | 3.11E-11 | 2.87E-03 | 9.63E-03 | 1.31E-03 | 1.01E-03 |
| Q1 | 3.59E-11 | 3.15E-04 | 9.73E-03 | 2.83E-03 | 4.86E-11 | 3.21E-11 | 1.36E-03 | 4.32E-03 |
| Q2 | 3.32E-03 | 3.11E-11 | 2.83E-03 | 1.44E-04 | 4.49E-11 | 2.04E-03 | 3.50E-11 | 8.10E-03 |
| S | 8.97E-04 | 2.87E-03 | 4.86E-11 | 4.49E-11 | 5.06E-03 | 1.39E-03 | 1.79E-03 | 5.64E-06 |
| C1 | 1.81E-03 | 9.63E-03 | 3.21E-11 | 2.04E-03 | 1.39E-03 | 1.63E-03 | 3.82E-03 | 1.09E-03 |
| C2 | 5.28E-03 | 1.31E-03 | 1.36E-03 | 3.50E-11 | 1.79E-03 | 3.82E-03 | 1.08E-03 | 8.29E-04 |
| C3 | 1.69E-03 | 1.01E-03 | 4.32E-03 | 8.10E-03 | 5.64E-06 | 1.09E-03 | 8.29E-04 | 1.19E-03 |

Mutual affinity of the oligos. Approximate dissociation constant K_d_ at 50 °C in M, as predicted by NUPACK:

|  | I1 | I2 | Q1 | Q2 | S | C1 | C2 | C3 |
| --- | --- | --- | --- | --- | --- | --- | --- | --- |
| I1 | 3.93E-03 | 6.60E-03 | 1.67E-07 | 3.91E-03 | 4.49E-03 | 4.67E-03 | 7.45E-03 | 3.54E-03 |
| I2 | 6.60E-03 | 8.92E-04 | 2.60E-03 | 1.58E-07 | 3.70E-03 | 9.60E-03 | 3.48E-03 | 3.10E-03 |
| Q1 | 1.67E-07 | 2.60E-03 | 1.02E-02 | 4.82E-03 | 2.22E-07 | 1.18E-06 | 4.16E-03 | 9.33E-03 |
| Q2 | 3.91E-03 | 1.58E-07 | 4.82E-03 | 6.18E-04 | 1.76E-07 | 5.34E-03 | 1.66E-06 | 9.09E-03 |
| S | 4.49E-03 | 3.70E-03 | 2.22E-07 | 1.76E-07 | 8.08E-03 | 3.60E-03 | 3.62E-03 | 6.59E-04 |
| C1 | 4.67E-03 | 9.60E-03 | 1.18E-06 | 5.34E-03 | 3.60E-03 | 4.72E-03 | 7.04E-03 | 3.49E-03 |
| C2 | 7.45E-03 | 3.48E-03 | 4.16E-03 | 1.66E-06 | 3.62E-03 | 7.04E-03 | 5.00E-03 | 3.32E-03 |
| C3 | 3.54E-03 | 3.10E-03 | 9.33E-03 | 9.09E-03 | 6.59E-04 | 3.49E-03 | 3.32E-03 | 1.06E-02 |

#### 0.1-50 °С control YES-gate (one compensatory oligo, Figure 3A)

| ID | Sequence, 5'-3' | Concentration, μM | Hairpin formation energy at 25 °C, kcal/mol |
| --- | --- | --- | --- |
| I | TTCCACTCCTGACCCCCAAG | 2.83 | -0.48 |
| Q | TGATGTAGATAGGAGTGGAA | 0.50 | 0.00 |
| S | TTCCACTCCTGCCAGCCGAC | 0.12 | 0.00 |
| C1 | TTACACTCCTAATTTTATCA | 1.50 | 0.00 |

Mutual affinity of the oligos. Concentration of the duplex C_AB_ (μM) when oligos are mixed at 1

μM each (at 25 °C), as predicted by NUPACK:

|  | I | Q | S | C1 |
| --- | --- | --- | --- | --- |
| I | 0.00 | 0.99 | 0.00 | 0.00 |
| Q | 0.99 | 0.00 | 0.99 | 0.97 |
| S | 0.00 | 0.99 | 0.00 | 0.00 |
| C1 | 0.00 | 0.97 | 0.00 | 0.00 |

Mutual affinity of the oligos. Concentration of the duplex C_AB_ (μM) when oligos are mixed at 1

μM each (at 50 °C), as predicted by NUPACK:

|  | I | Q | S | C1 |
| --- | --- | --- | --- | --- |
| I | 0.00 | 0.64 | 0.00 | 0.00 |
| Q | 0.64 | 0.00 | 0.65 | 0.28 |
| S | 0.00 | 0.65 | 0.00 | 0.00 |
| C1 | 0.00 | 0.28 | 0.00 | 0.00 |

Mutual affinity of the oligos. Approximate dissociation constant K_d_ at 25 °C in M, as predicted by NUPACK:

|  | I | Q | S | C1 |
| --- | --- | --- | --- | --- |
| I | 8.13E-03 | 7.82E-11 | 3.40E-03 | 6.82E-03 |
| Q | 7.82E-11 | 3.70E-03 | 6.38E-11 | 6.64E-10 |
| S | 3.40E-03 | 6.38E-11 | 5.78E-04 | 5.46E-03 |
| C1 | 6.82E-03 | 6.64E-10 | 5.46E-03 | 1.48E-03 |

Mutual affinity of the oligos. Approximate dissociation constant K_d_ at 50 °C in M, as predicted by NUPACK:

|  | I | Q | S | C1 |
| --- | --- | --- | --- | --- |
| I | 1.00E-02 | 1.96E-07 | 7.26E-03 | 1.03E-02 |
| Q | 1.96E-07 | 7.64E-03 | 1.82E-07 | 1.89E-06 |
| S | 7.26E-03 | 1.82E-07 | 3.16E-03 | 1.10E-02 |
| C1 | 1.03E-02 | 1.89E-06 | 1.10E-02 | 5.53E-03 |

#### 0.1-50 °С YES-gate (with compensatory cascade, Figure 4B)

| ID | Sequence, 5'-3' | Concentration, μM | Hairpin formation energy at 25 °C, kcal/mol |
| --- | --- | --- | --- |
| I | TAATAAGGAGGAAGCTAAAC | 5.00 | -0.72 |
| Q | ACCTACCTTCCTCCTTCCTA | 0.97 | 0.00 |
| S | GATTAAGGAGGCAGGTGTCG | 0.12 | 0.00 |
| C1 | GGCGAGGGAGGCAGGTAGAT | 2.93 | -1.34 |
| C2 | ATCTATCCCATTCCCTCGTC | 2.80 | 0.00 |
| C3 | GACGGGGTCATGGGATTGGT | 5.00 | -1.03 |

Mutual affinity of the oligos. Concentration of the duplex C_AB_ (μM) when oligos are mixed at 1

μM each (at 25 °C), as predicted by NUPACK:

|  | I | Q | S | C1 | C2 | C3 |
| --- | --- | --- | --- | --- | --- | --- |
| I | 0.00 | 0.99 | 0.00 | 0.00 | 0.03 | 0.00 |
| Q | 0.99 | 0.00 | 0.99 | 0.99 | 0.00 | 0.10 |
| S | 0.00 | 0.99 | 0.00 | 0.00 | 0.05 | 0.00 |
| C1 | 0.00 | 0.99 | 0.00 | 0.00 | 0.96 | 0.00 |
| C2 | 0.03 | 0.00 | 0.05 | 0.96 | 0.00 | 0.99 |
| C3 | 0.00 | 0.10 | 0.00 | 0.00 | 0.99 | 0.00 |

Mutual affinity of the oligos. Concentration of the duplex C_AB_ (μM) when oligos are mixed at 1

μM each (at 50 °C), as predicted by NUPACK:

|  | I | Q | S | C1 | C2 | C3 |
| --- | --- | --- | --- | --- | --- | --- |
| I | 0.00 | 0.67 | 0.00 | 0.00 | 0.00 | 0.00 |
| Q | 0.67 | 0.00 | 0.70 | 0.54 | 0.00 | 0.00 |
| S | 0.00 | 0.70 | 0.00 | 0.00 | 0.00 | 0.00 |
| C1 | 0.00 | 0.54 | 0.00 | 0.00 | 0.20 | 0.00 |
| C2 | 0.00 | 0.00 | 0.00 | 0.20 | 0.00 | 0.38 |
| C3 | 0.00 | 0.00 | 0.00 | 0.00 | 0.38 | 0.00 |

Mutual affinity of the oligos. Approximate dissociation constant K_d_ at 25 °C in M, as predicted by NUPACK:

|  | I | Q | S | C1 | C2 | C3 |
| --- | --- | --- | --- | --- | --- | --- |
| I | 6.53E-04 | 3.76E-11 | 1.13E-03 | 3.77E-03 | 2.95E-05 | 2.40E-03 |
| Q | 3.76E-11 | 3.00E-02 | 2.68E-11 | 9.10E-11 | 6.65E-03 | 7.67E-06 |
| S | 1.13E-03 | 2.68E-11 | 8.69E-04 | 1.92E-03 | 1.89E-05 | 3.11E-03 |
| C1 | 3.77E-03 | 9.10E-11 | 1.92E-03 | 6.12E-03 | 1.37E-09 | 6.21E-03 |
| C2 | 2.95E-05 | 6.65E-03 | 1.89E-05 | 1.37E-09 | 1.82E-03 | 2.82E-11 |
| C3 | 2.40E-03 | 7.67E-06 | 3.11E-03 | 6.21E-03 | 2.82E-11 | 1.54E-03 |

Mutual affinity of the oligos. Approximate dissociation constant K_d_ at 50 °C in M, as predicted by NUPACK:

|  | I | Q | S | C1 | C2 | C3 |
| --- | --- | --- | --- | --- | --- | --- |
| I | 1.48E-03 | 1.64E-07 | 3.80E-03 | 7.16E-03 | 4.79E-04 | 6.59E-03 |
| Q | 1.64E-07 | 2.80E-02 | 1.30E-07 | 3.82E-07 | 1.59E-02 | 5.79E-04 |
| S | 3.80E-03 | 1.30E-07 | 4.42E-03 | 6.02E-03 | 4.88E-04 | 6.85E-03 |
| C1 | 7.16E-03 | 3.82E-07 | 6.02E-03 | 6.83E-03 | 3.23E-06 | 8.93E-03 |
| C2 | 4.79E-04 | 1.59E-02 | 4.88E-04 | 3.23E-06 | 9.15E-03 | 1.01E-06 |
| C3 | 6.59E-03 | 5.79E-04 | 6.85E-03 | 8.93E-03 | 1.01E-06 | 5.67E-03 |

#### Kinetics study

| **ID** | **Sequence, 5'-3'** | **Concentration, μM** |
| --- | --- | --- |
| S | [Cy3]GGCTTAACATATTTCAGGGG | 0.10 |
| Q1 | TCCCTGGATTATGTTCTGCG[BHQ2] | 0.10 |
| I1 | GGCTTAACATATTTCAGGGA | 0.10 |
| Q2 | CCCCCGAAATATGTCATCAC[BHQ2] | 0.10 |
| I2 | GGCATAACATATTTCAGGGG | 0.10 |
| Q3 | CCCCTGAAATATATTATGTA[BHQ2] | 0.10 |
| I3 | TGCTTAACATATTTCAGGGG | 0.10 |
| Q4 | CCCGTGAAATATAGTAAGCC[BHQ2] | 0.10 |
| I4 | GGCTTAATATATTTCAGGGG | 0.10 |
| Q5 | CCCCTATAATATGTTAAGCC[BHQ2] | 0.10 |
| I5 | GGCTTAACATATTACAGGGG | 0.10 |

For **Figure 3a** we used concentrations listed in in the table above. For the similar experiment (**Figure S2**) concentrations of all oligonucleotides was 10 nM.

Mutual affinity of the oligos. Concentration of the duplex C_AB_ (μM) when oligos are mixed at 1

μM each (at 25 °C), as predicted by NUPACK:

|  | **S** | **Q1** | **I1** | **Q2** | **I2** | **Q3** | **I3** | **Q4** | **I4** | **Q5** | **I5** |
| --- | --- | --- | --- | --- | --- | --- | --- | --- | --- | --- | --- |
| **S** | 0.0005 | 0.9451 | 0.0006 | 0.9869 | 0.0008 | 0.9957 | 0.0005 | 0.9993 | 0.0012 | 0.9999 | 0.0005 |
| **Q1** | 0.9451 | 0.0095 | 0.9614 | 0.0015 | 0.9850 | 0.0015 | 0.9429 | 0.0029 | 0.2930 | 0.0018 | 0.9812 |
| **I1** | 0.0006 | 0.9614 | 0.0006 | 0.9728 | 0.0008 | 0.9937 | 0.0005 | 0.9989 | 0.0014 | 0.9999 | 0.0005 |
| **Q2** | 0.9869 | 0.0015 | 0.9728 | 0.0019 | 0.9877 | 0.0013 | 0.9862 | 0.0039 | 0.8884 | 0.0002 | 0.7803 |
| **I2** | 0.0008 | 0.9850 | 0.0008 | 0.9877 | 0.0012 | 0.9995 | 0.0012 | 0.9866 | 0.0013 | 0.9979 | 0.0008 |
| **Q3** | 0.9957 | 0.0015 | 0.9937 | 0.0013 | 0.9995 | 0.0163 | 0.9957 | 0.0035 | 0.9998 | 0.0124 | 0.9090 |
| **I3** | 0.0005 | 0.9429 | 0.0005 | 0.9862 | 0.0012 | 0.9957 | 0.0005 | 0.9985 | 0.0012 | 0.9998 | 0.0005 |
| **Q4** | 0.9993 | 0.0029 | 0.9989 | 0.0039 | 0.9866 | 0.0035 | 0.9985 | 0.0019 | 0.9999 | 0.0017 | 0.9823 |
| **I4** | 0.0012 | 0.2930 | 0.0014 | 0.8884 | 0.0013 | 0.9998 | 0.0012 | 0.9999 | 0.0908 | 0.9918 | 0.0018 |
| **Q5** | 0.9999 | 0.0018 | 0.9999 | 0.0002 | 0.9979 | 0.0124 | 0.9998 | 0.0017 | 0.9918 | 0.0004 | 0.9999 |
| **I5** | 0.0005 | 0.9812 | 0.0005 | 0.7803 | 0.0008 | 0.9090 | 0.0005 | 0.9823 | 0.0018 | 0.9999 | 0.0006 |

Mutual affinity of the oligos. Concentration of the duplex C_AB_ (μM) when oligos are mixed at 1

μM each (at 50 °C), as predicted by NUPACK:

|  | **S** | **Q1** | **I1** | **Q2** | **I2** | **Q3** | **I3** | **Q4** | **I4** | **Q5** | **I5** |
| --- | --- | --- | --- | --- | --- | --- | --- | --- | --- | --- | --- |
| **S** | 0.00 | 0.06 | 0.00 | 0.50 | 0.00 | 0.79 | 0.00 | 0.67 | 0.00 | 0.95 | 0.00 |
| **Q1** | 0.06 | 0.00 | 0.12 | 0.00 | 0.13 | 0.00 | 0.05 | 0.00 | 0.02 | 0.00 | 0.32 |
| **I1** | 0.00 | 0.12 | 0.00 | 0.40 | 0.00 | 0.75 | 0.00 | 0.61 | 0.00 | 0.94 | 0.00 |
| **Q2** | 0.50 | 0.00 | 0.40 | 0.00 | 0.48 | 0.00 | 0.48 | 0.00 | 0.07 | 0.00 | 0.04 |
| **I2** | 0.00 | 0.13 | 0.00 | 0.48 | 0.00 | 0.89 | 0.00 | 0.20 | 0.00 | 0.60 | 0.00 |
| **Q3** | 0.79 | 0.00 | 0.75 | 0.00 | 0.89 | 0.00 | 0.79 | 0.00 | 0.94 | 0.00 | 0.22 |
| **I3** | 0.00 | 0.05 | 0.00 | 0.48 | 0.00 | 0.79 | 0.00 | 0.51 | 0.00 | 0.92 | 0.00 |
| **Q4** | 0.67 | 0.00 | 0.61 | 0.00 | 0.20 | 0.00 | 0.51 | 0.00 | 0.95 | 0.00 | 0.08 |
| **I4** | 0.00 | 0.02 | 0.00 | 0.07 | 0.00 | 0.94 | 0.00 | 0.95 | 0.00 | 0.22 | 0.00 |
| **Q5** | 0.95 | 0.00 | 0.94 | 0.00 | 0.60 | 0.00 | 0.92 | 0.00 | 0.22 | 0.00 | 0.98 |
| **I5** | 0.00 | 0.32 | 0.00 | 0.04 | 0.00 | 0.22 | 0.00 | 0.08 | 0.00 | 0.98 | 0.00 |

Mutual affinity of the oligos. Approximate dissociation constant K_d_ at 25 °C in M, as predicted by NUPACK:

|  | **S** | **Q1** | **I1** | **Q2** | **I2** | **Q3** | **I3** | **Q4** | **I4** | **Q5** | **I5** |
| --- | --- | --- | --- | --- | --- | --- | --- | --- | --- | --- | --- |
| **S** | 1.9E-03 | 3.2E-09 | 1.8E-03 | 1.7E-10 | 1.3E-03 | 1.8E-11 | 2.0E-03 | 5.5E-13 | 8.6E-04 | 8.6E-15 | 1.9E-03 |
| **Q1** | 3.2E-09 | 1.0E-04 | 1.6E-09 | 6.5E-04 | 2.3E-10 | 6.7E-04 | 3.5E-09 | 3.5E-04 | 1.7E-06 | 5.5E-04 | 3.6E-10 |
| **I1** | 1.8E-03 | 1.6E-09 | 1.7E-03 | 7.6E-10 | 1.2E-03 | 4.0E-11 | 1.9E-03 | 1.1E-12 | 7.3E-04 | 1.9E-14 | 1.8E-03 |
| **Q2** | 1.7E-10 | 6.5E-04 | 7.6E-10 | 5.2E-04 | 1.5E-10 | 7.8E-04 | 1.9E-10 | 2.5E-04 | 1.4E-08 | 5.2E-03 | 6.2E-08 |
| **I2** | 1.3E-03 | 2.3E-10 | 1.2E-03 | 1.5E-10 | 8.3E-04 | 2.6E-13 | 8.1E-04 | 1.8E-10 | 7.7E-04 | 4.5E-12 | 1.3E-03 |
| **Q3** | 1.8E-11 | 6.7E-04 | 4.0E-11 | 7.8E-04 | 2.6E-13 | 5.9E-05 | 1.8E-11 | 2.8E-04 | 5.1E-14 | 7.8E-05 | 9.1E-09 |
| **I3** | 2.0E-03 | 3.5E-09 | 1.9E-03 | 1.9E-10 | 8.1E-04 | 1.8E-11 | 2.0E-03 | 2.1E-12 | 8.6E-04 | 3.3E-14 | 2.0E-03 |
| **Q4** | 5.5E-13 | 3.5E-04 | 1.1E-12 | 2.5E-04 | 1.8E-10 | 2.8E-04 | 2.1E-12 | 5.2E-04 | 8.6E-15 | 5.8E-04 | 3.2E-10 |
| **I4** | 8.6E-04 | 1.7E-06 | 7.3E-04 | 1.4E-08 | 7.7E-04 | 5.1E-14 | 8.6E-04 | 8.6E-15 | 9.1E-06 | 6.8E-11 | 5.4E-04 |
| **Q5** | 8.6E-15 | 5.5E-04 | 1.9E-14 | 5.2E-03 | 4.5E-12 | 7.8E-05 | 3.3E-14 | 5.8E-04 | 6.8E-11 | 2.4E-03 | 2.7E-15 |
| **I5** | 1.9E-03 | 3.6E-10 | 1.8E-03 | 6.2E-08 | 1.3E-03 | 9.1E-09 | 2.0E-03 | 3.2E-10 | 5.4E-04 | 2.7E-15 | 1.8E-03 |

Mutual affinity of the oligos. Approximate dissociation constant K_d_ at 50 °C in M, as predicted by NUPACK:

|  | **S** | **Q1** | **I1** | **Q2** | **I2** | **Q3** | **I3** | **Q4** | **I4** | **Q5** | **I5** |
| --- | --- | --- | --- | --- | --- | --- | --- | --- | --- | --- | --- |
| **S** | 3.5E-03 | 1.6E-05 | 3.4E-03 | 5.1E-07 | 3.6E-03 | 5.5E-08 | 3.7E-03 | 1.7E-07 | 2.4E-03 | 2.4E-09 | 3.4E-03 |
| **Q1** | 1.6E-05 | 8.6E-04 | 6.8E-06 | 1.8E-03 | 6.1E-06 | 2.6E-03 | 1.7E-05 | 1.6E-03 | 6.3E-05 | 1.7E-03 | 1.5E-06 |
| **I1** | 3.4E-03 | 6.8E-06 | 3.3E-03 | 9.1E-07 | 3.5E-03 | 8.3E-08 | 3.5E-03 | 2.4E-07 | 2.3E-03 | 3.5E-09 | 3.3E-03 |
| **Q2** | 5.1E-07 | 1.8E-03 | 9.1E-07 | 2.1E-03 | 5.5E-07 | 2.6E-03 | 5.5E-07 | 1.6E-03 | 1.3E-05 | 4.9E-03 | 2.1E-05 |
| **I2** | 3.6E-03 | 6.1E-06 | 3.5E-03 | 5.5E-07 | 3.6E-03 | 1.5E-08 | 3.1E-03 | 3.2E-06 | 2.5E-03 | 2.6E-07 | 3.5E-03 |
| **Q3** | 5.5E-08 | 2.6E-03 | 8.3E-08 | 2.6E-03 | 1.5E-08 | 1.1E-03 | 5.7E-08 | 2.0E-03 | 3.5E-09 | 1.8E-03 | 2.8E-06 |
| **I3** | 3.7E-03 | 1.7E-05 | 3.5E-03 | 5.5E-07 | 3.1E-03 | 5.7E-08 | 3.8E-03 | 4.6E-07 | 2.5E-03 | 7.3E-09 | 3.5E-03 |
| **Q4** | 1.7E-07 | 1.6E-03 | 2.4E-07 | 1.6E-03 | 3.2E-06 | 2.0E-03 | 4.6E-07 | 2.3E-03 | 3.1E-09 | 2.3E-03 | 1.0E-05 |
| **I4** | 2.4E-03 | 6.3E-05 | 2.3E-03 | 1.3E-05 | 2.5E-03 | 3.5E-09 | 2.5E-03 | 3.1E-09 | 4.9E-04 | 2.8E-06 | 2.0E-03 |
| **Q5** | 2.4E-09 | 1.7E-03 | 3.5E-09 | 4.9E-03 | 2.6E-07 | 1.8E-03 | 7.3E-09 | 2.3E-03 | 2.8E-06 | 3.4E-03 | 3.2E-10 |
| **I5** | 3.4E-03 | 1.5E-06 | 3.3E-03 | 2.1E-05 | 3.5E-03 | 2.8E-06 | 3.5E-03 | 1.0E-05 | 2.0E-03 | 3.2E-10 | 3.3E-03 |

NUPACK prediction accuracy screening (Figure 3B)

| ID | Sequence, 5'-3' | Concentration, nM |
| --- | --- | --- |
| S | [Cy3]AAAGAGTAAAGTGACGGGC | 50 |
| Q | CACCTACGCTTTACTCTAT[BHQ2] | 200 |
| I | See table below | 800 |

The following table lists affinity between I and Q as concentration of the duplex C_AB_ (μM) when oligos are mixed at 1 μM each. Affinity is either assessed from experiment or predicted by NUPACK.

| I sequence, 5'-3' | Experimental  affinity  at 25 °C | NUPACK  affinity  at 25 °C | Experimental  affinity  at 50 °C | NUPACK  affinity  at 50 °C |
| --- | --- | --- | --- | --- |
| TCAGAGTAAAGCGTGTGTA | 0.88 | 0.99975 | 0.62 | 0.93409 |
| ATAGAGTAAAGCGGAATAT | 0.90 | 0.99974 | 0.66 | 0.91294 |
| ATAGAGTAAAGCGATAAAG | 0.85 | 0.99974 | 0.62 | 0.90610 |
| TTAGAGTAAAGCGAGTGTT | 0.86 | 0.99973 | 0.49 | 0.90153 |
| TGAGAGTAAAGCGTCGATT | 0.79 | 0.99972 | 0.45 | 0.93687 |
| AGGGAGTAAAGCGTGGATA | 0.89 | 0.99972 | 0.67 | 0.87406 |
| GCAGAGTAAAGCGTATTAG | 0.87 | 0.99971 | 0.60 | 0.93357 |
| ATAGAGTAAAGCGACCATT | 0.81 | 0.99969 | 0.39 | 0.91016 |
| ATAGAGTAAAAGCGAGGTA | 0.93 | 0.99969 | 0.67 | 0.66933 |
| CTGGAGTAAAGCGTGGATT | 0.89 | 0.99968 | 0.64 | 0.86682 |
| GTGGAGTAAAGCGAGGGTA | 0.83 | 0.99967 | 0.60 | 0.85715 |
| TTAGAGTAAAGCGAAAATT | 0.66 | 0.99966 | 0.37 | 0.90454 |
| AATGAGTAAAGCGAGGATG | 0.79 | 0.99964 | 0.49 | 0.80298 |
| GAAGAGTAAAGCGTTCGTT | 0.82 | 0.99963 | 0.59 | 0.92359 |
| GCAGAGTAAAGCGGTGGGG | 0.89 | 0.99963 | 0.59 | 0.88837 |
| CTAGAGTAAAGCGGGATTG | 0.81 | 0.99962 | 0.34 | 0.88221 |
| GAGGTGTAAAGGGTAGGTG | 0.92 | 0.99959 | 0.66 | 0.81012 |
| TCAGTGTAAAGCGTTGGTA | 0.90 | 0.99959 | 0.59 | 0.88148 |
| GTAGAGTAATGCGTTGGAA | 0.89 | 0.99958 | 0.45 | 0.78540 |
| GGAGTGTAAAGGGTAGGTT | 0.89 | 0.99957 | 0.47 | 0.83694 |
| AGAGAGTAAAGCGGGAATG | 0.82 | 0.99957 | 0.42 | 0.88697 |
| TTTGAGTAAAGGGTAGGCA | 0.91 | 0.99956 | 0.61 | 0.82399 |
| TTAGAGTAAAGCGACCATG | 0.65 | 0.99955 | 0.16 | 0.89561 |
| GTAGAGTAAAGCGAATTGG | 0.80 | 0.99954 | 0.27 | 0.88657 |
| AAAGAGTAAAGGGTAGAAC | 0.87 | 0.99953 | 0.43 | 0.82496 |
| AAAGAGGAAAGCGTTGGGT | 0.92 | 0.99953 | 0.65 | 0.90185 |
| GTGGAGTAAAGCGTAAATC | 0.78 | 0.99953 | 0.54 | 0.88068 |
| TGAGAGTAAAGCGAGCGTG | 0.64 | 0.99952 | 0.27 | 0.89676 |
| GTGGAGTAAAGCGTGTGTC | 0.89 | 0.99952 | 0.58 | 0.86093 |
| TTGGAGTAAAGCGTATATT | 0.85 | 0.99950 | 0.61 | 0.86693 |
| TTGGAGTAAAGCGGGGGTC | 0.87 | 0.99948 | 0.47 | 0.82851 |
| CTAGTGTAAAGCGTTGGGA | 0.89 | 0.99948 | 0.52 | 0.78819 |
| CTAGAGAAAAGCGTTGGAT | 0.85 | 0.99948 | 0.32 | 0.84854 |
| AGAGAGTAAAGCAGGTGTG | 0.90 | 0.99947 | 0.48 | 0.81452 |
| AAAGAGTTAAGCGTGGTGG | 0.87 | 0.99947 | 0.47 | 0.76478 |
| ATTGAGTAAAGCGTAAAAC | 0.85 | 0.99946 | 0.55 | 0.87333 |
| GGTGAGTAAAGGGTTGGTG | 0.90 | 0.99945 | 0.42 | 0.74641 |
| CTAGAGTAAAGCGGACATC | 0.82 | 0.99944 | 0.52 | 0.88174 |
| AGAGAGTAAAGCGAATGAC | 0.57 | 0.99943 | 0.12 | 0.89242 |
| CAAGAGTAAAGCGAACGGG | 0.72 | 0.99943 | 0.15 | 0.89098 |
| TAAGAGTAAAGCATAGGAC | 0.85 | 0.99942 | 0.37 | 0.85810 |
| CTGGAGTAAAGAGTAGGAC | 0.84 | 0.99941 | 0.41 | 0.78648 |
| ATAGAGTAAAGGGATAGGG | 0.93 | 0.99941 | 0.51 | 0.55584 |
| CTAGAGGAAAGCGTGGATA | 0.88 | 0.99940 | 0.50 | 0.86036 |
| ATGGAGTAAAGCGGGGGGA | 0.80 | 0.99939 | 0.42 | 0.79678 |
| TAAGAGAAAAGGGTAGGTC | 0.84 | 0.99938 | 0.22 | 0.85573 |
| CAAGAGGAAAGCGTGGGTA | 0.91 | 0.99937 | 0.61 | 0.90223 |
| GAAGAGTAAAGCGGATGAG | 0.80 | 0.99937 | 0.33 | 0.88140 |
| ATAGGGTAAAGCGAGGTGG | 0.94 | 0.99934 | 0.67 | 0.65844 |
| TTGCGGTAAAGGGTAGGTG | 0.89 | 0.99932 | 0.50 | 0.77118 |
| AAGATGTAAAGCGTCGGTG | 0.88 | 0.99930 | 0.45 | 0.76674 |
| AGAGTATAAAGCGTTGGTA | 0.81 | 0.99930 | 0.15 | 0.81596 |
| GGAGAGTAAAGGGTAGTTC | 0.87 | 0.99930 | 0.30 | 0.77387 |
| AGAGAGAAAAGCGTTGGAA | 0.88 | 0.99928 | 0.40 | 0.84555 |
| ATAGAGGAAAGCGATGGAT | 0.88 | 0.99927 | 0.50 | 0.82222 |
| ATAGAGGAAAGCGTACAGA | 0.91 | 0.99926 | 0.59 | 0.88364 |
| TGGGAGTAAAGCGTCGTTG | 0.84 | 0.99925 | 0.50 | 0.85019 |
| TTAGAGTAAAGAGTAGAGA | 0.84 | 0.99925 | 0.28 | 0.73390 |
| TGGGAGTAATGGGTAGGTG | 0.87 | 0.99923 | 0.21 | 0.61236 |
| GATAAGTAAAGAGTAGGTG | 0.90 | 0.99922 | 0.38 | 0.80474 |
| TTTGAGGAAAGCGTTGGTC | 0.90 | 0.99920 | 0.51 | 0.87771 |
| TTAGAGTAAAGGGCAGGAA | 0.80 | 0.99916 | 0.06 | 0.73106 |
| ATAGAGGAAAGCGGGGATA | 0.89 | 0.99915 | 0.46 | 0.80554 |
| AGAGAGTAAAGCGGTCGTT | 0.68 | 0.99914 | 0.18 | 0.85439 |
| AGGAAGTAAAGGGTAGGGG | 0.88 | 0.99914 | 0.48 | 0.76946 |
| ATAGAGTAAAGCGGCAGGA | 0.92 | 0.99996 | 0.57 | 0.95230 |
| TGTGAGTAAAGCGTTGGTC | 0.92 | 0.99992 | 0.61 | 0.95843 |
| ACAGAGTAAAGCGAGGTAG | 0.85 | 0.99990 | 0.56 | 0.93999 |
| GGAGAGTAAAGCGTTGTGG | 0.93 | 0.99989 | 0.71 | 0.94370 |
| TTAGAGTAAAGCGTAAAAC | 0.89 | 0.99988 | 0.59 | 0.95303 |
| GTAGAGTAAAGCGGGGGGA | 0.84 | 0.99986 | 0.39 | 0.91965 |
| CTAGAGTAAAGCGTGATGG | 0.91 | 0.99986 | 0.63 | 0.94302 |
| ATAGAGTAAAGCGTCCCCA | 0.92 | 0.99985 | 0.67 | 0.94671 |
| ACTGAGTAAAGCGTTGGAT | 0.88 | 0.99985 | 0.45 | 0.92442 |
| CTAGAGTAAAGCGTATCAA | 0.89 | 0.99985 | 0.49 | 0.94701 |
| GACGAGTAAAGGGTAGGTT | 0.90 | 0.99985 | 0.55 | 0.92162 |
| AGAGAGTAAAGCGTGTGAT | 0.92 | 0.99985 | 0.70 | 0.94213 |
| TCAGAGTAAAGCGTAAATA | 0.84 | 0.99983 | 0.49 | 0.94774 |
| GCAGAGTAAAGCGTGGATC | 0.88 | 0.99982 | 0.53 | 0.93723 |
| AAAGAGTAAAGCGAGGGAC | 0.86 | 0.99982 | 0.44 | 0.91807 |
| AGAGAGTAAAGCGTGATAG | 0.87 | 0.99981 | 0.55 | 0.94196 |
| GCAGAGTAAAGAGTAGGAA | 0.87 | 0.99981 | 0.29 | 0.90773 |
| GCTGAGTAAAGGGTAGGAG | 0.90 | 0.99980 | 0.52 | 0.88529 |
| CCAGAGTAAAGCGTATGAC | 0.86 | 0.99980 | 0.47 | 0.94112 |
| GAAGAGTAAAGCGTGAATG | 0.84 | 0.99979 | 0.58 | 0.93776 |
| CTAGAGTAAAGCGTCGAGA | 0.87 | 0.99979 | 0.53 | 0.94173 |
| AATGAGTAAAGCGTCGGAC | 0.89 | 0.99979 | 0.52 | 0.90687 |
| GTAGAGTAAAGCGTCCATT | 0.86 | 0.99978 | 0.54 | 0.93590 |
| CAAGAGTAAAGAGTAGGGT | 0.90 | 0.99977 | 0.54 | 0.88199 |
| GGAGAGTAAAGCGTGCGAG | 0.79 | 0.99977 | 0.52 | 0.93906 |
| GTAGAGTAAAGCGTTAAAG | 0.90 | 0.99977 | 0.64 | 0.93254 |
| TGGGAGTAAAGGGTAGGCG | 0.90 | 0.99976 | 0.48 | 0.85413 |
| CGAGAGTAAAGCGGGGTTG | 0.87 | 0.99976 | 0.58 | 0.92020 |
| TTGGAGTAAAGCGTCGGGA | 0.92 | 0.99975 | 0.65 | 0.89581 |
| TTAGAGTAAAGCGGTGAGG | 0.91 | 0.99975 | 0.65 | 0.88884 |

#### Experimental temperature-independent YES-gate.

| ID | Sequence, 5'-3' | Concentration, μM | Hairpin formation energy at 25 °C, kcal/mol |
| --- | --- | --- | --- |
| I | GGAGAGTAAAGCGTTGTGG | 2.00 | 0.00 |
| C1 | TGGGAGTAATGGGTAGGTG | 1.40 | 0.00 |
| Q | CACCTACGCTTTACTCTAT[BHQ2] | 0.70 | 0.00 |
| S | [Cy3]AAAGAGTAAAGTGACGGGC | 0.24 | 0.00 |

Mutual affinity of the oligos. Concentration of the duplex C_AB_ (μM) when oligos are mixed at 1

μM each (at 25 °C), as predicted by NUPACK:

|  | I | C1 | Q | S |
| --- | --- | --- | --- | --- |
| I | 0.0033 | 0.0001 | 0.9999 | 0.0013 |
| C1 | 0.0001 | 0.0000 | 0.9992 | 0.0002 |
| Q | 0.9999 | 0.9992 | 0.0006 | 0.9691 |
| S | 0.0013 | 0.0002 | 0.9691 | 0.0012 |

Mutual affinity of the oligos. Concentration of the duplex C_AB_ (μM) when oligos are mixed at 1

μM each (at 50 °C), as predicted by NUPACK:

|  | I | C1 | Q | S |
| --- | --- | --- | --- | --- |
| I | 0.00 | 0.00 | 0.94 | 0.00 |
| C1 | 0.00 | 0.00 | 0.61 | 0.00 |
| Q | 0.94 | 0.61 | 0.00 | 0.14 |
| S | 0.00 | 0.00 | 0.14 | 0.00 |

Mutual affinity of the oligos. Approximate dissociation constant K_d_ at 25 °C in M, as predicted by NUPACK:

|  | I | C1 | Q | S |
| --- | --- | --- | --- | --- |
| I | 3.03E-04 | 1.53E-02 | 1.15E-14 | 7.94E-04 |
| C1 | 1.53E-02 | 3.11E-02 | 5.96E-13 | 5.37E-03 |
| Q | 1.15E-14 | 5.96E-13 | 1.63E-03 | 9.86E-10 |
| S | 7.94E-04 | 5.37E-03 | 9.86E-10 | 8.13E-04 |

Mutual affinity of the oligos. Approximate dissociation constant K_d_ at 50 °C in M, as predicted by NUPACK:

|  | I | C1 | Q | S |
| --- | --- | --- | --- | --- |
| I | 5.78E-03 | 2.23E-02 | 3.36E-09 | 3.66E-03 |
| C1 | 2.23E-02 | 3.50E-02 | 2.45E-07 | 1.04E-02 |
| Q | 3.36E-09 | 2.45E-07 | 7.63E-03 | 5.47E-06 |
| S | 3.66E-03 | 1.04E-02 | 5.47E-06 | 4.57E-03 |

Below we give affinities for the same oligos, approximately calculated from experiment. Values in blank cells were not measured.

Mutual affinity of the oligos. Concentration of the duplex C_AB_ (μM) when oligos are mixed at 1

μM each (at 25 °C), as measured experimentally:

|  | I | C1 | Q | S |
| --- | --- | --- | --- | --- |
| I |  |  | 0.93 |  |
| C1 |  |  | 0.87 |  |
| Q | 0.93 | 0.87 |  | 0.91 |
| S |  |  | 0.91 |  |

Mutual affinity of the oligos. Concentration of the duplex C_AB_ (μM) when oligos are mixed at 1

μM each (at 50 °C), as measured experimentally:

|  | I | C1 | Q | S |
| --- | --- | --- | --- | --- |
| I |  |  | 0.71 |  |
| C1 |  |  | 0.21 |  |
| Q | 0.71 | 0.21 |  | 0.59 |
| S |  |  | 0.59 |  |

Mutual affinity of the oligos. Approximate dissociation constant K_d_ at 25 °C in M, as measured experimentally:

|  | I | C1 | Q | S |
| --- | --- | --- | --- | --- |
| I |  |  | 5.27E-09 |  |
| C1 |  |  | 1.94E-08 |  |
| Q | 5.27E-09 | 1.94E-08 |  | 8.90E-09 |
| S |  |  | 8.90E-09 |  |

Mutual affinity of the oligos. Approximate dissociation constant K_d_ at 50 °C in M, as measured experimentally:

|  | I | C1 | Q | S |
| --- | --- | --- | --- | --- |
| I |  |  | 1.18E-07 |  |
| C1 |  |  | 2.97E-06 |  |
| Q | 1.18E-07 | 2.97E-06 |  | 2.85E-07 |
| S |  |  | 2.85E-07 |  |

#### Salt-independent gates: sodium

| ID | Sequence, 5'-3' | Concentration, μM | Hairpin formation energy at 25 °C and 1M Na^+^, kcal/mol |
| --- | --- | --- | --- |
| I | TTAATTCACGGTTTCTAT | 5.42 | 0.00 |
| Q | ATTGTAAACGTGAAGTAA | 0.99 | -0.97 |
| S | GTAATTCACGTCTACACT | 0.38 | 0.00 |
| C | TTAATTCACTTTTACCAT | 2.48 | 0.00 |

Mutual affinity of the oligos. Concentration of the duplex C_AB_ (μM) when oligos are mixed at 1

μM each (at 25 °C, 1M Na^+^), as predicted by NUPACK:

|  | I | Q | S | C |
| --- | --- | --- | --- | --- |
| I | 0.00 | 0.90 | 0.00 | 0.00 |
| Q | 0.90 | 0.00 | 0.92 | 0.86 |
| S | 0.00 | 0.92 | 0.00 | 0.00 |
| C | 0.00 | 0.86 | 0.00 | 0.00 |

Mutual affinity of the oligos. Concentration of the duplex C_AB_ (μM) when oligos are mixed at 1

μM each (at 25 °C, 0.1M Na^+^), as predicted by NUPACK:

|  | I | Q | S | C |
| --- | --- | --- | --- | --- |
| I | 0.00 | 0.62 | 0.00 | 0.00 |
| Q | 0.62 | 0.00 | 0.66 | 0.38 |
| S | 0.00 | 0.66 | 0.00 | 0.00 |
| C | 0.00 | 0.38 | 0.00 | 0.00 |

Mutual affinity of the oligos. Approximate dissociation constant K_d_ (at 25 °C, 1M Na^+^) in M, as predicted by NUPACK:

|  | I | Q | S | C |
| --- | --- | --- | --- | --- |
| I | 4.62E-04 | 1.01E-08 | 5.49E-04 | 3.52E-04 |
| Q | 1.01E-08 | 5.86E-04 | 6.73E-09 | 2.27E-08 |
| S | 5.49E-04 | 6.73E-09 | 3.13E-04 | 1.39E-03 |
| C | 3.52E-04 | 2.27E-08 | 1.39E-03 | 3.00E-03 |

Mutual affinity of the oligos. Approximate dissociation constant K_d_ (at 25 °C, 0.1M Na^+^) in M, as predicted by NUPACK:

|  | I | Q | S | C |
| --- | --- | --- | --- | --- |
| I | 1.03E-03 | 2.29E-07 | 9.97E-04 | 7.95E-04 |
| Q | 2.29E-07 | 8.75E-04 | 1.70E-07 | 9.87E-07 |
| S | 9.97E-04 | 1.70E-07 | 6.20E-04 | 3.52E-03 |
| C | 7.95E-04 | 9.87E-07 | 3.52E-03 | 5.46E-03 |

#### Salt-independent gates: magnesium

| ID | Sequence, 5'-3' | Concentration, μM | Hairpin formation energy at 25 °C, kcal/mol |
| --- | --- | --- | --- |
| I | TTAATTCACGGTTTCTAT | 9.29 | 0.00 |
| Q | ATTGTAAACGTGAATTTA | 0.55 | -0.98 |
| S | GTAATTCACGTCTACACT | 0.34 | 0.00 |
| M | TTAATTCACTTTTACCAT | 2.35 | 0.00 |

Mutual affinity of the oligos. Concentration of the duplex C_AB_ (μM) when oligos are mixed at 1

μM each (at 25 °C, 10 mM Mg^2+^, 0.05M Na^+^), as predicted by NUPACK:

|  | I | Q | S | M |
| --- | --- | --- | --- | --- |
| I | 0.00 | 0.94 | 0.00 | 0.00 |
| Q | 0.94 | 0.00 | 0.96 | 0.90 |
| S | 0.00 | 0.96 | 0.00 | 0.00 |
| M | 0.00 | 0.90 | 0.00 | 0.00 |

Mutual affinity of the oligos. Concentration of the duplex C_AB_ (μM) when oligos are mixed at 1

μM each (at 25 °C, 0 mM Mg^2+^, 0.05M Na^+^), as predicted by NUPACK:

|  | I | Q | S | M |
| --- | --- | --- | --- | --- |
| I | 0.00 | 0.80 | 0.00 | 0.00 |
| Q | 0.80 | 0.00 | 0.84 | 0.60 |
| S | 0.00 | 0.84 | 0.00 | 0.00 |
| M | 0.00 | 0.60 | 0.00 | 0.00 |

Mutual affinity of the oligos. Approximate dissociation constant K_d_ (at 25 °C, 10 mM Mg^2+^, 0.05M Na^+^) in M, as predicted by NUPACK:

|  | I | Q | S | M |
| --- | --- | --- | --- | --- |
| I | 6.52E-04 | 3.45E-09 | 7.06E-04 | 4.91E-04 |
| Q | 3.45E-09 | 6.36E-04 | 1.46E-09 | 9.98E-09 |
| S | 7.06E-04 | 1.46E-09 | 4.18E-04 | 2.07E-03 |
| M | 4.91E-04 | 9.98E-09 | 2.07E-03 | 3.78E-03 |

Mutual affinity of the oligos. Approximate dissociation constant K_d_ (at 25 °C, 0 mM Mg^2+^, 0.05M Na^+^) in M, as predicted by NUPACK:

|  | I | Q | S | M |
| --- | --- | --- | --- | --- |
| I | 1.30E-03 | 5.02E-08 | 1.20E-03 | 1.03E-03 |
| Q | 5.02E-08 | 7.16E-04 | 3.02E-08 | 2.72E-07 |
| S | 1.20E-03 | 3.02E-08 | 7.63E-04 | 4.60E-03 |
| M | 1.03E-03 | 2.72E-07 | 4.60E-03 | 6.71E-03 |

#### Receptor-activator networks

Please note, that parameters for receptor-activator networks with 100 receptors (Figure 5D) are provided in a separate file “Supplementary Data Figure 5D.xlsx”.

N = 1 case (1 receptor, 1 activator)

Mutual affinity of the oligos. Concentration of the duplex C_AB_ (μM) when oligos are mixed at 1

μM each (25 °C).

|  | R1 | A1 | C1 |
| --- | --- | --- | --- |
| R1 | 0.000 | 0.996 | 0.000 |
| A1 | 0.996 | 0.000 | 0.996 |
| C1 | 0.000 | 0.996 | 0.000 |

Mutual affinity of the oligos. Concentration of the duplex C_AB_ (μM) when oligos are mixed at 1

μM each (50 °C).

|  | R1 | A1 | C1 |
| --- | --- | --- | --- |
| R1 | 0.00 | 0.60 | 0.00 |
| A1 | 0.60 | 0.00 | 0.21 |
| C1 | 0.00 | 0.21 | 0.00 |

Mutual affinity of the oligos. Approximate dissociation constant K_d_ (25 °C) in M.

|  | R1 | A1 | C1 |
| --- | --- | --- | --- |
| R1 | 1.00E+00 | 1.55E-11 | 1.00E+00 |
| A1 | 1.55E-11 | 1.00E+00 | 1.54E-11 |
| C1 | 1.00E+00 | 1.54E-11 | 1.00E+00 |

Mutual affinity of the oligos. Approximate dissociation constant K_d_ (50 °C) in M.

|  | R1 | A1 | C1 |
| --- | --- | --- | --- |
| R1 | 1.00E+00 | 2.67E-07 | 1.00E+00 |
| A1 | 2.67E-07 | 1.00E+00 | 2.89E-06 |
| C1 | 1.00E+00 | 2.89E-06 | 1.00E+00 |

N = 2 case (2 receptors, 2 activators)

Mutual affinity of the oligos. Concentration of the duplex C_AB_ (μM) when oligos are mixed at 1

μM each (25 °C).

|  | R1 | R2 | A1 | A2 | C1 | C2 |
| --- | --- | --- | --- | --- | --- | --- |
| R1 | 0.000 | 0.000 | 0.992 | 0.990 | 0.000 | 0.000 |
| R2 | 0.000 | 0.000 | 0.995 | 0.996 | 0.000 | 0.000 |
| A1 | 0.992 | 0.995 | 0.000 | 0.000 | 0.995 | 0.000 |
| A2 | 0.990 | 0.996 | 0.000 | 0.000 | 0.000 | 0.997 |
| C1 | 0.000 | 0.000 | 0.995 | 0.000 | 0.000 | 0.000 |
| C2 | 0.000 | 0.000 | 0.000 | 0.997 | 0.000 | 0.000 |

Mutual affinity of the oligos. Concentration of the duplex C_AB_ (μM) when oligos are mixed at 1

μM each (50 °C).

|  | R1 | R2 | A1 | A2 | C1 | C2 |
| --- | --- | --- | --- | --- | --- | --- |
| R1 | 0.00 | 0.00 | 0.39 | 0.35 | 0.00 | 0.00 |
| R2 | 0.00 | 0.00 | 0.53 | 0.60 | 0.00 | 0.00 |
| A1 | 0.39 | 0.53 | 0.00 | 0.00 | 0.17 | 0.00 |
| A2 | 0.35 | 0.60 | 0.00 | 0.00 | 0.00 | 0.30 |
| C1 | 0.00 | 0.00 | 0.17 | 0.00 | 0.00 | 0.00 |
| C2 | 0.00 | 0.00 | 0.00 | 0.30 | 0.00 | 0.00 |

Mutual affinity of the oligos. Approximate dissociation constant K_d_ (25 °C) in M.

|  | R1 | R2 | A1 | A2 | C1 | C2 |
| --- | --- | --- | --- | --- | --- | --- |
| R1 | 1.00E+00 | 1.00E+00 | 7.15E-11 | 1.06E-10 | 1.00E+00 | 1.00E+00 |
| R2 | 1.00E+00 | 1.00E+00 | 2.67E-11 | 1.55E-11 | 1.00E+00 | 1.00E+00 |
| A1 | 7.15E-11 | 2.67E-11 | 1.00E+00 | 1.00E+00 | 2.39E-11 | 1.00E+00 |
| A2 | 1.06E-10 | 1.55E-11 | 1.00E+00 | 1.00E+00 | 1.00E+00 | 7.47E-12 |
| C1 | 1.00E+00 | 1.00E+00 | 2.39E-11 | 1.00E+00 | 1.00E+00 | 1.00E+00 |
| C2 | 1.00E+00 | 1.00E+00 | 1.00E+00 | 7.47E-12 | 1.00E+00 | 1.00E+00 |

Mutual affinity of the oligos. Approximate dissociation constant K_d_ (50 °C) in M.

|  | R1 | R2 | A1 | A2 | C1 | C2 |
| --- | --- | --- | --- | --- | --- | --- |
| R1 | 1.00E+00 | 1.00E+00 | 9.66E-07 | 1.20E-06 | 1.00E+00 | 1.00E+00 |
| R2 | 1.00E+00 | 1.00E+00 | 4.04E-07 | 2.67E-07 | 1.00E+00 | 1.00E+00 |
| A1 | 9.66E-07 | 4.04E-07 | 1.00E+00 | 1.00E+00 | 4.17E-06 | 1.00E+00 |
| A2 | 1.20E-06 | 2.67E-07 | 1.00E+00 | 1.00E+00 | 1.00E+00 | 1.63E-06 |
| C1 | 1.00E+00 | 1.00E+00 | 4.17E-06 | 1.00E+00 | 1.00E+00 | 1.00E+00 |
| C2 | 1.00E+00 | 1.00E+00 | 1.00E+00 | 1.63E-06 | 1.00E+00 | 1.00E+00 |

N = 3 case (3 receptors, 3 activators)

Mutual affinity of the oligos. Concentration of the duplex C_AB_ (μM) when oligos are mixed at 1

μM each (25 °C).

|  | R1 | R2 | R3 | A1 | A2 | A3 | C1 | C2 | C3 | D1 | D2 | D3 |
| --- | --- | --- | --- | --- | --- | --- | --- | --- | --- | --- | --- | --- |
| R1 | 0.000 | 0.000 | 0.000 | 0.955 | 0.954 | 0.964 | 0.000 | 0.000 | 0.000 | 0.996 | 0.000 | 0.000 |
| R2 | 0.000 | 0.000 | 0.000 | 0.981 | 0.994 | 0.972 | 0.000 | 0.000 | 0.000 | 0.000 | 0.911 | 0.000 |
| R3 | 0.000 | 0.000 | 0.000 | 0.996 | 0.996 | 0.993 | 0.000 | 0.000 | 0.000 | 0.000 | 0.000 | 0.924 |
| A1 | 0.955 | 0.981 | 0.996 | 0.000 | 0.000 | 0.000 | 0.997 | 0.000 | 0.000 | 0.000 | 0.000 | 0.000 |
| A2 | 0.954 | 0.994 | 0.996 | 0.000 | 0.000 | 0.000 | 0.000 | 0.985 | 0.000 | 0.000 | 0.000 | 0.000 |
| A3 | 0.964 | 0.972 | 0.993 | 0.000 | 0.000 | 0.000 | 0.000 | 0.000 | 0.900 | 0.000 | 0.000 | 0.000 |
| C1 | 0.000 | 0.000 | 0.000 | 0.997 | 0.000 | 0.000 | 0.000 | 0.000 | 0.000 | 0.000 | 0.000 | 0.000 |
| C2 | 0.000 | 0.000 | 0.000 | 0.000 | 0.985 | 0.000 | 0.000 | 0.000 | 0.000 | 0.000 | 0.000 | 0.000 |
| C3 | 0.000 | 0.000 | 0.000 | 0.000 | 0.000 | 0.900 | 0.000 | 0.000 | 0.000 | 0.000 | 0.000 | 0.000 |
| D1 | 0.996 | 0.000 | 0.000 | 0.000 | 0.000 | 0.000 | 0.000 | 0.000 | 0.000 | 0.000 | 0.000 | 0.000 |
| D2 | 0.000 | 0.911 | 0.000 | 0.000 | 0.000 | 0.000 | 0.000 | 0.000 | 0.000 | 0.000 | 0.000 | 0.000 |
| D3 | 0.000 | 0.000 | 0.924 | 0.000 | 0.000 | 0.000 | 0.000 | 0.000 | 0.000 | 0.000 | 0.000 | 0.000 |

Mutual affinity of the oligos. Concentration of the duplex C_AB_ (μM) when oligos are mixed at 1

μM each (50 °C).

|  | R1 | R2 | R3 | A1 | A2 | A3 | C1 | C2 | C3 | D1 | D2 | D3 |
| --- | --- | --- | --- | --- | --- | --- | --- | --- | --- | --- | --- | --- |
| R1 | 0.000 | 0.000 | 0.000 | 0.103 | 0.100 | 0.143 | 0.000 | 0.000 | 0.000 | 0.238 | 0.000 | 0.000 |
| R2 | 0.000 | 0.000 | 0.000 | 0.254 | 0.488 | 0.197 | 0.000 | 0.000 | 0.000 | 0.000 | 0.011 | 0.000 |
| R3 | 0.000 | 0.000 | 0.000 | 0.600 | 0.600 | 0.432 | 0.000 | 0.000 | 0.000 | 0.000 | 0.000 | 0.012 |
| A1 | 0.103 | 0.254 | 0.600 | 0.000 | 0.000 | 0.000 | 0.293 | 0.000 | 0.000 | 0.000 | 0.000 | 0.000 |
| A2 | 0.100 | 0.488 | 0.600 | 0.000 | 0.000 | 0.000 | 0.000 | 0.032 | 0.000 | 0.000 | 0.000 | 0.000 |
| A3 | 0.143 | 0.197 | 0.432 | 0.000 | 0.000 | 0.000 | 0.000 | 0.000 | 0.010 | 0.000 | 0.000 | 0.000 |
| C1 | 0.000 | 0.000 | 0.000 | 0.293 | 0.000 | 0.000 | 0.000 | 0.000 | 0.000 | 0.000 | 0.000 | 0.000 |
| C2 | 0.000 | 0.000 | 0.000 | 0.000 | 0.032 | 0.000 | 0.000 | 0.000 | 0.000 | 0.000 | 0.000 | 0.000 |
| C3 | 0.000 | 0.000 | 0.000 | 0.000 | 0.000 | 0.010 | 0.000 | 0.000 | 0.000 | 0.000 | 0.000 | 0.000 |
| D1 | 0.238 | 0.000 | 0.000 | 0.000 | 0.000 | 0.000 | 0.000 | 0.000 | 0.000 | 0.000 | 0.000 | 0.000 |
| D2 | 0.000 | 0.011 | 0.000 | 0.000 | 0.000 | 0.000 | 0.000 | 0.000 | 0.000 | 0.000 | 0.000 | 0.000 |
| D3 | 0.000 | 0.000 | 0.012 | 0.000 | 0.000 | 0.000 | 0.000 | 0.000 | 0.000 | 0.000 | 0.000 | 0.000 |

Mutual affinity of the oligos. Approximate dissociation constant K_d_ (25 °C) in M.

|  | R1 | R2 | R3 | A1 | A2 | A3 | C1 | C2 | C3 | D1 | D2 | D3 |
| --- | --- | --- | --- | --- | --- | --- | --- | --- | --- | --- | --- | --- |
| R1 | 1E+00 | 1E+00 | 1E+00 | 2E-09 | 2E-09 | 1E-09 | 1E+00 | 1E+00 | 1E+00 | 1E-11 | 1E+00 | 1E+00 |
| R2 | 1E+00 | 1E+00 | 1E+00 | 4E-10 | 4E-11 | 8E-10 | 1E+00 | 1E+00 | 1E+00 | 1E+00 | 9E-09 | 1E+00 |
| R3 | 1E+00 | 1E+00 | 1E+00 | 2E-11 | 2E-11 | 5E-11 | 1E+00 | 1E+00 | 1E+00 | 1E+00 | 1E+00 | 6E-09 |
| A1 | 2E-09 | 4E-10 | 2E-11 | 1E+00 | 1E+00 | 1E+00 | 8E-12 | 1E+00 | 1E+00 | 1E+00 | 1E+00 | 1E+00 |
| A2 | 2E-09 | 4E-11 | 2E-11 | 1E+00 | 1E+00 | 1E+00 | 1E+00 | 2E-10 | 1E+00 | 1E+00 | 1E+00 | 1E+00 |
| A3 | 1E-09 | 8E-10 | 5E-11 | 1E+00 | 1E+00 | 1E+00 | 1E+00 | 1E+00 | 1E-08 | 1E+00 | 1E+00 | 1E+00 |
| C1 | 1E+00 | 1E+00 | 1E+00 | 8E-12 | 1E+00 | 1E+00 | 1E+00 | 1E+00 | 1E+00 | 1E+00 | 1E+00 | 1E+00 |
| C2 | 1E+00 | 1E+00 | 1E+00 | 1E+00 | 2E-10 | 1E+00 | 1E+00 | 1E+00 | 1E+00 | 1E+00 | 1E+00 | 1E+00 |
| C3 | 1E+00 | 1E+00 | 1E+00 | 1E+00 | 1E+00 | 1E-08 | 1E+00 | 1E+00 | 1E+00 | 1E+00 | 1E+00 | 1E+00 |
| D1 | 1E-11 | 1E+00 | 1E+00 | 1E+00 | 1E+00 | 1E+00 | 1E+00 | 1E+00 | 1E+00 | 1E+00 | 1E+00 | 1E+00 |
| D2 | 1E+00 | 9E-09 | 1E+00 | 1E+00 | 1E+00 | 1E+00 | 1E+00 | 1E+00 | 1E+00 | 1E+00 | 1E+00 | 1E+00 |
| D3 | 1E+00 | 1E+00 | 6E-09 | 1E+00 | 1E+00 | 1E+00 | 1E+00 | 1E+00 | 1E+00 | 1E+00 | 1E+00 | 1E+00 |

Mutual affinity of the oligos. Approximate dissociation constant K_d_ (50 °C) in M.

|  | R1 | R2 | R3 | A1 | A2 | A3 | C1 | C2 | C3 | D1 | D2 | D3 |
| --- | --- | --- | --- | --- | --- | --- | --- | --- | --- | --- | --- | --- |
| R1 | 1E+00 | 1E+00 | 1E+00 | 8E-06 | 8E-06 | 5E-06 | 1E+00 | 1E+00 | 1E+00 | 2E-06 | 1E+00 | 1E+00 |
| R2 | 1E+00 | 1E+00 | 1E+00 | 2E-06 | 5E-07 | 3E-06 | 1E+00 | 1E+00 | 1E+00 | 1E+00 | 9E-05 | 1E+00 |
| R3 | 1E+00 | 1E+00 | 1E+00 | 3E-07 | 3E-07 | 7E-07 | 1E+00 | 1E+00 | 1E+00 | 1E+00 | 1E+00 | 8E-05 |
| A1 | 8E-06 | 2E-06 | 3E-07 | 1E+00 | 1E+00 | 1E+00 | 2E-06 | 1E+00 | 1E+00 | 1E+00 | 1E+00 | 1E+00 |
| A2 | 8E-06 | 5E-07 | 3E-07 | 1E+00 | 1E+00 | 1E+00 | 1E+00 | 3E-05 | 1E+00 | 1E+00 | 1E+00 | 1E+00 |
| A3 | 5E-06 | 3E-06 | 7E-07 | 1E+00 | 1E+00 | 1E+00 | 1E+00 | 1E+00 | 1E-04 | 1E+00 | 1E+00 | 1E+00 |
| C1 | 1E+00 | 1E+00 | 1E+00 | 2E-06 | 1E+00 | 1E+00 | 1E+00 | 1E+00 | 1E+00 | 1E+00 | 1E+00 | 1E+00 |
| C2 | 1E+00 | 1E+00 | 1E+00 | 1E+00 | 3E-05 | 1E+00 | 1E+00 | 1E+00 | 1E+00 | 1E+00 | 1E+00 | 1E+00 |
| C3 | 1E+00 | 1E+00 | 1E+00 | 1E+00 | 1E+00 | 1E-04 | 1E+00 | 1E+00 | 1E+00 | 1E+00 | 1E+00 | 1E+00 |
| D1 | 2E-06 | 1E+00 | 1E+00 | 1E+00 | 1E+00 | 1E+00 | 1E+00 | 1E+00 | 1E+00 | 1E+00 | 1E+00 | 1E+00 |
| D2 | 1E+00 | 9E-05 | 1E+00 | 1E+00 | 1E+00 | 1E+00 | 1E+00 | 1E+00 | 1E+00 | 1E+00 | 1E+00 | 1E+00 |
| D3 | 1E+00 | 1E+00 | 8E-05 | 1E+00 | 1E+00 | 1E+00 | 1E+00 | 1E+00 | 1E+00 | 1E+00 | 1E+00 | 1E+00 |

### Algebraic systems

#### Line function – temperature independent system.

Concentrations of the participants.

| **ID** | **Concentration, μM** |
| --- | --- |
| I | 0.54 |
| M1 | 6.09 |
| M2 | 1.93 |
| M3 | 7.52 |
| M4 | 6.68 |
| M5 | 3.17 |
| M6 | 9.18 |
| M7 | 8.60 |
| S | 0.03 |
| C1 | 4.93 |
| C2 | 8.96 |
| C3 | 9.93 |
| C4 | 8.64 |
| C5 | 0.21 |
| C6 | 1.09 |
| C7 | 7.00 |
| C8 | 9.53 |
| C9 | 0.55 |

Mutual affinity of the participants. Concentration of the duplex C_AB_ (μM) when oligos are mixed at 1 μM each (25 °C).

|  | **I** | **M1** | **M2** | **M3** | **M4** | **M5** | **M6** | **M7** | **S** | **C1** | **C2** | **C3** | **C4** | **C5** | **C6** | **C7** | **C8** | **C9** |
| --- | --- | --- | --- | --- | --- | --- | --- | --- | --- | --- | --- | --- | --- | --- | --- | --- | --- | --- |
| **I** | 0.00 | 0.01 | 0.92 | 0.96 | 0.94 | 0.01 | 0.47 | 0.01 | 0.00 | 0.89 | 0.00 | 0.00 | 0.00 | 0.00 | 0.00 | 0.00 | 0.00 | 0.00 |
| **M1** | 0.01 | 0.00 | 0.98 | 0.74 | 0.94 | 0.99 | 0.40 | 0.66 | 0.21 | 0.00 | 0.97 | 0.00 | 0.00 | 0.00 | 0.00 | 0.00 | 0.00 | 0.00 |
| **M2** | 0.92 | 0.98 | 0.00 | 0.95 | 0.77 | 0.93 | 0.94 | 0.92 | 0.43 | 0.00 | 0.00 | 0.05 | 0.00 | 0.00 | 0.00 | 0.00 | 0.00 | 0.00 |
| **M3** | 0.96 | 0.74 | 0.95 | 0.00 | 0.48 | 0.12 | 0.00 | 0.98 | 0.54 | 0.00 | 0.00 | 0.00 | 0.97 | 0.00 | 0.00 | 0.00 | 0.00 | 0.00 |
| **M4** | 0.94 | 0.94 | 0.77 | 0.48 | 0.00 | 0.45 | 0.46 | 0.41 | 0.01 | 0.00 | 0.00 | 0.00 | 0.00 | 0.06 | 0.00 | 0.00 | 0.00 | 0.00 |
| **M5** | 0.01 | 0.99 | 0.93 | 0.12 | 0.45 | 0.00 | 0.87 | 0.11 | 0.13 | 0.00 | 0.00 | 0.00 | 0.00 | 0.00 | 0.92 | 0.00 | 0.00 | 0.00 |
| **M6** | 0.47 | 0.40 | 0.94 | 0.00 | 0.46 | 0.87 | 0.00 | 0.97 | 0.86 | 0.00 | 0.00 | 0.00 | 0.00 | 0.00 | 0.00 | 0.99 | 0.00 | 0.00 |
| **M7** | 0.01 | 0.66 | 0.92 | 0.98 | 0.41 | 0.11 | 0.97 | 0.00 | 0.02 | 0.00 | 0.00 | 0.00 | 0.00 | 0.00 | 0.00 | 0.00 | 0.10 | 0.00 |
| **S** | 0.00 | 0.21 | 0.43 | 0.54 | 0.01 | 0.13 | 0.86 | 0.02 | 0.00 | 0.00 | 0.00 | 0.00 | 0.00 | 0.00 | 0.00 | 0.00 | 0.00 | 0.25 |
| **C1** | 0.89 | 0.00 | 0.00 | 0.00 | 0.00 | 0.00 | 0.00 | 0.00 | 0.00 | 0.00 | 0.00 | 0.00 | 0.00 | 0.00 | 0.00 | 0.00 | 0.00 | 0.00 |
| **C2** | 0.00 | 0.97 | 0.00 | 0.00 | 0.00 | 0.00 | 0.00 | 0.00 | 0.00 | 0.00 | 0.00 | 0.00 | 0.00 | 0.00 | 0.00 | 0.00 | 0.00 | 0.00 |
| **C3** | 0.00 | 0.00 | 0.05 | 0.00 | 0.00 | 0.00 | 0.00 | 0.00 | 0.00 | 0.00 | 0.00 | 0.00 | 0.00 | 0.00 | 0.00 | 0.00 | 0.00 | 0.00 |
| **C4** | 0.00 | 0.00 | 0.00 | 0.97 | 0.00 | 0.00 | 0.00 | 0.00 | 0.00 | 0.00 | 0.00 | 0.00 | 0.00 | 0.00 | 0.00 | 0.00 | 0.00 | 0.00 |
| **C5** | 0.00 | 0.00 | 0.00 | 0.00 | 0.06 | 0.00 | 0.00 | 0.00 | 0.00 | 0.00 | 0.00 | 0.00 | 0.00 | 0.00 | 0.00 | 0.00 | 0.00 | 0.00 |
| **C6** | 0.00 | 0.00 | 0.00 | 0.00 | 0.00 | 0.92 | 0.00 | 0.00 | 0.00 | 0.00 | 0.00 | 0.00 | 0.00 | 0.00 | 0.00 | 0.00 | 0.00 | 0.00 |
| **C7** | 0.00 | 0.00 | 0.00 | 0.00 | 0.00 | 0.00 | 0.99 | 0.00 | 0.00 | 0.00 | 0.00 | 0.00 | 0.00 | 0.00 | 0.00 | 0.00 | 0.00 | 0.00 |
| **C8** | 0.00 | 0.00 | 0.00 | 0.00 | 0.00 | 0.00 | 0.00 | 0.10 | 0.00 | 0.00 | 0.00 | 0.00 | 0.00 | 0.00 | 0.00 | 0.00 | 0.00 | 0.00 |
| **C9** | 0.00 | 0.00 | 0.00 | 0.00 | 0.00 | 0.00 | 0.00 | 0.00 | 0.25 | 0.00 | 0.00 | 0.00 | 0.00 | 0.00 | 0.00 | 0.00 | 0.00 | 0.00 |

Mutual affinity of the participants. Concentration of the duplex C_AB_ (μM) when oligos are mixed at 1 μM each (50 °C).

|  | **I** | **M1** | **M2** | **M3** | **M4** | **M5** | **M6** | **M7** | **S** | **C1** | **C2** | **C3** | **C4** | **C5** | **C6** | **C7** | **C8** | **C9** |
| --- | --- | --- | --- | --- | --- | --- | --- | --- | --- | --- | --- | --- | --- | --- | --- | --- | --- | --- |
| **I** | 0.00 | 0.00 | 0.31 | 0.46 | 0.37 | 0.00 | 0.04 | 0.00 | 0.00 | 0.01 | 0.00 | 0.00 | 0.00 | 0.00 | 0.00 | 0.00 | 0.00 | 0.00 |
| **M1** | 0.00 | 0.00 | 0.63 | 0.11 | 0.38 | 0.82 | 0.03 | 0.08 | 0.01 | 0.00 | 0.04 | 0.00 | 0.00 | 0.00 | 0.00 | 0.00 | 0.00 | 0.00 |
| **M2** | 0.31 | 0.63 | 0.00 | 0.41 | 0.12 | 0.35 | 0.40 | 0.33 | 0.03 | 0.00 | 0.00 | 0.00 | 0.00 | 0.00 | 0.00 | 0.00 | 0.00 | 0.00 |
| **M3** | 0.46 | 0.11 | 0.41 | 0.00 | 0.04 | 0.01 | 0.00 | 0.67 | 0.05 | 0.00 | 0.00 | 0.00 | 0.04 | 0.00 | 0.00 | 0.00 | 0.00 | 0.00 |
| **M4** | 0.37 | 0.38 | 0.12 | 0.04 | 0.00 | 0.04 | 0.04 | 0.03 | 0.00 | 0.00 | 0.00 | 0.00 | 0.00 | 0.00 | 0.00 | 0.00 | 0.00 | 0.00 |
| **M5** | 0.00 | 0.82 | 0.35 | 0.01 | 0.04 | 0.00 | 0.22 | 0.01 | 0.01 | 0.00 | 0.00 | 0.00 | 0.00 | 0.00 | 0.01 | 0.00 | 0.00 | 0.00 |
| **M6** | 0.04 | 0.03 | 0.40 | 0.00 | 0.04 | 0.22 | 0.00 | 0.53 | 0.20 | 0.00 | 0.00 | 0.00 | 0.00 | 0.00 | 0.00 | 0.09 | 0.00 | 0.00 |
| **M7** | 0.00 | 0.08 | 0.33 | 0.67 | 0.03 | 0.01 | 0.53 | 0.00 | 0.00 | 0.00 | 0.00 | 0.00 | 0.00 | 0.00 | 0.00 | 0.00 | 0.00 | 0.00 |
| **S** | 0.00 | 0.01 | 0.03 | 0.05 | 0.00 | 0.01 | 0.20 | 0.00 | 0.00 | 0.00 | 0.00 | 0.00 | 0.00 | 0.00 | 0.00 | 0.00 | 0.00 | 0.00 |
| **C1** | 0.01 | 0.00 | 0.00 | 0.00 | 0.00 | 0.00 | 0.00 | 0.00 | 0.00 | 0.00 | 0.00 | 0.00 | 0.00 | 0.00 | 0.00 | 0.00 | 0.00 | 0.00 |
| **C2** | 0.00 | 0.04 | 0.00 | 0.00 | 0.00 | 0.00 | 0.00 | 0.00 | 0.00 | 0.00 | 0.00 | 0.00 | 0.00 | 0.00 | 0.00 | 0.00 | 0.00 | 0.00 |
| **C3** | 0.00 | 0.00 | 0.00 | 0.00 | 0.00 | 0.00 | 0.00 | 0.00 | 0.00 | 0.00 | 0.00 | 0.00 | 0.00 | 0.00 | 0.00 | 0.00 | 0.00 | 0.00 |
| **C4** | 0.00 | 0.00 | 0.00 | 0.04 | 0.00 | 0.00 | 0.00 | 0.00 | 0.00 | 0.00 | 0.00 | 0.00 | 0.00 | 0.00 | 0.00 | 0.00 | 0.00 | 0.00 |
| **C5** | 0.00 | 0.00 | 0.00 | 0.00 | 0.00 | 0.00 | 0.00 | 0.00 | 0.00 | 0.00 | 0.00 | 0.00 | 0.00 | 0.00 | 0.00 | 0.00 | 0.00 | 0.00 |
| **C6** | 0.00 | 0.00 | 0.00 | 0.00 | 0.00 | 0.01 | 0.00 | 0.00 | 0.00 | 0.00 | 0.00 | 0.00 | 0.00 | 0.00 | 0.00 | 0.00 | 0.00 | 0.00 |
| **C7** | 0.00 | 0.00 | 0.00 | 0.00 | 0.00 | 0.00 | 0.09 | 0.00 | 0.00 | 0.00 | 0.00 | 0.00 | 0.00 | 0.00 | 0.00 | 0.00 | 0.00 | 0.00 |
| **C8** | 0.00 | 0.00 | 0.00 | 0.00 | 0.00 | 0.00 | 0.00 | 0.00 | 0.00 | 0.00 | 0.00 | 0.00 | 0.00 | 0.00 | 0.00 | 0.00 | 0.00 | 0.00 |
| **C9** | 0.00 | 0.00 | 0.00 | 0.00 | 0.00 | 0.00 | 0.00 | 0.00 | 0.00 | 0.00 | 0.00 | 0.00 | 0.00 | 0.00 | 0.00 | 0.00 | 0.00 | 0.00 |

Mutual affinity of the participants. Approximate dissociation constant K_d_ (25 °C) in M.

|  | **I** | **M1** | **M2** | **M3** | **M4** | **M5** | **M6** | **M7** | **S** | **C1** | **C2** | **C3** | **C4** | **C5** | **C6** | **C7** | **C8** | **C9** |
| --- | --- | --- | --- | --- | --- | --- | --- | --- | --- | --- | --- | --- | --- | --- | --- | --- | --- | --- |
| **I** | 1.0E+00 | 1.8E-04 | 7.9E-09 | 2.1E-09 | 4.4E-09 | 1.5E-04 | 6.0E-07 | 2.0E-04 | 3.1E-04 | 1.3E-08 | 1.0E+00 | 1.0E+00 | 1.0E+00 | 1.0E+00 | 1.0E+00 | 1.0E+00 | 1.0E+00 | 1.0E+00 |
| **M1** | 1.8E-04 | 1.0E+00 | 3.8E-10 | 9.1E-08 | 4.1E-09 | 3.0E-11 | 9.0E-07 | 1.8E-07 | 3.0E-06 | 1.0E+00 | 8.3E-10 | 1.0E+00 | 1.0E+00 | 1.0E+00 | 1.0E+00 | 1.0E+00 | 1.0E+00 | 1.0E+00 |
| **M2** | 7.9E-09 | 3.8E-10 | 1.0E+00 | 3.2E-09 | 6.8E-08 | 5.3E-09 | 3.6E-09 | 6.4E-09 | 7.5E-07 | 1.0E+00 | 1.0E+00 | 1.7E-05 | 1.0E+00 | 1.0E+00 | 1.0E+00 | 1.0E+00 | 1.0E+00 | 1.0E+00 |
| **M3** | 2.1E-09 | 9.1E-08 | 3.2E-09 | 1.0E+00 | 5.7E-07 | 6.2E-06 | 3.1E-04 | 2.5E-10 | 3.9E-07 | 1.0E+00 | 1.0E+00 | 1.0E+00 | 8.5E-10 | 1.0E+00 | 1.0E+00 | 1.0E+00 | 1.0E+00 | 1.0E+00 |
| **M4** | 4.4E-09 | 4.1E-09 | 6.8E-08 | 5.7E-07 | 1.0E+00 | 6.5E-07 | 6.3E-07 | 8.6E-07 | 1.4E-04 | 1.0E+00 | 1.0E+00 | 1.0E+00 | 1.0E+00 | 1.5E-05 | 1.0E+00 | 1.0E+00 | 1.0E+00 | 1.0E+00 |
| **M5** | 1.5E-04 | 3.0E-11 | 5.3E-09 | 6.2E-06 | 6.5E-07 | 1.0E+00 | 2.0E-08 | 6.9E-06 | 5.7E-06 | 1.0E+00 | 1.0E+00 | 1.0E+00 | 1.0E+00 | 1.0E+00 | 6.3E-09 | 1.0E+00 | 1.0E+00 | 1.0E+00 |
| **M6** | 6.0E-07 | 9.0E-07 | 3.6E-09 | 3.1E-04 | 6.3E-07 | 2.0E-08 | 1.0E+00 | 1.1E-09 | 2.4E-08 | 1.0E+00 | 1.0E+00 | 1.0E+00 | 1.0E+00 | 1.0E+00 | 1.0E+00 | 2.3E-10 | 1.0E+00 | 1.0E+00 |
| **M7** | 2.0E-04 | 1.8E-07 | 6.4E-09 | 2.5E-10 | 8.6E-07 | 6.9E-06 | 1.1E-09 | 1.0E+00 | 4.9E-05 | 1.0E+00 | 1.0E+00 | 1.0E+00 | 1.0E+00 | 1.0E+00 | 1.0E+00 | 1.0E+00 | 8.1E-06 | 1.0E+00 |
| **S** | 3.1E-04 | 3.0E-06 | 7.5E-07 | 3.9E-07 | 1.4E-04 | 5.7E-06 | 2.4E-08 | 4.9E-05 | 1.0E+00 | 1.0E+00 | 1.0E+00 | 1.0E+00 | 1.0E+00 | 1.0E+00 | 1.0E+00 | 1.0E+00 | 1.0E+00 | 2.2E-06 |
| **C1** | 1.3E-08 | 1.0E+00 | 1.0E+00 | 1.0E+00 | 1.0E+00 | 1.0E+00 | 1.0E+00 | 1.0E+00 | 1.0E+00 | 1.0E+00 | 1.0E+00 | 1.0E+00 | 1.0E+00 | 1.0E+00 | 1.0E+00 | 1.0E+00 | 1.0E+00 | 1.0E+00 |
| **C2** | 1.0E+00 | 8.3E-10 | 1.0E+00 | 1.0E+00 | 1.0E+00 | 1.0E+00 | 1.0E+00 | 1.0E+00 | 1.0E+00 | 1.0E+00 | 1.0E+00 | 1.0E+00 | 1.0E+00 | 1.0E+00 | 1.0E+00 | 1.0E+00 | 1.0E+00 | 1.0E+00 |
| **C3** | 1.0E+00 | 1.0E+00 | 1.7E-05 | 1.0E+00 | 1.0E+00 | 1.0E+00 | 1.0E+00 | 1.0E+00 | 1.0E+00 | 1.0E+00 | 1.0E+00 | 1.0E+00 | 1.0E+00 | 1.0E+00 | 1.0E+00 | 1.0E+00 | 1.0E+00 | 1.0E+00 |
| **C4** | 1.0E+00 | 1.0E+00 | 1.0E+00 | 8.5E-10 | 1.0E+00 | 1.0E+00 | 1.0E+00 | 1.0E+00 | 1.0E+00 | 1.0E+00 | 1.0E+00 | 1.0E+00 | 1.0E+00 | 1.0E+00 | 1.0E+00 | 1.0E+00 | 1.0E+00 | 1.0E+00 |
| **C5** | 1.0E+00 | 1.0E+00 | 1.0E+00 | 1.0E+00 | 1.5E-05 | 1.0E+00 | 1.0E+00 | 1.0E+00 | 1.0E+00 | 1.0E+00 | 1.0E+00 | 1.0E+00 | 1.0E+00 | 1.0E+00 | 1.0E+00 | 1.0E+00 | 1.0E+00 | 1.0E+00 |
| **C6** | 1.0E+00 | 1.0E+00 | 1.0E+00 | 1.0E+00 | 1.0E+00 | 6.3E-09 | 1.0E+00 | 1.0E+00 | 1.0E+00 | 1.0E+00 | 1.0E+00 | 1.0E+00 | 1.0E+00 | 1.0E+00 | 1.0E+00 | 1.0E+00 | 1.0E+00 | 1.0E+00 |
| **C7** | 1.0E+00 | 1.0E+00 | 1.0E+00 | 1.0E+00 | 1.0E+00 | 1.0E+00 | 2.3E-10 | 1.0E+00 | 1.0E+00 | 1.0E+00 | 1.0E+00 | 1.0E+00 | 1.0E+00 | 1.0E+00 | 1.0E+00 | 1.0E+00 | 1.0E+00 | 1.0E+00 |
| **C8** | 1.0E+00 | 1.0E+00 | 1.0E+00 | 1.0E+00 | 1.0E+00 | 1.0E+00 | 1.0E+00 | 8.1E-06 | 1.0E+00 | 1.0E+00 | 1.0E+00 | 1.0E+00 | 1.0E+00 | 1.0E+00 | 1.0E+00 | 1.0E+00 | 1.0E+00 | 1.0E+00 |
| **C9** | 1.0E+00 | 1.0E+00 | 1.0E+00 | 1.0E+00 | 1.0E+00 | 1.0E+00 | 1.0E+00 | 1.0E+00 | 2.2E-06 | 1.0E+00 | 1.0E+00 | 1.0E+00 | 1.0E+00 | 1.0E+00 | 1.0E+00 | 1.0E+00 | 1.0E+00 | 1.0E+00 |

Mutual affinity of the participants. Approximate dissociation constant K_d_ (50 °C) in M.

|  | **I** | **M1** | **M2** | **M3** | **M4** | **M5** | **M6** | **M7** | **S** | **C1** | **C2** | **C3** | **C4** | **C5** | **C6** | **C7** | **C8** | **C9** |
| --- | --- | --- | --- | --- | --- | --- | --- | --- | --- | --- | --- | --- | --- | --- | --- | --- | --- | --- |
| **I** | 1.0E+00 | 8.0E-04 | 1.6E-06 | 6.5E-07 | 1.1E-06 | 7.1E-04 | 2.4E-05 | 8.4E-04 | 1.1E-03 | 1.1E-04 | 1.0E+00 | 1.0E+00 | 1.0E+00 | 1.0E+00 | 1.0E+00 | 1.0E+00 | 1.0E+00 | 1.0E+00 |
| **M1** | 8.0E-04 | 1.0E+00 | 2.1E-07 | 7.5E-06 | 1.0E-06 | 3.9E-08 | 3.1E-05 | 1.1E-05 | 6.6E-05 | 1.0E+00 | 2.1E-05 | 1.0E+00 | 1.0E+00 | 1.0E+00 | 1.0E+00 | 1.0E+00 | 1.0E+00 | 1.0E+00 |
| **M2** | 1.6E-06 | 2.1E-07 | 1.0E+00 | 8.6E-07 | 6.2E-06 | 1.2E-06 | 9.3E-07 | 1.4E-06 | 2.8E-05 | 1.0E+00 | 1.0E+00 | 2.6E-03 | 1.0E+00 | 1.0E+00 | 1.0E+00 | 1.0E+00 | 1.0E+00 | 1.0E+00 |
| **M3** | 6.5E-07 | 7.5E-06 | 8.6E-07 | 1.0E+00 | 2.4E-05 | 1.0E-04 | 1.1E-03 | 1.6E-07 | 1.9E-05 | 1.0E+00 | 1.0E+00 | 1.0E+00 | 2.2E-05 | 1.0E+00 | 1.0E+00 | 1.0E+00 | 1.0E+00 | 1.0E+00 |
| **M4** | 1.1E-06 | 1.0E-06 | 6.2E-06 | 2.4E-05 | 1.0E+00 | 2.6E-05 | 2.5E-05 | 3.1E-05 | 7.0E-04 | 1.0E+00 | 1.0E+00 | 1.0E+00 | 1.0E+00 | 2.5E-03 | 1.0E+00 | 1.0E+00 | 1.0E+00 | 1.0E+00 |
| **M5** | 7.1E-04 | 3.9E-08 | 1.2E-06 | 1.0E-04 | 2.6E-05 | 1.0E+00 | 2.8E-06 | 1.1E-04 | 9.9E-05 | 1.0E+00 | 1.0E+00 | 1.0E+00 | 1.0E+00 | 1.0E+00 | 7.3E-05 | 1.0E+00 | 1.0E+00 | 1.0E+00 |
| **M6** | 2.4E-05 | 3.1E-05 | 9.3E-07 | 1.1E-03 | 2.5E-05 | 2.8E-06 | 1.0E+00 | 4.2E-07 | 3.2E-06 | 1.0E+00 | 1.0E+00 | 1.0E+00 | 1.0E+00 | 1.0E+00 | 1.0E+00 | 9.3E-06 | 1.0E+00 | 1.0E+00 |
| **M7** | 8.4E-04 | 1.1E-05 | 1.4E-06 | 1.6E-07 | 3.1E-05 | 1.1E-04 | 4.2E-07 | 1.0E+00 | 3.7E-04 | 1.0E+00 | 1.0E+00 | 1.0E+00 | 1.0E+00 | 1.0E+00 | 1.0E+00 | 1.0E+00 | 2.0E-03 | 1.0E+00 |
| **S** | 1.1E-03 | 6.6E-05 | 2.8E-05 | 1.9E-05 | 7.0E-04 | 9.9E-05 | 3.2E-06 | 3.7E-04 | 1.0E+00 | 1.0E+00 | 1.0E+00 | 1.0E+00 | 1.0E+00 | 1.0E+00 | 1.0E+00 | 1.0E+00 | 1.0E+00 | 1.3E-03 |
| **C1** | 1.1E-04 | 1.0E+00 | 1.0E+00 | 1.0E+00 | 1.0E+00 | 1.0E+00 | 1.0E+00 | 1.0E+00 | 1.0E+00 | 1.0E+00 | 1.0E+00 | 1.0E+00 | 1.0E+00 | 1.0E+00 | 1.0E+00 | 1.0E+00 | 1.0E+00 | 1.0E+00 |
| **C2** | 1.0E+00 | 2.1E-05 | 1.0E+00 | 1.0E+00 | 1.0E+00 | 1.0E+00 | 1.0E+00 | 1.0E+00 | 1.0E+00 | 1.0E+00 | 1.0E+00 | 1.0E+00 | 1.0E+00 | 1.0E+00 | 1.0E+00 | 1.0E+00 | 1.0E+00 | 1.0E+00 |
| **C3** | 1.0E+00 | 1.0E+00 | 2.6E-03 | 1.0E+00 | 1.0E+00 | 1.0E+00 | 1.0E+00 | 1.0E+00 | 1.0E+00 | 1.0E+00 | 1.0E+00 | 1.0E+00 | 1.0E+00 | 1.0E+00 | 1.0E+00 | 1.0E+00 | 1.0E+00 | 1.0E+00 |
| **C4** | 1.0E+00 | 1.0E+00 | 1.0E+00 | 2.2E-05 | 1.0E+00 | 1.0E+00 | 1.0E+00 | 1.0E+00 | 1.0E+00 | 1.0E+00 | 1.0E+00 | 1.0E+00 | 1.0E+00 | 1.0E+00 | 1.0E+00 | 1.0E+00 | 1.0E+00 | 1.0E+00 |
| **C5** | 1.0E+00 | 1.0E+00 | 1.0E+00 | 1.0E+00 | 2.5E-03 | 1.0E+00 | 1.0E+00 | 1.0E+00 | 1.0E+00 | 1.0E+00 | 1.0E+00 | 1.0E+00 | 1.0E+00 | 1.0E+00 | 1.0E+00 | 1.0E+00 | 1.0E+00 | 1.0E+00 |
| **C6** | 1.0E+00 | 1.0E+00 | 1.0E+00 | 1.0E+00 | 1.0E+00 | 7.3E-05 | 1.0E+00 | 1.0E+00 | 1.0E+00 | 1.0E+00 | 1.0E+00 | 1.0E+00 | 1.0E+00 | 1.0E+00 | 1.0E+00 | 1.0E+00 | 1.0E+00 | 1.0E+00 |
| **C7** | 1.0E+00 | 1.0E+00 | 1.0E+00 | 1.0E+00 | 1.0E+00 | 1.0E+00 | 9.3E-06 | 1.0E+00 | 1.0E+00 | 1.0E+00 | 1.0E+00 | 1.0E+00 | 1.0E+00 | 1.0E+00 | 1.0E+00 | 1.0E+00 | 1.0E+00 | 1.0E+00 |
| **C8** | 1.0E+00 | 1.0E+00 | 1.0E+00 | 1.0E+00 | 1.0E+00 | 1.0E+00 | 1.0E+00 | 2.0E-03 | 1.0E+00 | 1.0E+00 | 1.0E+00 | 1.0E+00 | 1.0E+00 | 1.0E+00 | 1.0E+00 | 1.0E+00 | 1.0E+00 | 1.0E+00 |
| **C9** | 1.0E+00 | 1.0E+00 | 1.0E+00 | 1.0E+00 | 1.0E+00 | 1.0E+00 | 1.0E+00 | 1.0E+00 | 1.3E-03 | 1.0E+00 | 1.0E+00 | 1.0E+00 | 1.0E+00 | 1.0E+00 | 1.0E+00 | 1.0E+00 | 1.0E+00 | 1.0E+00 |

#### Line function – control system without compensatory participants.

Concentrations of the participants.

| **ID** | **Concentration, μM** |
| --- | --- |
| I | 0.54 |
| M1 | 5.78 |
| M2 | 1.91 |
| M3 | 7.58 |
| M4 | 7.01 |
| M5 | 3.36 |
| M6 | 9.23 |
| M7 | 8.59 |
| S | 0.03 |

Mutual affinity of the participants. Concentration of the duplex C_AB_ (μM) when oligos are mixed at 1 μM each (25 °C).

|  | **I** | **M1** | **M2** | **M3** | **M4** | **M5** | **M6** | **M7** | **S** |
| --- | --- | --- | --- | --- | --- | --- | --- | --- | --- |
| **I** | 0.00 | 0.75 | 0.80 | 0.90 | 0.95 | 0.02 | 0.77 | 0.65 | 0.21 |
| **M1** | 0.75 | 0.00 | 0.98 | 0.63 | 0.97 | 0.99 | 0.97 | 0.55 | 0.07 |
| **M2** | 0.80 | 0.98 | 0.00 | 0.99 | 0.93 | 0.98 | 0.97 | 0.97 | 0.66 |
| **M3** | 0.90 | 0.63 | 0.99 | 0.00 | 0.70 | 0.20 | 0.01 | 0.99 | 0.55 |
| **M4** | 0.95 | 0.97 | 0.93 | 0.70 | 0.00 | 0.43 | 0.49 | 0.10 | 0.01 |
| **M5** | 0.02 | 0.99 | 0.98 | 0.20 | 0.43 | 0.00 | 0.72 | 0.06 | 0.30 |
| **M6** | 0.77 | 0.97 | 0.97 | 0.01 | 0.49 | 0.72 | 0.00 | 0.99 | 0.88 |
| **M7** | 0.65 | 0.55 | 0.97 | 0.99 | 0.10 | 0.06 | 0.99 | 0.00 | 0.05 |
| **S** | 0.21 | 0.07 | 0.66 | 0.55 | 0.01 | 0.30 | 0.88 | 0.05 | 0.00 |

Mutual affinity of the participants. Concentration of the duplex C_AB_ (μM) when oligos are mixed at 1 μM each (50 °C).

|  | **I** | **M1** | **M2** | **M3** | **M4** | **M5** | **M6** | **M7** | **S** |
| --- | --- | --- | --- | --- | --- | --- | --- | --- | --- |
| **I** | 0.00 | 0.11 | 0.14 | 0.27 | 0.44 | 0.00 | 0.13 | 0.07 | 0.01 |
| **M1** | 0.11 | 0.00 | 0.61 | 0.07 | 0.53 | 0.69 | 0.52 | 0.05 | 0.01 |
| **M2** | 0.14 | 0.61 | 0.00 | 0.74 | 0.35 | 0.60 | 0.57 | 0.53 | 0.08 |
| **M3** | 0.27 | 0.07 | 0.74 | 0.00 | 0.09 | 0.01 | 0.00 | 0.77 | 0.05 |
| **M4** | 0.44 | 0.53 | 0.35 | 0.09 | 0.00 | 0.03 | 0.04 | 0.01 | 0.00 |
| **M5** | 0.00 | 0.69 | 0.60 | 0.01 | 0.03 | 0.00 | 0.10 | 0.01 | 0.02 |
| **M6** | 0.13 | 0.52 | 0.57 | 0.00 | 0.04 | 0.10 | 0.00 | 0.71 | 0.24 |
| **M7** | 0.07 | 0.05 | 0.53 | 0.77 | 0.01 | 0.01 | 0.71 | 0.00 | 0.01 |
| **S** | 0.01 | 0.01 | 0.08 | 0.05 | 0.00 | 0.02 | 0.24 | 0.01 | 0.00 |

Mutual affinity of the participants. Approximate dissociation constant K_d_ (25 °C) in M.

|  | **I** | **M1** | **M2** | **M3** | **M4** | **M5** | **M6** | **M7** | **S** |
| --- | --- | --- | --- | --- | --- | --- | --- | --- | --- |
| **I** | 1.0E+00 | 8.0E-08 | 5.2E-08 | 1.2E-08 | 2.5E-09 | 5.1E-05 | 6.7E-08 | 1.9E-07 | 3.1E-06 |
| **M1** | 8.0E-08 | 1.0E+00 | 4.8E-10 | 2.2E-07 | 1.1E-09 | 1.9E-10 | 1.2E-09 | 3.6E-07 | 1.3E-05 |
| **M2** | 5.2E-08 | 4.8E-10 | 1.0E+00 | 1.0E-10 | 5.5E-09 | 5.4E-10 | 7.4E-10 | 1.1E-09 | 1.7E-07 |
| **M3** | 1.2E-08 | 2.2E-07 | 1.0E-10 | 1.0E+00 | 1.3E-07 | 3.2E-06 | 6.8E-05 | 7.3E-11 | 3.8E-07 |
| **M4** | 2.5E-09 | 1.1E-09 | 5.5E-09 | 1.3E-07 | 1.0E+00 | 7.6E-07 | 5.2E-07 | 8.3E-06 | 1.1E-04 |
| **M5** | 5.1E-05 | 1.9E-10 | 5.4E-10 | 3.2E-06 | 7.6E-07 | 1.0E+00 | 1.1E-07 | 1.5E-05 | 1.6E-06 |
| **M6** | 6.7E-08 | 1.2E-09 | 7.4E-10 | 6.8E-05 | 5.2E-07 | 1.1E-07 | 1.0E+00 | 1.5E-10 | 1.6E-08 |
| **M7** | 1.9E-07 | 3.6E-07 | 1.1E-09 | 7.3E-11 | 8.3E-06 | 1.5E-05 | 1.5E-10 | 1.0E+00 | 1.7E-05 |
| **S** | 3.1E-06 | 1.3E-05 | 1.7E-07 | 3.8E-07 | 1.1E-04 | 1.6E-06 | 1.6E-08 | 1.7E-05 | 1.0E+00 |

Mutual affinity of the participants. Approximate dissociation constant K_d_ (50 °C) in M.

|  | **I** | **M1** | **M2** | **M3** | **M4** | **M5** | **M6** | **M7** | **S** |
| --- | --- | --- | --- | --- | --- | --- | --- | --- | --- |
| **I** | 1.0E+00 | 6.9E-06 | 5.2E-06 | 2.0E-06 | 7.3E-07 | 3.7E-04 | 6.1E-06 | 1.2E-05 | 6.7E-05 |
| **M1** | 6.9E-06 | 1.0E+00 | 2.5E-07 | 1.3E-05 | 4.2E-07 | 1.4E-07 | 4.5E-07 | 1.8E-05 | 1.6E-04 |
| **M2** | 5.2E-06 | 2.5E-07 | 1.0E+00 | 8.9E-08 | 1.2E-06 | 2.7E-07 | 3.3E-07 | 4.3E-07 | 1.1E-05 |
| **M3** | 2.0E-06 | 1.3E-05 | 8.9E-08 | 1.0E+00 | 9.2E-06 | 6.9E-05 | 4.4E-04 | 7.1E-08 | 1.8E-05 |
| **M4** | 7.3E-07 | 4.2E-07 | 1.2E-06 | 9.2E-06 | 1.0E+00 | 2.8E-05 | 2.2E-05 | 1.2E-04 | 5.9E-04 |
| **M5** | 3.7E-04 | 1.4E-07 | 2.7E-07 | 6.9E-05 | 2.8E-05 | 1.0E+00 | 8.5E-06 | 1.8E-04 | 4.5E-05 |
| **M6** | 6.1E-06 | 4.5E-07 | 3.3E-07 | 4.4E-04 | 2.2E-05 | 8.5E-06 | 1.0E+00 | 1.1E-07 | 2.5E-06 |
| **M7** | 1.2E-05 | 1.8E-05 | 4.3E-07 | 7.1E-08 | 1.2E-04 | 1.8E-04 | 1.1E-07 | 1.0E+00 | 1.9E-04 |
| **S** | 6.7E-05 | 1.6E-04 | 1.1E-05 | 1.8E-05 | 5.9E-04 | 4.5E-05 | 2.5E-06 | 1.9E-04 | 1.0E+00 |

#### Quadratic function – temperature-independent system

Concentrations of the participants.

| **ID** | **Concentration, μM** |
| --- | --- |
| I | 4.66 |
| M1 | 2.52 |
| M2 | 2.75 |
| M3 | 1.70 |
| M4 | 6.92 |
| M5 | 5.33 |
| M6 | 7.11 |
| S | 0.03 |
| C1 | 2.77 |
| C2 | 0.14 |
| C3 | 1.73 |
| C4 | 1.75 |
| C5 | 4.86 |
| C6 | 0.00 |
| C7 | 4.25 |
| C8 | 6.80 |

Mutual affinity of the participants. Concentration of the duplex C_AB_ (μM) when oligos are mixed at 1 μM each (25 °C).

|  | **I** | **M1** | **M2** | **M3** | **M4** | **M5** | **M6** | **S** | **C1** | **C2** | **C3** | **C4** | **C5** | **C6** | **C7** | **C8** |
| --- | --- | --- | --- | --- | --- | --- | --- | --- | --- | --- | --- | --- | --- | --- | --- | --- |
| **I** | 0.00 | 0.97 | 0.45 | 0.01 | 0.99 | 0.00 | 0.84 | 0.00 | 0.91 | 0.00 | 0.00 | 0.00 | 0.00 | 0.00 | 0.00 | 0.00 |
| **M1** | 0.97 | 0.00 | 0.03 | 0.99 | 0.99 | 0.84 | 0.93 | 0.34 | 0.00 | 0.69 | 0.00 | 0.00 | 0.00 | 0.00 | 0.00 | 0.00 |
| **M2** | 0.45 | 0.03 | 0.00 | 0.82 | 0.07 | 0.82 | 0.72 | 0.00 | 0.00 | 0.00 | 0.94 | 0.00 | 0.00 | 0.00 | 0.00 | 0.00 |
| **M3** | 0.01 | 0.99 | 0.82 | 0.00 | 0.01 | 0.12 | 0.00 | 0.25 | 0.00 | 0.00 | 0.00 | 0.99 | 0.00 | 0.00 | 0.00 | 0.00 |
| **M4** | 0.99 | 0.99 | 0.07 | 0.01 | 0.00 | 0.51 | 0.85 | 0.01 | 0.00 | 0.00 | 0.00 | 0.00 | 0.97 | 0.00 | 0.00 | 0.00 |
| **M5** | 0.00 | 0.84 | 0.82 | 0.12 | 0.51 | 0.00 | 0.98 | 0.10 | 0.00 | 0.00 | 0.00 | 0.00 | 0.00 | 0.04 | 0.00 | 0.00 |
| **M6** | 0.84 | 0.93 | 0.72 | 0.00 | 0.85 | 0.98 | 0.00 | 0.91 | 0.00 | 0.00 | 0.00 | 0.00 | 0.00 | 0.00 | 0.99 | 0.00 |
| **S** | 0.00 | 0.34 | 0.00 | 0.25 | 0.01 | 0.10 | 0.91 | 0.00 | 0.00 | 0.00 | 0.00 | 0.00 | 0.00 | 0.00 | 0.00 | 0.02 |
| **C1** | 0.91 | 0.00 | 0.00 | 0.00 | 0.00 | 0.00 | 0.00 | 0.00 | 0.00 | 0.00 | 0.00 | 0.00 | 0.00 | 0.00 | 0.00 | 0.00 |
| **C2** | 0.00 | 0.69 | 0.00 | 0.00 | 0.00 | 0.00 | 0.00 | 0.00 | 0.00 | 0.00 | 0.00 | 0.00 | 0.00 | 0.00 | 0.00 | 0.00 |
| **C3** | 0.00 | 0.00 | 0.94 | 0.00 | 0.00 | 0.00 | 0.00 | 0.00 | 0.00 | 0.00 | 0.00 | 0.00 | 0.00 | 0.00 | 0.00 | 0.00 |
| **C4** | 0.00 | 0.00 | 0.00 | 0.99 | 0.00 | 0.00 | 0.00 | 0.00 | 0.00 | 0.00 | 0.00 | 0.00 | 0.00 | 0.00 | 0.00 | 0.00 |
| **C5** | 0.00 | 0.00 | 0.00 | 0.00 | 0.97 | 0.00 | 0.00 | 0.00 | 0.00 | 0.00 | 0.00 | 0.00 | 0.00 | 0.00 | 0.00 | 0.00 |
| **C6** | 0.00 | 0.00 | 0.00 | 0.00 | 0.00 | 0.04 | 0.00 | 0.00 | 0.00 | 0.00 | 0.00 | 0.00 | 0.00 | 0.00 | 0.00 | 0.00 |
| **C7** | 0.00 | 0.00 | 0.00 | 0.00 | 0.00 | 0.00 | 0.99 | 0.00 | 0.00 | 0.00 | 0.00 | 0.00 | 0.00 | 0.00 | 0.00 | 0.00 |
| **C8** | 0.00 | 0.00 | 0.00 | 0.00 | 0.00 | 0.00 | 0.00 | 0.02 | 0.00 | 0.00 | 0.00 | 0.00 | 0.00 | 0.00 | 0.00 | 0.00 |

Mutual affinity of the participants. Concentration of the duplex C_AB_ (μM) when oligos are mixed at 1 μM each (50 °C).

|  | **I** | **M1** | **M2** | **M3** | **M4** | **M5** | **M6** | **S** | **C1** | **C2** | **C3** | **C4** | **C5** | **C6** | **C7** | **C8** |
| --- | --- | --- | --- | --- | --- | --- | --- | --- | --- | --- | --- | --- | --- | --- | --- | --- |
| **I** | 0.00 | 0.57 | 0.04 | 0.00 | 0.82 | 0.00 | 0.18 | 0.00 | 0.01 | 0.00 | 0.00 | 0.00 | 0.00 | 0.00 | 0.00 | 0.00 |
| **M1** | 0.57 | 0.00 | 0.00 | 0.82 | 0.81 | 0.18 | 0.35 | 0.02 | 0.00 | 0.00 | 0.00 | 0.00 | 0.00 | 0.00 | 0.00 | 0.00 |
| **M2** | 0.04 | 0.00 | 0.00 | 0.17 | 0.01 | 0.16 | 0.10 | 0.00 | 0.00 | 0.00 | 0.02 | 0.00 | 0.00 | 0.00 | 0.00 | 0.00 |
| **M3** | 0.00 | 0.82 | 0.17 | 0.00 | 0.00 | 0.01 | 0.00 | 0.02 | 0.00 | 0.00 | 0.00 | 0.22 | 0.00 | 0.00 | 0.00 | 0.00 |
| **M4** | 0.82 | 0.81 | 0.01 | 0.00 | 0.00 | 0.04 | 0.19 | 0.00 | 0.00 | 0.00 | 0.00 | 0.00 | 0.04 | 0.00 | 0.00 | 0.00 |
| **M5** | 0.00 | 0.18 | 0.16 | 0.01 | 0.04 | 0.00 | 0.60 | 0.01 | 0.00 | 0.00 | 0.00 | 0.00 | 0.00 | 0.00 | 0.00 | 0.00 |
| **M6** | 0.18 | 0.35 | 0.10 | 0.00 | 0.19 | 0.60 | 0.00 | 0.29 | 0.00 | 0.00 | 0.00 | 0.00 | 0.00 | 0.00 | 0.09 | 0.00 |
| **S** | 0.00 | 0.02 | 0.00 | 0.02 | 0.00 | 0.01 | 0.29 | 0.00 | 0.00 | 0.00 | 0.00 | 0.00 | 0.00 | 0.00 | 0.00 | 0.00 |
| **C1** | 0.01 | 0.00 | 0.00 | 0.00 | 0.00 | 0.00 | 0.00 | 0.00 | 0.00 | 0.00 | 0.00 | 0.00 | 0.00 | 0.00 | 0.00 | 0.00 |
| **C2** | 0.00 | 0.00 | 0.00 | 0.00 | 0.00 | 0.00 | 0.00 | 0.00 | 0.00 | 0.00 | 0.00 | 0.00 | 0.00 | 0.00 | 0.00 | 0.00 |
| **C3** | 0.00 | 0.00 | 0.02 | 0.00 | 0.00 | 0.00 | 0.00 | 0.00 | 0.00 | 0.00 | 0.00 | 0.00 | 0.00 | 0.00 | 0.00 | 0.00 |
| **C4** | 0.00 | 0.00 | 0.00 | 0.22 | 0.00 | 0.00 | 0.00 | 0.00 | 0.00 | 0.00 | 0.00 | 0.00 | 0.00 | 0.00 | 0.00 | 0.00 |
| **C5** | 0.00 | 0.00 | 0.00 | 0.00 | 0.04 | 0.00 | 0.00 | 0.00 | 0.00 | 0.00 | 0.00 | 0.00 | 0.00 | 0.00 | 0.00 | 0.00 |
| **C6** | 0.00 | 0.00 | 0.00 | 0.00 | 0.00 | 0.00 | 0.00 | 0.00 | 0.00 | 0.00 | 0.00 | 0.00 | 0.00 | 0.00 | 0.00 | 0.00 |
| **C7** | 0.00 | 0.00 | 0.00 | 0.00 | 0.00 | 0.00 | 0.09 | 0.00 | 0.00 | 0.00 | 0.00 | 0.00 | 0.00 | 0.00 | 0.00 | 0.00 |
| **C8** | 0.00 | 0.00 | 0.00 | 0.00 | 0.00 | 0.00 | 0.00 | 0.00 | 0.00 | 0.00 | 0.00 | 0.00 | 0.00 | 0.00 | 0.00 | 0.00 |

Mutual affinity of the participants. Approximate dissociation constant K_d_ (25 °C) in M.

|  | **I** | **M1** | **M2** | **M3** | **M4** | **M5** | **M6** | **S** | **C1** | **C2** | **C3** | **C4** | **C5** | **C6** | **C7** | **C8** |
| --- | --- | --- | --- | --- | --- | --- | --- | --- | --- | --- | --- | --- | --- | --- | --- | --- |
| **I** | 1.0E+00 | 7.6E-10 | 6.6E-07 | 1.7E-04 | 3.0E-11 | 2.4E-04 | 3.0E-08 | 3.1E-04 | 9.6E-09 | 1.0E+00 | 1.0E+00 | 1.0E+00 | 1.0E+00 | 1.0E+00 | 1.0E+00 | 1.0E+00 |
| **M1** | 7.6E-10 | 1.0E+00 | 3.0E-05 | 3.0E-11 | 3.4E-11 | 3.1E-08 | 5.4E-09 | 1.3E-06 | 1.0E+00 | 1.4E-07 | 1.0E+00 | 1.0E+00 | 1.0E+00 | 1.0E+00 | 1.0E+00 | 1.0E+00 |
| **M2** | 6.6E-07 | 3.0E-05 | 1.0E+00 | 3.7E-08 | 1.3E-05 | 3.8E-08 | 1.1E-07 | 2.4E-04 | 1.0E+00 | 1.0E+00 | 4.2E-09 | 1.0E+00 | 1.0E+00 | 1.0E+00 | 1.0E+00 | 1.0E+00 |
| **M3** | 1.7E-04 | 3.0E-11 | 3.7E-08 | 1.0E+00 | 6.9E-05 | 6.2E-06 | 3.0E-04 | 2.2E-06 | 1.0E+00 | 1.0E+00 | 1.0E+00 | 3.6E-11 | 1.0E+00 | 1.0E+00 | 1.0E+00 | 1.0E+00 |
| **M4** | 3.0E-11 | 3.4E-11 | 1.3E-05 | 6.9E-05 | 1.0E+00 | 4.6E-07 | 2.8E-08 | 9.5E-05 | 1.0E+00 | 1.0E+00 | 1.0E+00 | 1.0E+00 | 1.1E-09 | 1.0E+00 | 1.0E+00 | 1.0E+00 |
| **M5** | 2.4E-04 | 3.1E-08 | 3.8E-08 | 6.2E-06 | 4.6E-07 | 1.0E+00 | 5.5E-10 | 8.5E-06 | 1.0E+00 | 1.0E+00 | 1.0E+00 | 1.0E+00 | 1.0E+00 | 2.4E-05 | 1.0E+00 | 1.0E+00 |
| **M6** | 3.0E-08 | 5.4E-09 | 1.1E-07 | 3.0E-04 | 2.8E-08 | 5.5E-10 | 1.0E+00 | 9.2E-09 | 1.0E+00 | 1.0E+00 | 1.0E+00 | 1.0E+00 | 1.0E+00 | 1.0E+00 | 2.3E-10 | 1.0E+00 |
| **S** | 3.1E-04 | 1.3E-06 | 2.4E-04 | 2.2E-06 | 9.5E-05 | 8.5E-06 | 9.2E-09 | 1.0E+00 | 1.0E+00 | 1.0E+00 | 1.0E+00 | 1.0E+00 | 1.0E+00 | 1.0E+00 | 1.0E+00 | 5.5E-05 |
| **C1** | 9.6E-09 | 1.0E+00 | 1.0E+00 | 1.0E+00 | 1.0E+00 | 1.0E+00 | 1.0E+00 | 1.0E+00 | 1.0E+00 | 1.0E+00 | 1.0E+00 | 1.0E+00 | 1.0E+00 | 1.0E+00 | 1.0E+00 | 1.0E+00 |
| **C2** | 1.0E+00 | 1.4E-07 | 1.0E+00 | 1.0E+00 | 1.0E+00 | 1.0E+00 | 1.0E+00 | 1.0E+00 | 1.0E+00 | 1.0E+00 | 1.0E+00 | 1.0E+00 | 1.0E+00 | 1.0E+00 | 1.0E+00 | 1.0E+00 |
| **C3** | 1.0E+00 | 1.0E+00 | 4.2E-09 | 1.0E+00 | 1.0E+00 | 1.0E+00 | 1.0E+00 | 1.0E+00 | 1.0E+00 | 1.0E+00 | 1.0E+00 | 1.0E+00 | 1.0E+00 | 1.0E+00 | 1.0E+00 | 1.0E+00 |
| **C4** | 1.0E+00 | 1.0E+00 | 1.0E+00 | 3.6E-11 | 1.0E+00 | 1.0E+00 | 1.0E+00 | 1.0E+00 | 1.0E+00 | 1.0E+00 | 1.0E+00 | 1.0E+00 | 1.0E+00 | 1.0E+00 | 1.0E+00 | 1.0E+00 |
| **C5** | 1.0E+00 | 1.0E+00 | 1.0E+00 | 1.0E+00 | 1.1E-09 | 1.0E+00 | 1.0E+00 | 1.0E+00 | 1.0E+00 | 1.0E+00 | 1.0E+00 | 1.0E+00 | 1.0E+00 | 1.0E+00 | 1.0E+00 | 1.0E+00 |
| **C6** | 1.0E+00 | 1.0E+00 | 1.0E+00 | 1.0E+00 | 1.0E+00 | 2.4E-05 | 1.0E+00 | 1.0E+00 | 1.0E+00 | 1.0E+00 | 1.0E+00 | 1.0E+00 | 1.0E+00 | 1.0E+00 | 1.0E+00 | 1.0E+00 |
| **C7** | 1.0E+00 | 1.0E+00 | 1.0E+00 | 1.0E+00 | 1.0E+00 | 1.0E+00 | 2.3E-10 | 1.0E+00 | 1.0E+00 | 1.0E+00 | 1.0E+00 | 1.0E+00 | 1.0E+00 | 1.0E+00 | 1.0E+00 | 1.0E+00 |
| **C8** | 1.0E+00 | 1.0E+00 | 1.0E+00 | 1.0E+00 | 1.0E+00 | 1.0E+00 | 1.0E+00 | 5.5E-05 | 1.0E+00 | 1.0E+00 | 1.0E+00 | 1.0E+00 | 1.0E+00 | 1.0E+00 | 1.0E+00 | 1.0E+00 |

Mutual affinity of the participants. Approximate dissociation constant K_d_ (50 °C) in M.

|  | **I** | **M1** | **M2** | **M3** | **M4** | **M5** | **M6** | **S** | **C1** | **C2** | **C3** | **C4** | **C5** | **C6** | **C7** | **C8** |
| --- | --- | --- | --- | --- | --- | --- | --- | --- | --- | --- | --- | --- | --- | --- | --- | --- |
| **I** | 1.0E+00 | 3.3E-07 | 2.6E-05 | 7.6E-04 | 3.9E-08 | 9.5E-04 | 3.7E-06 | 1.1E-03 | 9.3E-05 | 1.0E+00 | 1.0E+00 | 1.0E+00 | 1.0E+00 | 1.0E+00 | 1.0E+00 | 1.0E+00 |
| **M1** | 3.3E-07 | 1.0E+00 | 2.7E-04 | 3.9E-08 | 4.2E-08 | 3.7E-06 | 1.2E-06 | 3.9E-05 | 1.0E+00 | 3.8E-04 | 1.0E+00 | 1.0E+00 | 1.0E+00 | 1.0E+00 | 1.0E+00 | 1.0E+00 |
| **M2** | 2.6E-05 | 2.7E-04 | 1.0E+00 | 4.2E-06 | 1.6E-04 | 4.2E-06 | 8.4E-06 | 9.5E-04 | 1.0E+00 | 1.0E+00 | 5.7E-05 | 1.0E+00 | 1.0E+00 | 1.0E+00 | 1.0E+00 | 1.0E+00 |
| **M3** | 7.6E-04 | 3.9E-08 | 4.2E-06 | 1.0E+00 | 4.5E-04 | 1.0E-04 | 1.1E-03 | 5.5E-05 | 1.0E+00 | 1.0E+00 | 1.0E+00 | 2.7E-06 | 1.0E+00 | 1.0E+00 | 1.0E+00 | 1.0E+00 |
| **M4** | 3.9E-08 | 4.2E-08 | 1.6E-04 | 4.5E-04 | 1.0E+00 | 2.1E-05 | 3.5E-06 | 5.4E-04 | 1.0E+00 | 1.0E+00 | 1.0E+00 | 1.0E+00 | 2.6E-05 | 1.0E+00 | 1.0E+00 | 1.0E+00 |
| **M5** | 9.5E-04 | 3.7E-06 | 4.2E-06 | 1.0E-04 | 2.1E-05 | 1.0E+00 | 2.7E-07 | 1.3E-04 | 1.0E+00 | 1.0E+00 | 1.0E+00 | 1.0E+00 | 1.0E+00 | 2.9E-03 | 1.0E+00 | 1.0E+00 |
| **M6** | 3.7E-06 | 1.2E-06 | 8.4E-06 | 1.1E-03 | 3.5E-06 | 2.7E-07 | 1.0E+00 | 1.7E-06 | 1.0E+00 | 1.0E+00 | 1.0E+00 | 1.0E+00 | 1.0E+00 | 1.0E+00 | 9.3E-06 | 1.0E+00 |
| **S** | 1.1E-03 | 3.9E-05 | 9.5E-04 | 5.5E-05 | 5.4E-04 | 1.3E-04 | 1.7E-06 | 1.0E+00 | 1.0E+00 | 1.0E+00 | 1.0E+00 | 1.0E+00 | 1.0E+00 | 1.0E+00 | 1.0E+00 | 3.6E-03 |
| **C1** | 9.3E-05 | 1.0E+00 | 1.0E+00 | 1.0E+00 | 1.0E+00 | 1.0E+00 | 1.0E+00 | 1.0E+00 | 1.0E+00 | 1.0E+00 | 1.0E+00 | 1.0E+00 | 1.0E+00 | 1.0E+00 | 1.0E+00 | 1.0E+00 |
| **C2** | 1.0E+00 | 3.8E-04 | 1.0E+00 | 1.0E+00 | 1.0E+00 | 1.0E+00 | 1.0E+00 | 1.0E+00 | 1.0E+00 | 1.0E+00 | 1.0E+00 | 1.0E+00 | 1.0E+00 | 1.0E+00 | 1.0E+00 | 1.0E+00 |
| **C3** | 1.0E+00 | 1.0E+00 | 5.7E-05 | 1.0E+00 | 1.0E+00 | 1.0E+00 | 1.0E+00 | 1.0E+00 | 1.0E+00 | 1.0E+00 | 1.0E+00 | 1.0E+00 | 1.0E+00 | 1.0E+00 | 1.0E+00 | 1.0E+00 |
| **C4** | 1.0E+00 | 1.0E+00 | 1.0E+00 | 2.7E-06 | 1.0E+00 | 1.0E+00 | 1.0E+00 | 1.0E+00 | 1.0E+00 | 1.0E+00 | 1.0E+00 | 1.0E+00 | 1.0E+00 | 1.0E+00 | 1.0E+00 | 1.0E+00 |
| **C5** | 1.0E+00 | 1.0E+00 | 1.0E+00 | 1.0E+00 | 2.6E-05 | 1.0E+00 | 1.0E+00 | 1.0E+00 | 1.0E+00 | 1.0E+00 | 1.0E+00 | 1.0E+00 | 1.0E+00 | 1.0E+00 | 1.0E+00 | 1.0E+00 |
| **C6** | 1.0E+00 | 1.0E+00 | 1.0E+00 | 1.0E+00 | 1.0E+00 | 2.9E-03 | 1.0E+00 | 1.0E+00 | 1.0E+00 | 1.0E+00 | 1.0E+00 | 1.0E+00 | 1.0E+00 | 1.0E+00 | 1.0E+00 | 1.0E+00 |
| **C7** | 1.0E+00 | 1.0E+00 | 1.0E+00 | 1.0E+00 | 1.0E+00 | 1.0E+00 | 9.3E-06 | 1.0E+00 | 1.0E+00 | 1.0E+00 | 1.0E+00 | 1.0E+00 | 1.0E+00 | 1.0E+00 | 1.0E+00 | 1.0E+00 |
| **C8** | 1.0E+00 | 1.0E+00 | 1.0E+00 | 1.0E+00 | 1.0E+00 | 1.0E+00 | 1.0E+00 | 3.6E-03 | 1.0E+00 | 1.0E+00 | 1.0E+00 | 1.0E+00 | 1.0E+00 | 1.0E+00 | 1.0E+00 | 1.0E+00 |

#### Quadratic function – control system without compensatory participants

Concentrations of the participants.

| **ID** | **Concentration, μM** |
| --- | --- |
| I | 4.66 |
| M1 | 5.44 |
| M2 | 2.29 |
| M3 | 1.50 |
| M4 | 7.66 |
| M5 | 5.69 |
| M6 | 7.45 |
| M7 | 0.75 |
| S | 0.03 |

Mutual affinity of the participants. Concentration of the duplex C_AB_ (μM) when oligos are mixed at 1 μM each (25 °C).

|  | **I** | **M1** | **M2** | **M3** | **M4** | **M5** | **M6** | **M7** | **S** |
| --- | --- | --- | --- | --- | --- | --- | --- | --- | --- |
| **I** | 0.00 | 0.99 | 0.07 | 0.22 | 0.99 | 0.13 | 0.05 | 0.73 | 0.18 |
| **M1** | 0.99 | 0.00 | 0.77 | 0.37 | 0.06 | 0.89 | 0.99 | 0.01 | 0.62 |
| **M2** | 0.07 | 0.77 | 0.00 | 0.75 | 0.16 | 0.33 | 0.01 | 0.99 | 0.00 |
| **M3** | 0.22 | 0.37 | 0.75 | 0.00 | 0.33 | 0.38 | 0.03 | 0.01 | 0.01 |
| **M4** | 0.99 | 0.06 | 0.16 | 0.33 | 0.00 | 0.90 | 0.99 | 0.95 | 0.33 |
| **M5** | 0.13 | 0.89 | 0.33 | 0.38 | 0.90 | 0.00 | 0.94 | 0.73 | 0.86 |
| **M6** | 0.05 | 0.99 | 0.01 | 0.03 | 0.99 | 0.94 | 0.00 | 0.00 | 0.02 |
| **M7** | 0.73 | 0.01 | 0.99 | 0.01 | 0.95 | 0.73 | 0.00 | 0.00 | 0.03 |
| **S** | 0.18 | 0.62 | 0.00 | 0.01 | 0.33 | 0.86 | 0.02 | 0.03 | 0.00 |

Mutual affinity of the participants. Concentration of the duplex C_AB_ (μM) when oligos are mixed at 1 μM each (50 °C).

|  | **I** | **M1** | **M2** | **M3** | **M4** | **M5** | **M6** | **M7** | **S** |
| --- | --- | --- | --- | --- | --- | --- | --- | --- | --- |
| **I** | 0.00 | 0.82 | 0.01 | 0.02 | 0.82 | 0.01 | 0.00 | 0.10 | 0.01 |
| **M1** | 0.82 | 0.00 | 0.12 | 0.03 | 0.01 | 0.25 | 0.81 | 0.00 | 0.06 |
| **M2** | 0.01 | 0.12 | 0.00 | 0.11 | 0.01 | 0.02 | 0.00 | 0.70 | 0.00 |
| **M3** | 0.02 | 0.03 | 0.11 | 0.00 | 0.02 | 0.03 | 0.00 | 0.00 | 0.00 |
| **M4** | 0.82 | 0.01 | 0.01 | 0.02 | 0.00 | 0.28 | 0.82 | 0.45 | 0.02 |
| **M5** | 0.01 | 0.25 | 0.02 | 0.03 | 0.28 | 0.00 | 0.39 | 0.10 | 0.21 |
| **M6** | 0.00 | 0.81 | 0.00 | 0.00 | 0.82 | 0.39 | 0.00 | 0.00 | 0.00 |
| **M7** | 0.10 | 0.00 | 0.70 | 0.00 | 0.45 | 0.10 | 0.00 | 0.00 | 0.00 |
| **S** | 0.01 | 0.06 | 0.00 | 0.00 | 0.02 | 0.21 | 0.00 | 0.00 | 0.00 |

Mutual affinity of the participants. Approximate dissociation constant K_d_ (25 °C) in M.

|  | **I** | **M1** | **M2** | **M3** | **M4** | **M5** | **M6** | **M7** | **S** |
| --- | --- | --- | --- | --- | --- | --- | --- | --- | --- |
| **I** | 1.0E+00 | 3.0E-11 | 1.2E-05 | 2.8E-06 | 3.0E-11 | 6.0E-06 | 2.0E-05 | 9.8E-08 | 3.6E-06 |
| **M1** | 3.0E-11 | 1.0E+00 | 7.2E-08 | 1.1E-06 | 1.5E-05 | 1.4E-08 | 3.5E-11 | 1.1E-04 | 2.4E-07 |
| **M2** | 1.2E-05 | 7.2E-08 | 1.0E+00 | 8.4E-08 | 4.3E-06 | 1.4E-06 | 1.1E-04 | 1.9E-10 | 2.4E-04 |
| **M3** | 2.8E-06 | 1.1E-06 | 8.4E-08 | 1.0E+00 | 1.4E-06 | 1.0E-06 | 3.7E-05 | 8.1E-05 | 1.9E-04 |
| **M4** | 3.0E-11 | 1.5E-05 | 4.3E-06 | 1.4E-06 | 1.0E+00 | 1.1E-08 | 3.1E-11 | 2.2E-09 | 1.4E-06 |
| **M5** | 6.0E-06 | 1.4E-08 | 1.4E-06 | 1.0E-06 | 1.1E-08 | 1.0E+00 | 3.6E-09 | 1.0E-07 | 2.2E-08 |
| **M6** | 2.0E-05 | 3.5E-11 | 1.1E-04 | 3.7E-05 | 3.1E-11 | 3.6E-09 | 1.0E+00 | 2.9E-04 | 4.4E-05 |
| **M7** | 9.8E-08 | 1.1E-04 | 1.9E-10 | 8.1E-05 | 2.2E-09 | 1.0E-07 | 2.9E-04 | 1.0E+00 | 3.7E-05 |
| **S** | 3.6E-06 | 2.4E-07 | 2.4E-04 | 1.9E-04 | 1.4E-06 | 2.2E-08 | 4.4E-05 | 3.7E-05 | 1.0E+00 |

Mutual affinity of the participants. Approximate dissociation constant K_d_ (50 °C) in M.

|  | **I** | **M1** | **M2** | **M3** | **M4** | **M5** | **M6** | **M7** | **S** |
| --- | --- | --- | --- | --- | --- | --- | --- | --- | --- |
| **I** | 1.0E+00 | 3.9E-08 | 1.6E-04 | 6.3E-05 | 3.9E-08 | 1.0E-04 | 2.1E-04 | 7.8E-06 | 7.5E-05 |
| **M1** | 3.9E-08 | 1.0E+00 | 6.4E-06 | 3.5E-05 | 1.8E-04 | 2.2E-06 | 4.3E-08 | 6.0E-04 | 1.4E-05 |
| **M2** | 1.6E-04 | 6.4E-06 | 1.0E+00 | 7.1E-06 | 8.3E-05 | 4.1E-05 | 5.8E-04 | 1.3E-07 | 9.3E-04 |
| **M3** | 6.3E-05 | 3.5E-05 | 7.1E-06 | 1.0E+00 | 4.1E-05 | 3.4E-05 | 3.1E-04 | 4.9E-04 | 8.3E-04 |
| **M4** | 3.9E-08 | 1.8E-04 | 8.3E-05 | 4.1E-05 | 1.0E+00 | 1.9E-06 | 3.9E-08 | 6.7E-07 | 4.1E-05 |
| **M5** | 1.0E-04 | 2.2E-06 | 4.1E-05 | 3.4E-05 | 1.9E-06 | 1.0E+00 | 9.3E-07 | 8.0E-06 | 3.0E-06 |
| **M6** | 2.1E-04 | 4.3E-08 | 5.8E-04 | 3.1E-04 | 3.9E-08 | 9.3E-07 | 1.0E+00 | 1.0E-03 | 3.4E-04 |
| **M7** | 7.8E-06 | 6.0E-04 | 1.3E-07 | 4.9E-04 | 6.7E-07 | 8.0E-06 | 1.0E-03 | 1.0E+00 | 3.1E-04 |
| **S** | 7.5E-05 | 1.4E-05 | 9.3E-04 | 8.3E-04 | 4.1E-05 | 3.0E-06 | 3.4E-04 | 3.1E-04 | 1.0E+00 |

#### Cubic function – temperature-independent system

Concentrations of the participants.

| **ID** | **Concentration, μM** |
| --- | --- |
| I | 8.68 |
| M1 | 0.66 |
| M2 | 10.00 |
| M3 | 4.45 |
| M4 | 3.53 |
| M5 | 4.33 |
| M6 | 7.02 |
| M7 | 9.55 |
| S | 0.03 |
| C1 | 1.42 |
| C2 | 10.00 |
| C3 | 9.21 |
| C4 | 4.20 |
| C5 | 4.91 |
| C6 | 1.57 |
| C7 | 3.06 |
| C8 | 2.50 |

Mutual affinity of the participants. Concentration of the duplex C_AB_ (μM) when oligos are mixed at 1 μM each (25 °C).

|  | **I** | **M1** | **M2** | **M3** | **M4** | **M5** | **M6** | **M7** | **S** | **C1** | **C2** | **C3** | **C4** | **C5** | **C6** | **C7** | **C8** |
| --- | --- | --- | --- | --- | --- | --- | --- | --- | --- | --- | --- | --- | --- | --- | --- | --- | --- |
| **I** | 0.00 | 0.97 | 0.09 | 0.00 | 0.99 | 0.99 | 0.00 | 0.49 | 0.00 | 0.97 | 0.00 | 0.00 | 0.00 | 0.00 | 0.00 | 0.00 | 0.00 |
| **M1** | 0.97 | 0.00 | 0.01 | 0.98 | 0.99 | 0.57 | 0.99 | 0.35 | 0.99 | 0.00 | 0.99 | 0.00 | 0.00 | 0.00 | 0.00 | 0.00 | 0.00 |
| **M2** | 0.09 | 0.01 | 0.00 | 0.58 | 0.75 | 0.00 | 0.97 | 0.01 | 0.07 | 0.00 | 0.00 | 0.98 | 0.00 | 0.00 | 0.00 | 0.00 | 0.00 |
| **M3** | 0.00 | 0.98 | 0.58 | 0.00 | 0.30 | 0.33 | 0.24 | 0.99 | 0.41 | 0.00 | 0.00 | 0.00 | 0.00 | 0.00 | 0.00 | 0.00 | 0.00 |
| **M4** | 0.99 | 0.99 | 0.75 | 0.30 | 0.00 | 0.27 | 0.00 | 0.96 | 0.25 | 0.00 | 0.00 | 0.00 | 0.97 | 0.00 | 0.00 | 0.00 | 0.00 |
| **M5** | 0.99 | 0.57 | 0.00 | 0.33 | 0.27 | 0.00 | 0.02 | 0.07 | 0.57 | 0.00 | 0.00 | 0.00 | 0.00 | 0.95 | 0.00 | 0.00 | 0.00 |
| **M6** | 0.00 | 0.99 | 0.97 | 0.24 | 0.00 | 0.02 | 0.00 | 0.75 | 0.04 | 0.00 | 0.00 | 0.00 | 0.00 | 0.00 | 0.99 | 0.00 | 0.00 |
| **M7** | 0.49 | 0.35 | 0.01 | 0.99 | 0.96 | 0.07 | 0.75 | 0.00 | 0.47 | 0.00 | 0.00 | 0.00 | 0.00 | 0.00 | 0.00 | 0.93 | 0.00 |
| **S** | 0.00 | 0.99 | 0.07 | 0.41 | 0.25 | 0.57 | 0.04 | 0.47 | 0.00 | 0.00 | 0.00 | 0.00 | 0.00 | 0.00 | 0.00 | 0.00 | 0.01 |
| **C1** | 0.97 | 0.00 | 0.00 | 0.00 | 0.00 | 0.00 | 0.00 | 0.00 | 0.00 | 0.00 | 0.00 | 0.00 | 0.00 | 0.00 | 0.00 | 0.00 | 0.00 |
| **C2** | 0.00 | 0.99 | 0.00 | 0.00 | 0.00 | 0.00 | 0.00 | 0.00 | 0.00 | 0.00 | 0.00 | 0.00 | 0.00 | 0.00 | 0.00 | 0.00 | 0.00 |
| **C3** | 0.00 | 0.00 | 0.98 | 0.00 | 0.00 | 0.00 | 0.00 | 0.00 | 0.00 | 0.00 | 0.00 | 0.00 | 0.00 | 0.00 | 0.00 | 0.00 | 0.00 |
| **C4** | 0.00 | 0.00 | 0.00 | 0.00 | 0.97 | 0.00 | 0.00 | 0.00 | 0.00 | 0.00 | 0.00 | 0.00 | 0.00 | 0.00 | 0.00 | 0.00 | 0.00 |
| **C5** | 0.00 | 0.00 | 0.00 | 0.00 | 0.00 | 0.95 | 0.00 | 0.00 | 0.00 | 0.00 | 0.00 | 0.00 | 0.00 | 0.00 | 0.00 | 0.00 | 0.00 |
| **C6** | 0.00 | 0.00 | 0.00 | 0.00 | 0.00 | 0.00 | 0.99 | 0.00 | 0.00 | 0.00 | 0.00 | 0.00 | 0.00 | 0.00 | 0.00 | 0.00 | 0.00 |
| **C7** | 0.00 | 0.00 | 0.00 | 0.00 | 0.00 | 0.00 | 0.00 | 0.93 | 0.00 | 0.00 | 0.00 | 0.00 | 0.00 | 0.00 | 0.00 | 0.00 | 0.00 |
| **C8** | 0.00 | 0.00 | 0.00 | 0.00 | 0.00 | 0.00 | 0.00 | 0.00 | 0.01 | 0.00 | 0.00 | 0.00 | 0.00 | 0.00 | 0.00 | 0.00 | 0.00 |

Mutual affinity of the participants. Concentration of the duplex C_AB_ (μM) when oligos are mixed at 1 μM each (50 °C).

|  | **I** | **M1** | **M2** | **M3** | **M4** | **M5** | **M6** | **M7** | **S** | **C1** | **C2** | **C3** | **C4** | **C5** | **C6** | **C7** | **C8** |
| --- | --- | --- | --- | --- | --- | --- | --- | --- | --- | --- | --- | --- | --- | --- | --- | --- | --- |
| **I** | 0.00 | 0.58 | 0.01 | 0.00 | 0.75 | 0.82 | 0.00 | 0.04 | 0.00 | 0.05 | 0.00 | 0.00 | 0.00 | 0.00 | 0.00 | 0.00 | 0.00 |
| **M1** | 0.58 | 0.00 | 0.00 | 0.66 | 0.77 | 0.05 | 0.77 | 0.03 | 0.82 | 0.00 | 0.13 | 0.00 | 0.00 | 0.00 | 0.00 | 0.00 | 0.00 |
| **M2** | 0.01 | 0.00 | 0.00 | 0.06 | 0.11 | 0.00 | 0.52 | 0.00 | 0.01 | 0.00 | 0.00 | 0.08 | 0.00 | 0.00 | 0.00 | 0.00 | 0.00 |
| **M3** | 0.00 | 0.66 | 0.06 | 0.00 | 0.02 | 0.02 | 0.02 | 0.80 | 0.03 | 0.00 | 0.00 | 0.00 | 0.00 | 0.00 | 0.00 | 0.00 | 0.00 |
| **M4** | 0.75 | 0.77 | 0.11 | 0.02 | 0.00 | 0.02 | 0.00 | 0.47 | 0.02 | 0.00 | 0.00 | 0.00 | 0.05 | 0.00 | 0.00 | 0.00 | 0.00 |
| **M5** | 0.82 | 0.05 | 0.00 | 0.02 | 0.02 | 0.00 | 0.00 | 0.01 | 0.05 | 0.00 | 0.00 | 0.00 | 0.00 | 0.02 | 0.00 | 0.00 | 0.00 |
| **M6** | 0.00 | 0.77 | 0.52 | 0.02 | 0.00 | 0.00 | 0.00 | 0.11 | 0.00 | 0.00 | 0.00 | 0.00 | 0.00 | 0.00 | 0.22 | 0.00 | 0.00 |
| **M7** | 0.04 | 0.03 | 0.00 | 0.80 | 0.47 | 0.01 | 0.11 | 0.00 | 0.04 | 0.00 | 0.00 | 0.00 | 0.00 | 0.00 | 0.00 | 0.01 | 0.00 |
| **S** | 0.00 | 0.82 | 0.01 | 0.03 | 0.02 | 0.05 | 0.00 | 0.04 | 0.00 | 0.00 | 0.00 | 0.00 | 0.00 | 0.00 | 0.00 | 0.00 | 0.00 |
| **C1** | 0.05 | 0.00 | 0.00 | 0.00 | 0.00 | 0.00 | 0.00 | 0.00 | 0.00 | 0.00 | 0.00 | 0.00 | 0.00 | 0.00 | 0.00 | 0.00 | 0.00 |
| **C2** | 0.00 | 0.13 | 0.00 | 0.00 | 0.00 | 0.00 | 0.00 | 0.00 | 0.00 | 0.00 | 0.00 | 0.00 | 0.00 | 0.00 | 0.00 | 0.00 | 0.00 |
| **C3** | 0.00 | 0.00 | 0.08 | 0.00 | 0.00 | 0.00 | 0.00 | 0.00 | 0.00 | 0.00 | 0.00 | 0.00 | 0.00 | 0.00 | 0.00 | 0.00 | 0.00 |
| **C4** | 0.00 | 0.00 | 0.00 | 0.00 | 0.05 | 0.00 | 0.00 | 0.00 | 0.00 | 0.00 | 0.00 | 0.00 | 0.00 | 0.00 | 0.00 | 0.00 | 0.00 |
| **C5** | 0.00 | 0.00 | 0.00 | 0.00 | 0.00 | 0.02 | 0.00 | 0.00 | 0.00 | 0.00 | 0.00 | 0.00 | 0.00 | 0.00 | 0.00 | 0.00 | 0.00 |
| **C6** | 0.00 | 0.00 | 0.00 | 0.00 | 0.00 | 0.00 | 0.22 | 0.00 | 0.00 | 0.00 | 0.00 | 0.00 | 0.00 | 0.00 | 0.00 | 0.00 | 0.00 |
| **C7** | 0.00 | 0.00 | 0.00 | 0.00 | 0.00 | 0.00 | 0.00 | 0.01 | 0.00 | 0.00 | 0.00 | 0.00 | 0.00 | 0.00 | 0.00 | 0.00 | 0.00 |
| **C8** | 0.00 | 0.00 | 0.00 | 0.00 | 0.00 | 0.00 | 0.00 | 0.00 | 0.00 | 0.00 | 0.00 | 0.00 | 0.00 | 0.00 | 0.00 | 0.00 | 0.00 |

Mutual affinity of the participants. Approximate dissociation constant K_d_ (25 °C) in M.

|  | **I** | **M1** | **M2** | **M3** | **M4** | **M5** | **M6** | **M7** | **S** | **C1** | **C2** | **C3** | **C4** | **C5** | **C6** | **C7** | **C8** |
| --- | --- | --- | --- | --- | --- | --- | --- | --- | --- | --- | --- | --- | --- | --- | --- | --- | --- |
| **I** | 1.0E+00 | 6.5E-10 | 9.5E-06 | 2.5E-04 | 9.7E-11 | 3.1E-11 | 2.9E-04 | 5.4E-07 | 3.0E-04 | 7.6E-10 | 1.0E+00 | 1.0E+00 | 1.0E+00 | 1.0E+00 | 1.0E+00 | 1.0E+00 | 1.0E+00 |
| **M1** | 6.5E-10 | 1.0E+00 | 1.5E-04 | 2.9E-10 | 7.2E-11 | 3.3E-07 | 7.1E-11 | 1.2E-06 | 3.1E-11 | 1.0E+00 | 1.1E-10 | 1.0E+00 | 1.0E+00 | 1.0E+00 | 1.0E+00 | 1.0E+00 | 1.0E+00 |
| **M2** | 9.5E-06 | 1.5E-04 | 1.0E+00 | 3.0E-07 | 8.3E-08 | 2.1E-04 | 1.1E-09 | 7.5E-05 | 1.2E-05 | 1.0E+00 | 1.0E+00 | 2.6E-10 | 1.0E+00 | 1.0E+00 | 1.0E+00 | 1.0E+00 | 1.0E+00 |
| **M3** | 2.5E-04 | 2.9E-10 | 3.0E-07 | 1.0E+00 | 1.6E-06 | 1.3E-06 | 2.4E-06 | 4.1E-11 | 8.3E-07 | 1.0E+00 | 1.0E+00 | 1.0E+00 | 1.0E+00 | 1.0E+00 | 1.0E+00 | 1.0E+00 | 1.0E+00 |
| **M4** | 9.7E-11 | 7.2E-11 | 8.3E-08 | 1.6E-06 | 1.0E+00 | 2.0E-06 | 2.5E-04 | 1.8E-09 | 2.2E-06 | 1.0E+00 | 1.0E+00 | 1.0E+00 | 6.5E-10 | 1.0E+00 | 1.0E+00 | 1.0E+00 | 1.0E+00 |
| **M5** | 3.1E-11 | 3.3E-07 | 2.1E-04 | 1.3E-06 | 2.0E-06 | 1.0E+00 | 5.5E-05 | 1.2E-05 | 3.2E-07 | 1.0E+00 | 1.0E+00 | 1.0E+00 | 1.0E+00 | 2.3E-09 | 1.0E+00 | 1.0E+00 | 1.0E+00 |
| **M6** | 2.9E-04 | 7.1E-11 | 1.1E-09 | 2.4E-06 | 2.5E-04 | 5.5E-05 | 1.0E+00 | 8.6E-08 | 2.2E-05 | 1.0E+00 | 1.0E+00 | 1.0E+00 | 1.0E+00 | 1.0E+00 | 3.7E-11 | 1.0E+00 | 1.0E+00 |
| **M7** | 5.4E-07 | 1.2E-06 | 7.5E-05 | 4.1E-11 | 1.8E-09 | 1.2E-05 | 8.6E-08 | 1.0E+00 | 5.8E-07 | 1.0E+00 | 1.0E+00 | 1.0E+00 | 1.0E+00 | 1.0E+00 | 1.0E+00 | 5.4E-09 | 1.0E+00 |
| **S** | 3.0E-04 | 3.1E-11 | 1.2E-05 | 8.3E-07 | 2.2E-06 | 3.2E-07 | 2.2E-05 | 5.8E-07 | 1.0E+00 | 1.0E+00 | 1.0E+00 | 1.0E+00 | 1.0E+00 | 1.0E+00 | 1.0E+00 | 1.0E+00 | 1.5E-04 |
| **C1** | 7.6E-10 | 1.0E+00 | 1.0E+00 | 1.0E+00 | 1.0E+00 | 1.0E+00 | 1.0E+00 | 1.0E+00 | 1.0E+00 | 1.0E+00 | 1.0E+00 | 1.0E+00 | 1.0E+00 | 1.0E+00 | 1.0E+00 | 1.0E+00 | 1.0E+00 |
| **C2** | 1.0E+00 | 1.1E-10 | 1.0E+00 | 1.0E+00 | 1.0E+00 | 1.0E+00 | 1.0E+00 | 1.0E+00 | 1.0E+00 | 1.0E+00 | 1.0E+00 | 1.0E+00 | 1.0E+00 | 1.0E+00 | 1.0E+00 | 1.0E+00 | 1.0E+00 |
| **C3** | 1.0E+00 | 1.0E+00 | 2.6E-10 | 1.0E+00 | 1.0E+00 | 1.0E+00 | 1.0E+00 | 1.0E+00 | 1.0E+00 | 1.0E+00 | 1.0E+00 | 1.0E+00 | 1.0E+00 | 1.0E+00 | 1.0E+00 | 1.0E+00 | 1.0E+00 |
| **C4** | 1.0E+00 | 1.0E+00 | 1.0E+00 | 1.0E+00 | 6.5E-10 | 1.0E+00 | 1.0E+00 | 1.0E+00 | 1.0E+00 | 1.0E+00 | 1.0E+00 | 1.0E+00 | 1.0E+00 | 1.0E+00 | 1.0E+00 | 1.0E+00 | 1.0E+00 |
| **C5** | 1.0E+00 | 1.0E+00 | 1.0E+00 | 1.0E+00 | 1.0E+00 | 2.3E-09 | 1.0E+00 | 1.0E+00 | 1.0E+00 | 1.0E+00 | 1.0E+00 | 1.0E+00 | 1.0E+00 | 1.0E+00 | 1.0E+00 | 1.0E+00 | 1.0E+00 |
| **C6** | 1.0E+00 | 1.0E+00 | 1.0E+00 | 1.0E+00 | 1.0E+00 | 1.0E+00 | 3.7E-11 | 1.0E+00 | 1.0E+00 | 1.0E+00 | 1.0E+00 | 1.0E+00 | 1.0E+00 | 1.0E+00 | 1.0E+00 | 1.0E+00 | 1.0E+00 |
| **C7** | 1.0E+00 | 1.0E+00 | 1.0E+00 | 1.0E+00 | 1.0E+00 | 1.0E+00 | 1.0E+00 | 5.4E-09 | 1.0E+00 | 1.0E+00 | 1.0E+00 | 1.0E+00 | 1.0E+00 | 1.0E+00 | 1.0E+00 | 1.0E+00 | 1.0E+00 |
| **C8** | 1.0E+00 | 1.0E+00 | 1.0E+00 | 1.0E+00 | 1.0E+00 | 1.0E+00 | 1.0E+00 | 1.0E+00 | 1.5E-04 | 1.0E+00 | 1.0E+00 | 1.0E+00 | 1.0E+00 | 1.0E+00 | 1.0E+00 | 1.0E+00 | 1.0E+00 |

Mutual affinity of the participants. Approximate dissociation constant K_d_ (50 °C) in M.

|  | **I** | **M1** | **M2** | **M3** | **M4** | **M5** | **M6** | **M7** | **S** | **C1** | **C2** | **C3** | **C4** | **C5** | **C6** | **C7** | **C8** |
| --- | --- | --- | --- | --- | --- | --- | --- | --- | --- | --- | --- | --- | --- | --- | --- | --- | --- |
| **I** | 1.0E+00 | 3.0E-07 | 1.3E-04 | 9.7E-04 | 8.6E-08 | 4.0E-08 | 1.1E-03 | 2.3E-05 | 1.1E-03 | 2.0E-05 | 1.0E+00 | 1.0E+00 | 1.0E+00 | 1.0E+00 | 1.0E+00 | 1.0E+00 | 1.0E+00 |
| **M1** | 3.0E-07 | 1.0E+00 | 7.2E-04 | 1.8E-07 | 7.0E-08 | 1.7E-05 | 6.9E-08 | 3.7E-05 | 3.9E-08 | 1.0E+00 | 5.7E-06 | 1.0E+00 | 1.0E+00 | 1.0E+00 | 1.0E+00 | 1.0E+00 | 1.0E+00 |
| **M2** | 1.3E-04 | 7.2E-04 | 1.0E+00 | 1.6E-05 | 7.0E-06 | 8.7E-04 | 4.4E-07 | 4.7E-04 | 1.6E-04 | 1.0E+00 | 1.0E+00 | 1.0E-05 | 1.0E+00 | 1.0E+00 | 1.0E+00 | 1.0E+00 | 1.0E+00 |
| **M3** | 9.7E-04 | 1.8E-07 | 1.6E-05 | 1.0E+00 | 4.5E-05 | 4.0E-05 | 5.8E-05 | 4.8E-08 | 3.0E-05 | 1.0E+00 | 1.0E+00 | 1.0E+00 | 1.0E+00 | 1.0E+00 | 1.0E+00 | 1.0E+00 | 1.0E+00 |
| **M4** | 8.6E-08 | 7.0E-08 | 7.0E-06 | 4.5E-05 | 1.0E+00 | 5.2E-05 | 9.6E-04 | 5.9E-07 | 5.6E-05 | 1.0E+00 | 1.0E+00 | 1.0E+00 | 1.8E-05 | 1.0E+00 | 1.0E+00 | 1.0E+00 | 1.0E+00 |
| **M5** | 4.0E-08 | 1.7E-05 | 8.7E-04 | 4.0E-05 | 5.2E-05 | 1.0E+00 | 3.9E-04 | 1.6E-04 | 1.6E-05 | 1.0E+00 | 1.0E+00 | 1.0E+00 | 1.0E+00 | 4.0E-05 | 1.0E+00 | 1.0E+00 | 1.0E+00 |
| **M6** | 1.1E-03 | 6.9E-08 | 4.4E-07 | 5.8E-05 | 9.6E-04 | 3.9E-04 | 1.0E+00 | 7.2E-06 | 2.3E-04 | 1.0E+00 | 1.0E+00 | 1.0E+00 | 1.0E+00 | 1.0E+00 | 2.7E-06 | 1.0E+00 | 1.0E+00 |
| **M7** | 2.3E-05 | 3.7E-05 | 4.7E-04 | 4.8E-08 | 5.9E-07 | 1.6E-04 | 7.2E-06 | 1.0E+00 | 2.4E-05 | 1.0E+00 | 1.0E+00 | 1.0E+00 | 1.0E+00 | 1.0E+00 | 1.0E+00 | 6.6E-05 | 1.0E+00 |
| **S** | 1.1E-03 | 3.9E-08 | 1.6E-04 | 3.0E-05 | 5.6E-05 | 1.6E-05 | 2.3E-04 | 2.4E-05 | 1.0E+00 | 1.0E+00 | 1.0E+00 | 1.0E+00 | 1.0E+00 | 1.0E+00 | 1.0E+00 | 1.0E+00 | 4.6E-03 |
| **C1** | 2.0E-05 | 1.0E+00 | 1.0E+00 | 1.0E+00 | 1.0E+00 | 1.0E+00 | 1.0E+00 | 1.0E+00 | 1.0E+00 | 1.0E+00 | 1.0E+00 | 1.0E+00 | 1.0E+00 | 1.0E+00 | 1.0E+00 | 1.0E+00 | 1.0E+00 |
| **C2** | 1.0E+00 | 5.7E-06 | 1.0E+00 | 1.0E+00 | 1.0E+00 | 1.0E+00 | 1.0E+00 | 1.0E+00 | 1.0E+00 | 1.0E+00 | 1.0E+00 | 1.0E+00 | 1.0E+00 | 1.0E+00 | 1.0E+00 | 1.0E+00 | 1.0E+00 |
| **C3** | 1.0E+00 | 1.0E+00 | 1.0E-05 | 1.0E+00 | 1.0E+00 | 1.0E+00 | 1.0E+00 | 1.0E+00 | 1.0E+00 | 1.0E+00 | 1.0E+00 | 1.0E+00 | 1.0E+00 | 1.0E+00 | 1.0E+00 | 1.0E+00 | 1.0E+00 |
| **C4** | 1.0E+00 | 1.0E+00 | 1.0E+00 | 1.0E+00 | 1.8E-05 | 1.0E+00 | 1.0E+00 | 1.0E+00 | 1.0E+00 | 1.0E+00 | 1.0E+00 | 1.0E+00 | 1.0E+00 | 1.0E+00 | 1.0E+00 | 1.0E+00 | 1.0E+00 |
| **C5** | 1.0E+00 | 1.0E+00 | 1.0E+00 | 1.0E+00 | 1.0E+00 | 4.0E-05 | 1.0E+00 | 1.0E+00 | 1.0E+00 | 1.0E+00 | 1.0E+00 | 1.0E+00 | 1.0E+00 | 1.0E+00 | 1.0E+00 | 1.0E+00 | 1.0E+00 |
| **C6** | 1.0E+00 | 1.0E+00 | 1.0E+00 | 1.0E+00 | 1.0E+00 | 1.0E+00 | 2.7E-06 | 1.0E+00 | 1.0E+00 | 1.0E+00 | 1.0E+00 | 1.0E+00 | 1.0E+00 | 1.0E+00 | 1.0E+00 | 1.0E+00 | 1.0E+00 |
| **C7** | 1.0E+00 | 1.0E+00 | 1.0E+00 | 1.0E+00 | 1.0E+00 | 1.0E+00 | 1.0E+00 | 6.6E-05 | 1.0E+00 | 1.0E+00 | 1.0E+00 | 1.0E+00 | 1.0E+00 | 1.0E+00 | 1.0E+00 | 1.0E+00 | 1.0E+00 |
| **C8** | 1.0E+00 | 1.0E+00 | 1.0E+00 | 1.0E+00 | 1.0E+00 | 1.0E+00 | 1.0E+00 | 1.0E+00 | 4.6E-03 | 1.0E+00 | 1.0E+00 | 1.0E+00 | 1.0E+00 | 1.0E+00 | 1.0E+00 | 1.0E+00 | 1.0E+00 |

#### Cubic function – control system without compensatory participants

Concentrations of the participants.

| **ID** | **Concentration, μM** |
| --- | --- |
| I | 8.68 |
| M1 | 0.73 |
| M2 | 3.75 |
| M3 | 4.66 |
| M4 | 2.71 |
| M5 | 2.58 |
| M6 | 5.15 |
| M7 | 9.79 |
| S | 0.03 |

Mutual affinity of the participants. Concentration of the duplex C_AB_ (μM) when oligos are mixed at 1 μM each (25 °C).

|  | **I** | **M1** | **M2** | **M3** | **M4** | **M5** | **M6** | **M7** | **S** |
| --- | --- | --- | --- | --- | --- | --- | --- | --- | --- |
| **I** | 0.00 | 0.98 | 0.00 | 0.33 | 0.99 | 0.99 | 0.98 | 0.64 | 0.10 |
| **M1** | 0.98 | 0.00 | 0.00 | 0.99 | 0.99 | 0.03 | 0.99 | 0.69 | 0.99 |
| **M2** | 0.00 | 0.00 | 0.00 | 0.13 | 0.99 | 0.96 | 0.96 | 0.57 | 0.19 |
| **M3** | 0.33 | 0.99 | 0.13 | 0.00 | 0.11 | 0.27 | 0.00 | 0.99 | 0.90 |
| **M4** | 0.99 | 0.99 | 0.99 | 0.11 | 0.00 | 0.12 | 0.00 | 0.93 | 0.86 |
| **M5** | 0.99 | 0.03 | 0.96 | 0.27 | 0.12 | 0.00 | 0.16 | 0.93 | 0.80 |
| **M6** | 0.98 | 0.99 | 0.96 | 0.00 | 0.00 | 0.16 | 0.00 | 0.79 | 0.53 |
| **M7** | 0.64 | 0.69 | 0.57 | 0.99 | 0.93 | 0.93 | 0.79 | 0.00 | 0.69 |
| **S** | 0.10 | 0.99 | 0.19 | 0.90 | 0.86 | 0.80 | 0.53 | 0.69 | 0.00 |

Mutual affinity of the participants. Concentration of the duplex C_AB_ (μM) when oligos are mixed at 1 μM each (50 °C).

|  | **I** | **M1** | **M2** | **M3** | **M4** | **M5** | **M6** | **M7** | **S** |
| --- | --- | --- | --- | --- | --- | --- | --- | --- | --- |
| **I** | 0.00 | 0.66 | 0.00 | 0.02 | 0.82 | 0.76 | 0.62 | 0.07 | 0.01 |
| **M1** | 0.66 | 0.00 | 0.00 | 0.71 | 0.75 | 0.00 | 0.76 | 0.09 | 0.72 |
| **M2** | 0.00 | 0.00 | 0.00 | 0.01 | 0.70 | 0.49 | 0.49 | 0.05 | 0.01 |
| **M3** | 0.02 | 0.71 | 0.01 | 0.00 | 0.01 | 0.02 | 0.00 | 0.82 | 0.28 |
| **M4** | 0.82 | 0.75 | 0.70 | 0.01 | 0.00 | 0.01 | 0.00 | 0.36 | 0.21 |
| **M5** | 0.76 | 0.00 | 0.49 | 0.02 | 0.01 | 0.00 | 0.01 | 0.34 | 0.14 |
| **M6** | 0.62 | 0.76 | 0.49 | 0.00 | 0.00 | 0.01 | 0.00 | 0.14 | 0.05 |
| **M7** | 0.07 | 0.09 | 0.05 | 0.82 | 0.36 | 0.34 | 0.14 | 0.00 | 0.08 |
| **S** | 0.01 | 0.72 | 0.01 | 0.28 | 0.21 | 0.14 | 0.05 | 0.08 | 0.00 |

Mutual affinity of the participants. Approximate dissociation constant K_d_ (25 °C) in M.

|  | **I** | **M1** | **M2** | **M3** | **M4** | **M5** | **M6** | **M7** | **S** |
| --- | --- | --- | --- | --- | --- | --- | --- | --- | --- |
| **I** | 1.0E+00 | 2.9E-10 | 2.9E-04 | 1.3E-06 | 3.0E-11 | 8.2E-11 | 4.2E-10 | 2.0E-07 | 8.1E-06 |
| **M1** | 2.9E-10 | 1.0E+00 | 2.4E-04 | 1.7E-10 | 8.9E-11 | 3.3E-05 | 7.6E-11 | 1.3E-07 | 1.5E-10 |
| **M2** | 2.9E-04 | 2.4E-04 | 1.0E+00 | 5.7E-06 | 1.8E-10 | 1.6E-09 | 1.5E-09 | 3.2E-07 | 3.5E-06 |
| **M3** | 1.3E-06 | 1.7E-10 | 5.7E-06 | 1.0E+00 | 7.5E-06 | 2.0E-06 | 2.8E-04 | 3.0E-11 | 1.1E-08 |
| **M4** | 3.0E-11 | 8.9E-11 | 1.8E-10 | 7.5E-06 | 1.0E+00 | 6.4E-06 | 2.9E-04 | 4.8E-09 | 2.2E-08 |
| **M5** | 8.2E-11 | 3.3E-05 | 1.6E-09 | 2.0E-06 | 6.4E-06 | 1.0E+00 | 4.2E-06 | 5.7E-09 | 5.0E-08 |
| **M6** | 4.2E-10 | 7.6E-11 | 1.5E-09 | 2.8E-04 | 2.9E-04 | 4.2E-06 | 1.0E+00 | 5.4E-08 | 4.1E-07 |
| **M7** | 2.0E-07 | 1.3E-07 | 3.2E-07 | 3.0E-11 | 4.8E-09 | 5.7E-09 | 5.4E-08 | 1.0E+00 | 1.4E-07 |
| **S** | 8.1E-06 | 1.5E-10 | 3.5E-06 | 1.1E-08 | 2.2E-08 | 5.0E-08 | 4.1E-07 | 1.4E-07 | 1.0E+00 |

Mutual affinity of the participants. Approximate dissociation constant K_d_ (50 °C) in M.

|  | **I** | **M1** | **M2** | **M3** | **M4** | **M5** | **M6** | **M7** | **S** |
| --- | --- | --- | --- | --- | --- | --- | --- | --- | --- |
| **I** | 1.0E+00 | 1.8E-07 | 1.1E-03 | 4.0E-05 | 3.9E-08 | 7.7E-08 | 2.3E-07 | 1.2E-05 | 1.2E-04 |
| **M1** | 1.8E-07 | 1.0E+00 | 9.4E-04 | 1.2E-07 | 8.1E-08 | 2.9E-04 | 7.3E-08 | 9.6E-06 | 1.1E-07 |
| **M2** | 1.1E-03 | 9.4E-04 | 1.0E+00 | 9.9E-05 | 1.3E-07 | 5.4E-07 | 5.2E-07 | 1.6E-05 | 7.3E-05 |
| **M3** | 4.0E-05 | 1.2E-07 | 9.9E-05 | 1.0E+00 | 1.2E-04 | 5.2E-05 | 1.0E-03 | 3.9E-08 | 1.9E-06 |
| **M4** | 3.9E-08 | 8.1E-08 | 1.3E-07 | 1.2E-04 | 1.0E+00 | 1.1E-04 | 1.1E-03 | 1.1E-06 | 3.0E-06 |
| **M5** | 7.7E-08 | 2.9E-04 | 5.4E-07 | 5.2E-05 | 1.1E-04 | 1.0E+00 | 8.2E-05 | 1.3E-06 | 5.1E-06 |
| **M6** | 2.3E-07 | 7.3E-08 | 5.2E-07 | 1.0E-03 | 1.1E-03 | 8.2E-05 | 1.0E+00 | 5.3E-06 | 1.9E-05 |
| **M7** | 1.2E-05 | 9.6E-06 | 1.6E-05 | 3.9E-08 | 1.1E-06 | 1.3E-06 | 5.3E-06 | 1.0E+00 | 9.9E-06 |
| **S** | 1.2E-04 | 1.1E-07 | 7.3E-05 | 1.9E-06 | 3.0E-06 | 5.1E-06 | 1.9E-05 | 9.9E-06 | 1.0E+00 |
